# Phosphorylation-dependent sub-functionalization of the calcium-dependent protein kinase CPK28

**DOI:** 10.1101/2020.10.16.338442

**Authors:** Melissa Bredow, Kyle W. Bender, Alexandra Johnson Dingee, Danalyn R. Holmes, Alysha Thomson, Danielle Ciren, Cailun A. S. Tanney, Katherine E. Dunning, Marco Trujillo, Steven C. Huber, Jacqueline Monaghan

## Abstract

Calcium (Ca^2+^)-dependent protein kinases (CDPKs or CPKs) are a unique family of Ca^2+^-sensor/kinase-effector proteins with diverse functions in plants. In *Arabidopsis thaliana*, CPK28 contributes to immune homeostasis by promoting degradation of the key immune signaling receptor-like cytoplasmic kinase BOTRYTIS-INDUCED KINASE 1 (BIK1), and additionally functions in vegetative-to-reproductive stage transition. How CPK28 controls these seemingly disparate pathways is unknown. Here, we identify a single phosphorylation site in the kinase domain of CPK28 (Ser318) that is differentially required for its function in immune homeostasis and stem elongation. We show that CPK28 undergoes intra- and inter-molecular auto-phosphorylation on Ser318 and can additionally be trans-phosphorylated on this residue by BIK1. Analysis of several other phosphorylation sites demonstrates that Ser318 phosphorylation is uniquely required to prime CPK28 for Ca^2+^ activation at physiological concentrations of Ca^2+^, possibly through stabilization of the Ca^2+^-bound active state as indicated by intrinsic fluorescence experiments. Together, our data indicate that phosphorylation of Ser318 is required for the activation of CPK28 at low intracellular [Ca^2+^] to prevent initiation of an immune response in the absence of infection. By comparison, phosphorylation of Ser318 is not required for stem-elongation, indicating pathway specific requirements for phosphorylation-based Ca^2+^-sensitivity priming. We additionally provide evidence for a conserved function for Ser318 phosphorylation in related group IV CDPKs which holds promise for biotechnological applications by generating CDPK alleles that enhance resistance to microbial pathogens without consequences to yield.

## INTRODUCTION

Protein kinases represent one of the largest eukaryotic protein superfamilies. While roughly 500 protein kinases have been identified in humans (1), the genomes of *Arabidopsis thaliana* (hereafter, Arabidopsis) (2) and *Oryza sativa* (3) encode more than 1000 and 1500 protein kinases, respectively, including several families unique to plants. Among these protein kinases are the receptor-like kinases (RLKs), receptor-like cytoplasmic kinases (RLCKs), and calcium-dependent protein kinases (CDPKs or CPKs) that have emerged as key regulators of plant immunity (4–6). Despite encompassing only 2% of most eukaryotic genomes, protein kinases phosphorylate more than 40% of cellular proteins (7, 8), reflecting their diverse roles in coordinating intracellular signaling events. Reversible phosphorylation of serine (Ser), threonine (Thr), and tyrosine (Tyr) residues can serve an array of functions including changes in protein conformation and activation state (9, 10), protein stability and degradation (11, 12), subcellular localization (13–15), and interaction with protein substrates (16–18).

Calcium (Ca^2+^) is a ubiquitous secondary messenger that acts cooperatively with protein phosphorylation to propagate intracellular signals. Spatial and temporal changes in intracellular Ca^2+^ levels occur in response to environmental and developmental cues (19–23). In plants, Ca^2+^ transients are decoded by four major groups of calcium sensor proteins (CSPs), which possess one or more Ca^2+^-binding EF-hand motifs (24, 25): calmodulins (CaM), CaM-like proteins (CMLs), calcineurin B-like proteins (CBLs), CDPKs, and Ca^2+^ and Ca^2+^/CaM-dependent protein kinases (CCaMKs).

At the intersection of phosphorylation cascades and Ca^2+^ signalling are CDPKs, a unique family of Ca^2+^-sensor/kinase-effector proteins. CDPKs have been identified in all land plants, green algae, as well as certain protozoan ciliates and apicomplexan parasites (26, 27). CDPKs have a conserved domain architecture, consisting of a canonical Ser/Thr protein kinase domain and an EF-hand containing Ca^2+^-binding CaM-like domain, linked together by an autoinhibitory junction (AIJ) and flanked by variable regions on both the amino (N) and carboxyl (C) termini (28, 29). As their name implies, most CDPKs require Ca^2+^ for their activation (30). Upon Ca^2+^ binding to all EF-hands in the CaM-like domain, a dramatic conformational change occurs, freeing the AIJ from the catalytic site of the kinase, rendering the enzyme active (31–33). CDPKs vary in their sensitivity to Ca^2+^ (30), presumably allowing proteins to perceive distinct stimuli through differences in Ca^2+^-binding affinity. For example, *Arabidopsis* CPK4 displays half maximal kinase activity in the presence of ∼ 3 μM of free Ca^2+^ (30) while CPK5 only requires ∼100 nM (34). Importantly, CDPKs are signaling hubs with documented roles in multiple distinct pathways (4, 24, 35–37) and are therefore likely regulated beyond Ca^2+^ activation.

Sub-functionationization is at least partially mediated by protein localization and interaction with pathway-specific binding partners, as is well-documented for Arabidopsis CPK3 which functions in response to biotic and abiotic stimuli in distinct cellular compartments (38). Recent attention has been drawn to site-specific phosphorylation as a way to regulate the activity of multifunctional kinases. For example, phosphorylation sites on the RLK BRASSINOSTEROID INSENSITIVE 1-ASSOCIATED KINASE 1 (BAK1) are differentially required for its function as a co-receptor with a subset of leucine-rich repeat (LRR)-RLKs (39). Phosphoproteomic analyses indicate that CDPKs are differentially phosphorylated following exposure to distinct stimuli (40–47); however, the biochemical mechanisms by which site-specific phosphorylation regulates multifunctional CDPKs is still poorly understood.

Arabidopsis CPK28 is a plasma membrane-localized protein kinase with dual roles in plant immune homeostasis (48–50) and phytohormone-mediated reproductive growth (51, 52). In vegetative plants, CPK28 serves as a negative regulator of immune signal amplitude by phosphorylating and activating two PLANT U-BOX type E3 ubiquitin ligases, PUB25 and PUB26, which target the key immune RLCK BOTRYTIS-INDUCED KINASE 1 (BIK1) for proteasomal degradation (49). Owing to elevated levels of BIK1, CPK28 null plants (*cpk28-1*) have heightened immune responses and enhanced resistance to the bacterial pathogen *Pseudomonas syringae* pv. *tomato* DC3000 (*Pto* DC3000) (50). Upon transition to the reproductive stage, *cpk28-1* plants additionally present shorter leaf petioles, enhanced anthocyanin production, and a reduction in stem elongation (51, 52). The molecular basis for developmental phenotypes in the *cpk28-1* knockout mutant, beyond hormonal imbalance (51, 52), are comparatively unknown.

Our recent work demonstrated that autophosphorylation status dictates Ca^2+^ sensitivity of CPK28 peptide kinase activity *in vitro* (53). While dephosphorylated CPK28 is stimulated by the addition of 100 μM CaCl₂ compared to untreated protein, hyperphosphorylated CPK28 displayed similar levels of activity at basal Ca^2+^ concentrations (53). These results highlight the interesting possibility that phosphorylation status may control the activation of multifunctional kinases in distinct pathways by allowing CDPKs to respond to stimuli-specific Ca^2+^ signatures.

In the present study, we identify a single autophosphorylation site, Ser318, that decouples the activity of CPK28 in immune signaling from its role in reproductive growth. We show that expression of a non-phosphorylatable Ser-to-Ala variant (CPK28^S318A^) is unable to complement the immune phenotypes of *cpk28-1* null mutants but is able to complement defects in stem growth. Further, we uncover a functional role for phosphorylation of Ser318 in priming CPK28 for activation at low free [Ca^2+^]. Together, we demonstrate that site-specific phosphorylation can direct the activity of a multifunctional kinase in distinct pathways and provide evidence for a conserved mechanism in orthologous group IV CDPKs.

## RESULTS

### Phosphorylation on Ser318 is required for CPK28-mediated immune homeostasis

CPK28 is phosphorylated on multiple sites *in vitro* and *in vivo* (51, 53–55). To determine whether the function of CPK28 in reproductive stage transition and immunity is bifurcated by site-specific phosphorylation, we generated phospho-ablative (Ser-to-Ala) mutations in three known *in vivo* auto-phosphorylation sites conserved in CPK28 orthologs across land plants (Fig S1): Ser228, Ser318, and Ser495 (51) (Fig 1A). When driven by the cauliflower mosaic virus (CaMV) *35S* promoter, CPK28^S228A^, CPK28^S318A^ and CPK28^S495A^ functionally complement the stem elongation phenotype observed in *cpk28-1* (51); however, it is unknown if these phosphosites regulate CPK28 function in immune homeostasis. To mitigate the possible effects of ectopic overexpression, we chose to stably express CPK28 mutants under the control of the native *pCPK28* promoter in the *cpk28-1* background and assessed functional complementation of defects in both stem elongation and immune signaling. Interestingly, we found that while all three mutations were able to complement the stem elongation phenotype of *cpk28-1* (Fig 1B), only CPK28^S228A^ and CPK28^S495A^ complemented the enhanced oxidative burst in *cpk28-1* following treatment with the endogenous immune elicitor peptide AtPep1 (Fig 1C). In addition, *cpk28-1/pCPK28:CPK28^S318A^-FLAG* lines did not complement *cpk28-1* in oxidative burst assays following treatment with the bacterial elicitor peptide elf18 (Fig S2), remained hyper-responsive to AtPep1 in seedling growth inhibition assays (Fig 1D), and more resistant than Col-0 to infection with the virulent bacterial pathogen *Pto* DC3000 (Fig 1E). Together, these results suggest that phosphorylation of Ser318 is uniquely required for CPK28 function in immune signaling.

**Figure 1.**
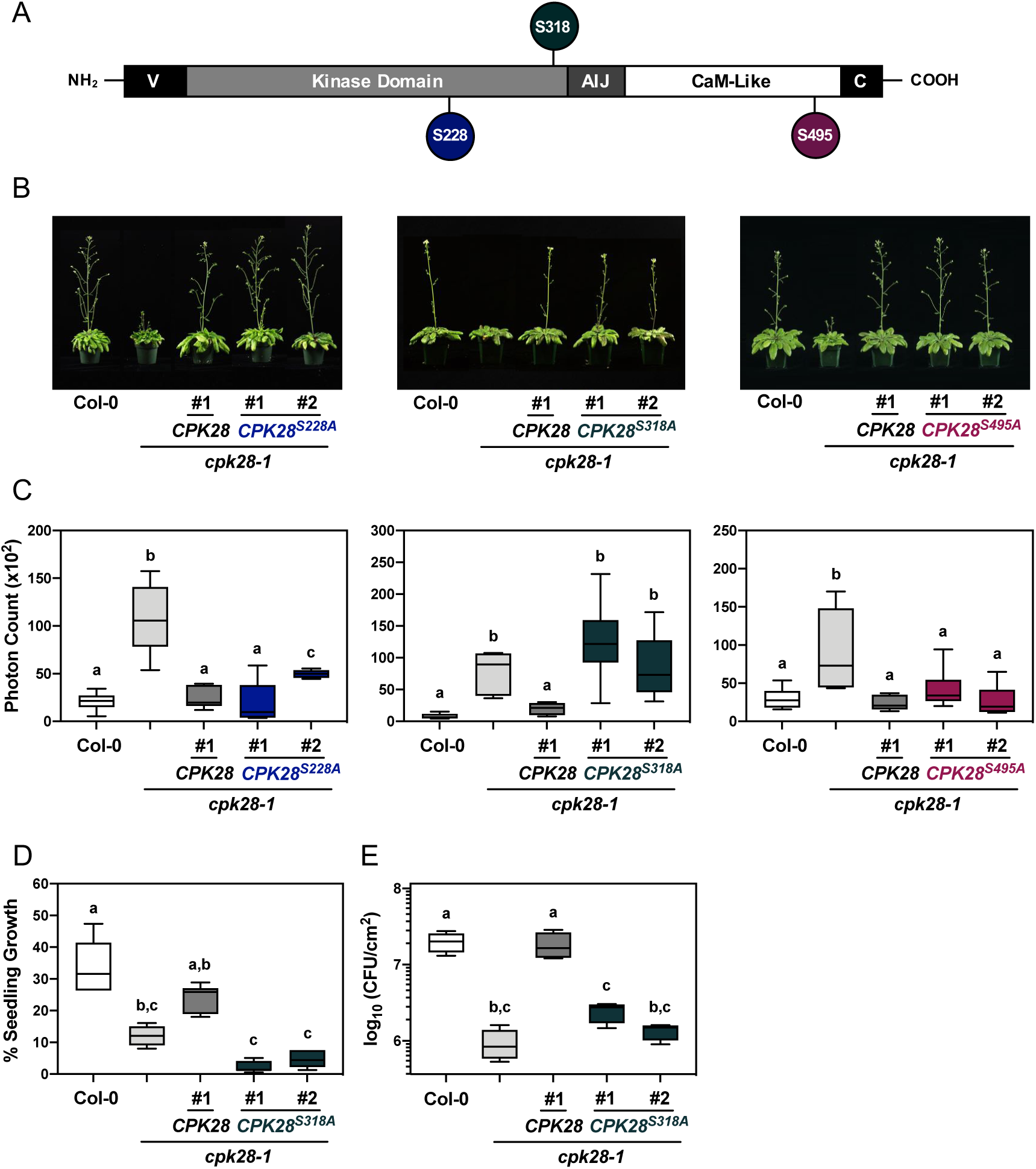
Ser318 phosphorylation differentially regulates CPK28 function in immune homeostasis. *(A)* Domain structure of CPK28 and position of tested autophosphorylation sites. V= amino-terminal variable domain; AIJ= autoinhibitory junction; C= carboxyl-terminal variable domain. *(B)* Stem elongation of six-week-old *Arabidopsis* plants and *(C)* AtPep1 (500 nM) triggered oxidative burst in 5-week-old plants (n=6) in the indicated genotypes. Data are presented as boxplots indicating first and third quartiles, split by a median line. Whiskers represent maximum and minimum values. *(D)* Seedling growth inhibition of Col-0, *cpk28-1*, *cpk28-1/pCPK28:CPK28-FLAG*, and *cpk28-1/pCPK28:CPK28^S318A^-FLAG* lines (n=6) resulting from continual growth in AtPep1 (500 nM) for 12 days. Values are normalized to untreated seedlings and presented as boxplots indicating first and third quartiles, split by a median line. Whiskers represent maximum and minimum values. *(E)* Growth of virulent *Pseudomonas syringae pv. tomato* (DC3000) in Col-0, *cpk28-1*, *cpk28-1/pCPK28:CPK28-FLAG*, and *cpk28-1/pCPK28:CPK28^S318A^-FLAG* lines (n=4). Samples were collected 3 days post infection and serially diluted. Values are presented as log transformed colony forming units (CFU) per cm^2^ and displayed as boxplots indicating first and third quartiles, split by a median line. Whiskers represent maximum and minimum values. At least three independent biological replicates were conducted for all experiments with similar results. Statistically different groups (p<0.005) are indicated with lowercase letters, as determined by ANOVA analysis followed by Tukey’s posthoc test.

Confocal imaging confirmed that CPK28^S318A^-YFP localizes to the plasma membrane in Arabidopsis stably expressing *35S:CPK28^S318A^-YFP* (Fig S3A), suggesting that phosphorylation of Ser318 does not affect the subcellular localization of CPK28. Catalytically inactive CPK28^D188A^-YFP was also observed at the plasma membrane (Fig S3A), indicating that CPK28 autophosphorylation is not required for appropriate localization. Furthermore, ablation of Ser318 phosphorylation did not compromise catalytic activity toward biological substrates PUB25 and PUB26 in *in vitro* kinase assays (Fig S3B), leading us to conclude that Ser318 phosphorylation does not regulate the function of CPK28 in immunity through altered subcellular localization or substrate specificity.

### CPK28-Ser318 undergoes intra- and inter-molecular autophosphorylation

Most protein kinases autophosphorylate *in vitro* (56). CPK28 peptides containing a phosphorylated Ser318 have been observed in mass spectra from several studies (51, 53–55). We validated that Ser318 is a Ca^2+^-dependent autophosphorylation site by conducting autophosphorylation assays with increasing levels of CaCl_2_ (Fig 2A) and probing with a phosphorylation and site-specific antibody raised against phosphorylated Ser318 (Fig S4). We also used the pIMAGO phospho-protein detection reagent to observe total autophosphorylation levels of CPK28. To determine if CPK28 autophosphorylates Ser318 *in cis* or *in trans*, we additionally conducted *in vitro* kinase assays using recombinantly produced MBP-His_6_-CPK28^D188A^ as a substrate for His_6_- CPK28 (Fig 2B). His_6_-CPK28 could trans-phosphorylate MBP-His_6_-CPK28^D188A^, indicating that Ser318 autophosphorylation can occur in both *cis* and *trans* (Fig 2B). Importantly, these results indicate that CPK28 autophosphorylation, including on Ser318, can occur at levels of free Ca^2+^ expected to occur under resting conditions *in vivo*.

**Figure 2.**
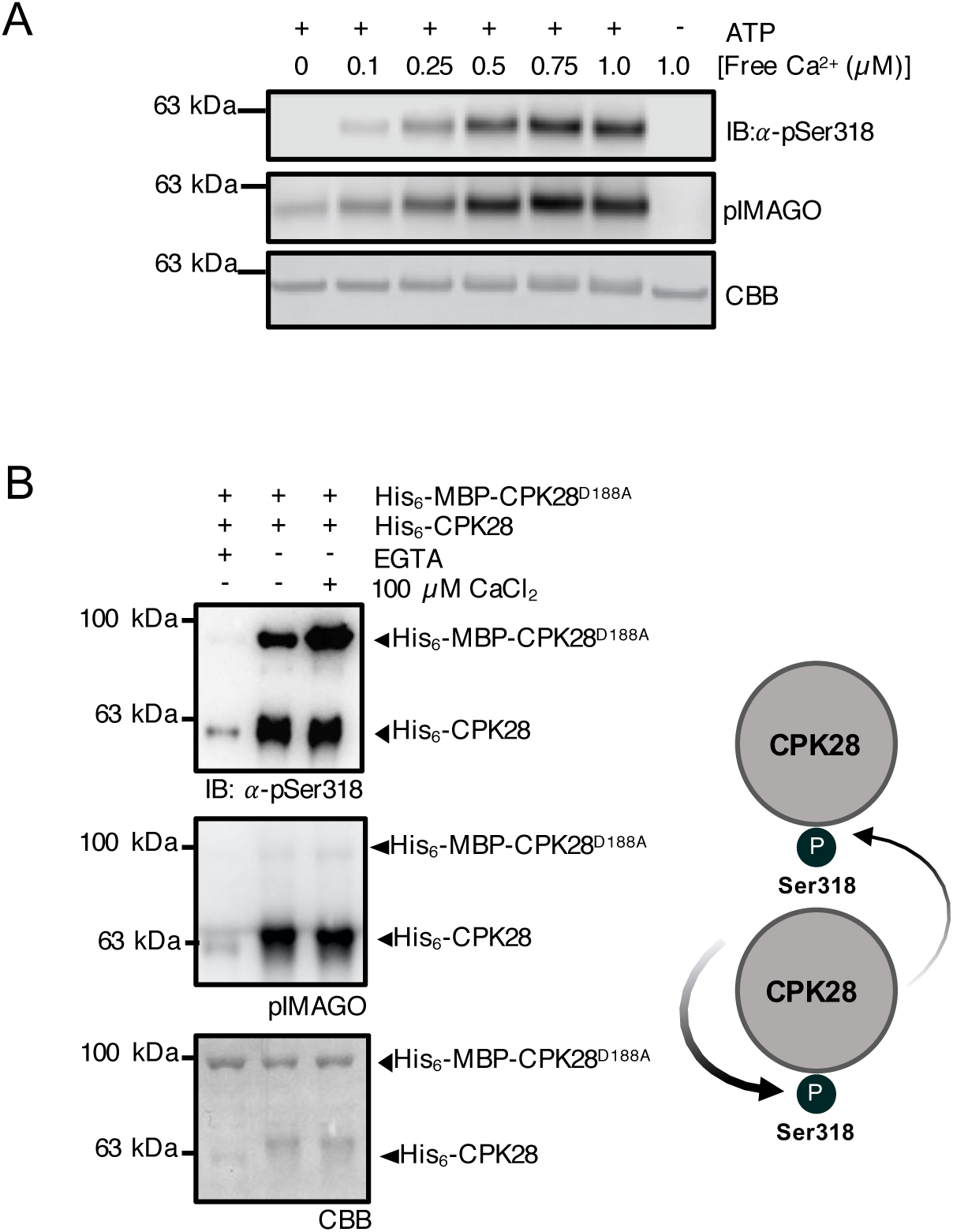
CPK28 undergoes intra- and inter-molecular autophosphorylation at Ser318. *(A)* His_6_- CPK28 autophosphorylation under increasing Ca^2+^ concentrations. *(B) In vitro* kinase assays using recombinant His_6_-CPK28, His_6_-CPK28^S318A^, or His_6_-CPK28^D188A^ and His_6_-MBP-CPK28^D188A^. Blots were probed with pIMAGO for the detection of phosphoproteins or *α*-pSer318 (1:5,000) antibody. Nylons were stained with Coomassie brilliant blue (CBB) to assess protein loading. Experiments were conducted three times with similar results.

### BIK1 can phosphorylate CPK28 on Ser318

CPK28 is a highly active kinase even at basal cellular levels of Ca^2+^ (49, 53). Increasing [Ca^2+^], either by adding CaCl_2_ to *in vitro* kinase assays (53), or via immune treatment *in vivo* (49), increases overall phosphorylation on CPK28, including on Ser318 (Fig 2A). Although CPK28 can autophosphorylate on Ser318, it is possible that this site is phosphorylated by additional protein kinases. BIK1 is a critical convergent substrate of multiple immune receptors whose activity and abundance is tightly regulated by layers of dynamic post-translational modifications including phosphorylation/dephosphorylation (57–61) and mono- (62) and poly-ubiquitination (49, 50, 63). CPK28 phosphorylates both BIK1 (50) and the E3 ubiquitin ligases PUB25 and PUB26 (49). A recent study demonstrated reciprocal phosphorylation between the rice orthologs of CPK28 and BIK1, OsCPK4 and OsRLCK176 (64), leading us to hypothesize that a similar mechanism may exist in Arabidopsis. To test if CPK28 is a substrate of BIK1, we conducted *in vitro* kinase assays using recombinantly produced GST-BIK1 and catalytically inactive His_6_-CPK28^D188A^. Phospho-tag gel stain for the detection of phosphorylated proteins indicated that BIK1 is indeed able to phosphorylate CPK28 *in vitro* (Fig 3A). Next, we tested if BIK1 can phosphorylate Ser318 by conducting *in vitro* kinase assays between GST-BIK1 and His_6_-CPK28^D188A/S318A^ compared to His_6_- CPK28^D188A^. We observed comparably less phosphorylation when CPK28^D188A/S318A^ was used as a substrate (Fig 3A), suggesting that Ser318 can be phosphorylated by BIK1. Furthermore, immunoblot analysis using anti-pSer318 confirmed that BIK1 is capable of phosphorylating Ser318 *in vitro* (Fig 3B). As we still observed some level of transphosphorylation by BIK1 on CPK28^D188A/S318A^ (Fig 3A), Ser318 is likely not the only site BIK1 phosphorylates on CPK28. Additional BIK1-mediated phosphorylation sites on CPK28 await to be discovered.

**Figure 3.**
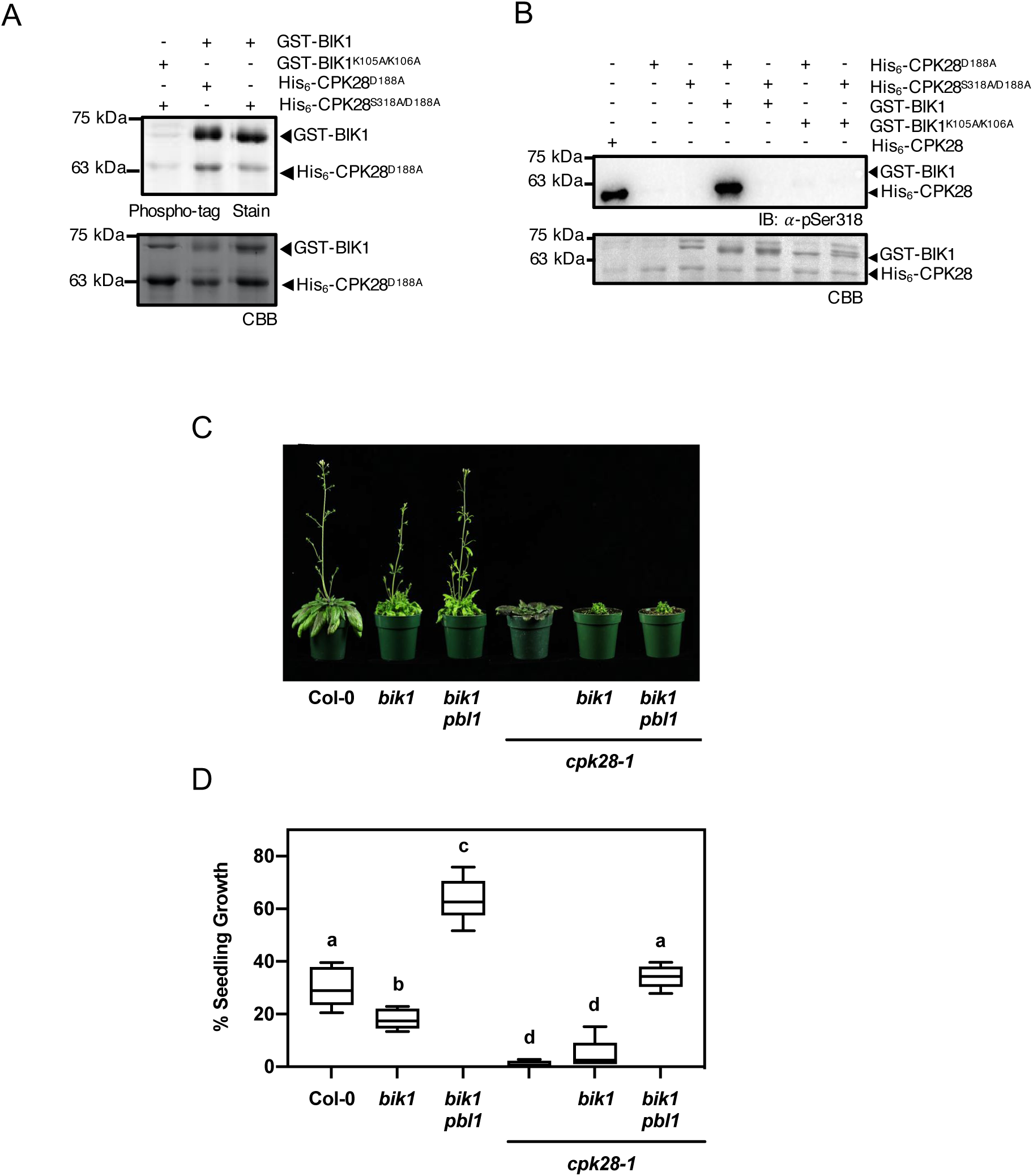
BIK1 transphosphorylates CPK28 at Ser318 and is required for CPK28-dependent immune signaling. *(A)* Phospho-tag gel stain and *(B)* Western blot analysis (*α*-pSer318) of *in vitro* kinase assay using recombinant GST-BIK1 or GST-BIK1^K105A/K106A^ and His_6_-CPK28^D188A^ or His_6_-CPK28^S318A/D188A^. In order to visualize both Ser318 autophosphorylation and GST-BIK1 transphosphorylation, 500 ng of His_6_-CPK28 and and 2 µg of His_6_-CPK28^D188A^ were loaded for auto- and trans-phosphorylation reactions, respectively. Gels and nylons were stained with Coomassie Brilliant Blue (CBB) to assess protein loading. *(C)* Stem-elongation of six week old plants and *(D)* seedling growth following continual treatment with 500 nM AtPep1 for 12 days (n=6). Values were compared to plants grown without AtPep1 and presented as boxplots indicating first and third quartiles, median values, and whiskers representing maximum and minimum values. Statistically different (p<0.005) values are denoted by lowercase letters according to an ANOVA analysis followed by Tukey’s posthoc test. All experiments were conducted at least three times with similar results.

BIK1 is part of a large gene family in Arabidopsis, and shares biological function with its closest homolog PBL1 (57, 65–67). To investigate the genetic requirement of BIK1 and PBL1 on CPK28-mediated signaling, we generated both *bik1 cpk28-1* double and *bik1 pbl1 cpk28-1* triple mutants and assessed whether loss of BIK1/PBL1 was able to suppress *cpk28-1* phenotypes. In congruence with our finding that Ser318 differentially regulates CPK28 function in immune homeostasis, we found that delayed stem elongation was not suppressed in *bik1 cpk28-1* or *bik1 pbl1 cpk28-1* (Fig 3C), but that AtPep1-triggered seedling growth inhibition was partially or fully restored in *bik1 cpk28-1* and *bik1 pbl1 cpk28-1*, respectively (Fig 3D). These data suggest that the function of CPK28 in immune signaling, but not in stem elongation, is dependent on BIK1/PBL1, and provides further evidence for complex regulatory feedback between BIK1/PBL1 and CPK28.

### Phosphorylation of Ser318 primes CPK28 Ca^2+^-responsiveness

We previously reported on the phosphorylation-dependent Ca^2+^-sensitivity priming of CPK28 peptide kinase activity (53), and were interested to understand which phosphorylation site or sites mediated this priming function. We suspected that a phosphorylation site in either the protein kinase domain, the AIJ, or the CLD would be responsible for Ca^2+^-priming and therefore generated individual phospho-null (Ser/Thr-to-Ala) mutants for autophosphorylation sites within these domains of CPK28 that we identified from *in situ* phosphorylated recombinant protein (Fig S5A) (53). Wild-type hyperphosphorylated CPK28 is insensitive to the addition of excess Ca^2+^ in peptide kinase assays (Fig S5B) (53). We thus hypothesized that phospho-null mutation of the phosphorylation site(s) responsible for Ca^2+^-sensitivity priming would restore CPK28 activation by excess Ca^2+^. To test this hypothesis, we expressed hyperphosphorylated forms of each phospho-null mutant and compared peptide kinase activity in untreated samples versus samples supplemented with 100 µM CaCl_2_. Much to our surprise, our biochemical analysis converged on Ser318 as a critical regulatory phosphorylation site of CPK28. Of all the phospho-null mutants tested, only the S318A mutant had enhanced peptide kinase activity upon addition of excess Ca^2+^. We therefore further characterized the CPK28^S318A^ mutant for Ca^2+^-dependent autophosphorylation and for peptide kinase activity at different concentrations of Ca^2+^.

To confirm that phosphorylation of Ser318 plays a role in Ca^2+^-responsiveness, we used *Escherichia coli* Lambda phosphatase (LamP)-expressing cells to produce dephosphorylated His_6_-CPK28 and His_6_-CPK28^S318A^ and conducted comparative *in vitro* autophosphorylation assays either in the complete absence of Ca^2+^ (+ 10 mM EGTA), in the presence of background Ca^2+^ (no treatment), or with the addition of excess Ca^2+^ (+ 100 μM CaCl_2_). Autophosphorylation levels were detected using pIMAGO. Both His_6_- CPK28 and His_6_-CPK28^S318A^ showed low levels of autophosphorylation in chelation experiments (Fig 4A), confirming their Ca^2+^-dependence. His_6_-CPK28 was highly active at background Ca^2+^ and was not substantially stimulated by additional CaCl_2_ (Fig 4A). In contrast, at background Ca^2+^ levels, His_6_-CPK28^S318A^ displayed dramatically reduced autophosphorylation activity compared to wild-type protein (Fig 4A). However, when assays were conducted in the presence of excess Ca^2+^, His_6_-CPK28 and His_6_- CPK28^S318A^ exhibited comparable levels of autophosphorylation (Fig 4A). Together, these data suggest that phosphorylation of Ser318 is required for full activity when Ca^2+^ levels are not saturating. Interestingly, when autophosphorylation assays were allowed to proceed for longer time intervals (30 min and 1 hr), His_6_-CPK28^S318A^ displayed similar levels of activity as wild-type protein at basal [Ca^2+^] (Fig 4A). Collectively, these data suggest that phosphorylation of Ser318 is important for the rapid autophosphorylation of CPK28 under limiting [Ca^2+^].

**Figure 4.**
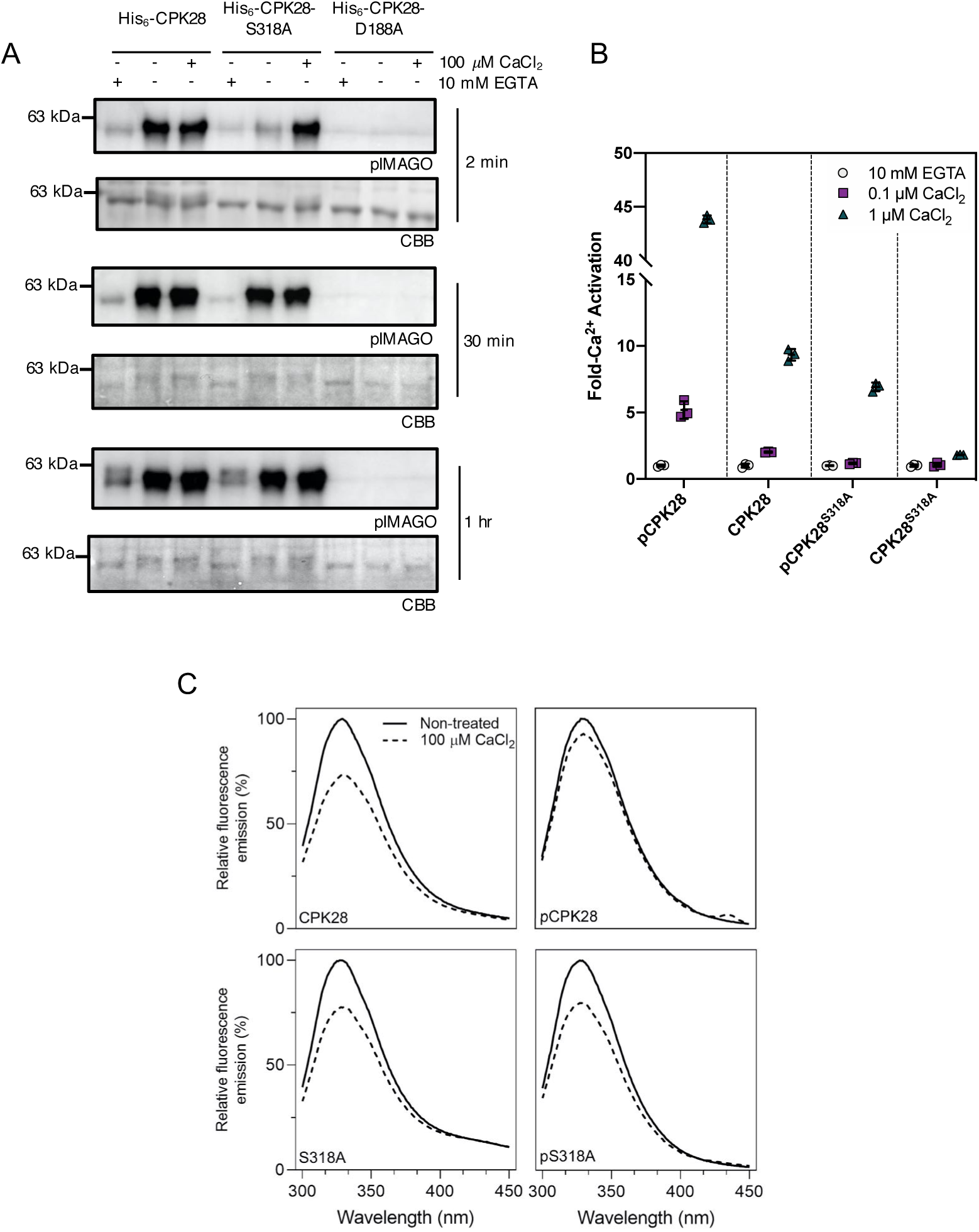
Phosphorylation at Ser318 primes CPK28 Ca^2+^-responsiveness. *(A)* Autophosphorylation of recombinant His_6_-CPK28 or His_6_-CPK28^S318A^ without Ca^2+^ (+10 mM EGTA), at background Ca^2+^ (-EGTA, - CaCl_2_), or with the addition of 100 µM CaCl_2_ for 1-30 min as indicated. Kinase dead His_6_-CPK28^D188A^ was used as a negative control. Phosphorylated proteins were detected using pIMAGO and nylons were stained using Coomassie Brilliant Blue (CBB) to assess protein loading. *(B)* Peptide kinase activity of phosphorylated (pCPK28, pS318A) and dephosphorylated (CPK28, S318A) purified recombinant proteins at physiological Ca^2+^ concentrations. Free Ca^2+^ concentrations were established as described in the Experimental Procedures. Mean with standard deviation of three replicate reactions as well as individual values are shown. The experiment was repeated twice with independent preparations of all recombinant proteins with similar results observed each time. *(C)* Intrinsic Trp fluorescence of dephosphorylated (CPK28, S318A) or hyperphosphorylated (pCPK28, pS318A) His-tagged recombinant proteins before (‘non-treated’, solid line) or after (dashed line) the addition of 100 µM CaCl_2_. Curves are averages of two scans following appropriate background subtraction. Experiments were performed twice using independent preparations of recombinant proteins with similar results observed in both experiments.

To better understand how phosphorylation of Ser318 affects activation of CPK28 by Ca^2+^, we assessed CPK28 peptide kinase activity using the ACSM+1 synthetic peptide as substrate (53) at Ca^2+^ concentrations that would be expected at the lower (0.1 μM) and upper (1.0 μM) range of intracellular physiological conditions. Phosphorylated and dephosphorylated forms of both His_6_-CPK28 and His-CPK28^S318A^ were used to determine if overall phosphorylation status could supersede the requirement for site-specific phosphorylation at Ser318. At free Ca^2+^ concentrations of 0.1 and 1.0 μM, dephosphorylated His_6_-CPK28 (Fig 4B) displayed ∼2- and 9-fold activation by Ca^2+^ relative to EGTA-treated protein, respectively (Fig 4B). Similarly, phosphorylated CPK28 (Fig 4B) had ∼5-fold activation with the addition of 0.1 μM free Ca^2+^ and ∼43-fold activation with 1.0 μM free Ca^2+^ (Fig 4B), indicating that phosphorylation enhances CPK28 peptide kinase activity at physiological Ca^2+^. By comparison, His_6_-CPK38^S318A^ was substantially less active at physiological Ca^2+^, having negligible activity at 0.1 µM Ca^2+^ (relative to EGTA-treated protein) regardless of its overall phosphorylation status and only ∼2 and 7-fold activation for dephosphorylated and phosphorylated His_6_-CPK28^S318A^, respectively, at 1.0 µM Ca^2+^. Collectively, these results indicate that phosphorylation of Ser318 is prerequisite for substrate phosphorylation by CPK28 at low free Ca^2+^, and that phosphorylation of Ser318 is required for full responsiveness to Ca^2+^ elevations within the physiological range. Taken together, these data support the hypothesis that phosphorylation of Ser318 is uniquely important in priming CPK28 for activation by Ca^2+^ at concentrations that would be expected under cellular conditions.

### Ser318 phosphorylation promotes a Ca^2+^-bound conformation

Ser318 is located at the C-terminal end of the canonical protein kinase domain of CPK28, very close to the autoinhibitory junction (AIJ) (Fig 1A), suggesting that phosphorylation of Ser318 might affect Ca^2+^-dependent conformational changes of the AIJ-CAD fragment of CPK28. To better understand the biochemical function of Ser318 phosphorylation, we measured Ca^2+^-induced conformational changes in hyper- and hypo-phosphorylated CPK28 and CPK28^S318A^ by intrinsic Trp fluorescence. Trp fluorescence emission properties depend on the local environment of Trp residues, and can thus be used to assess protein conformational changes (68). For dephosphorylated His_6_-CPK28 and His_6_-CPK28^S318A^ (Fig 4C), Trp fluorescence emission decreased following the addition of 100 µM CaCl_2_ (relative to non-treated protein), indicating that CPK28 undergoes a Ca^2+^-dependent conformational change. By comparison, Trp fluorescence of hyper-phosphorylated His_6_-CPK28 showed only a marginal decrease after addition of excess Ca^2+^ (Fig 4C), suggesting that CPK28 phosphorylation promotes a Ca^2+^-bound conformation at low levels of Ca^2+^. In agreement with our activity assays, Trp fluorescence of hyper-phosphorylated His_6_-CPK28^S318A^ decreased similar to the wild-type dephosphorylated protein, suggesting that phosphorylation of Ser318 is responsible for the effect observed with hyperphosphorylated CPK28.

### Ser318 is a conserved and unique feature of subgroup IV CDPKs

CDPK gene families are highly conserved across land plants and form four major subgroups (26, 27). To determine the level of conservation of Ser318, the amino acid sequences of CPK28 and other subgroup IV orthologs from all genomes available on Phytozome were compared. Amongst the 114 sequences included in our analysis, a Ser residue was strictly conserved at the position orthologous to Ser318 of AtCPK28 (Fig 5A and Fig S1). Comparison of all subgroup I-III CDPKs from 12 representative species spanning all major taxonomic groups indicated that although several flanking residues are highly conserved in all subgroups, conservation of Ser318 is a unique and specific feature of subgroup IV CDPKs (Fig 5A).

**Figure 5.**
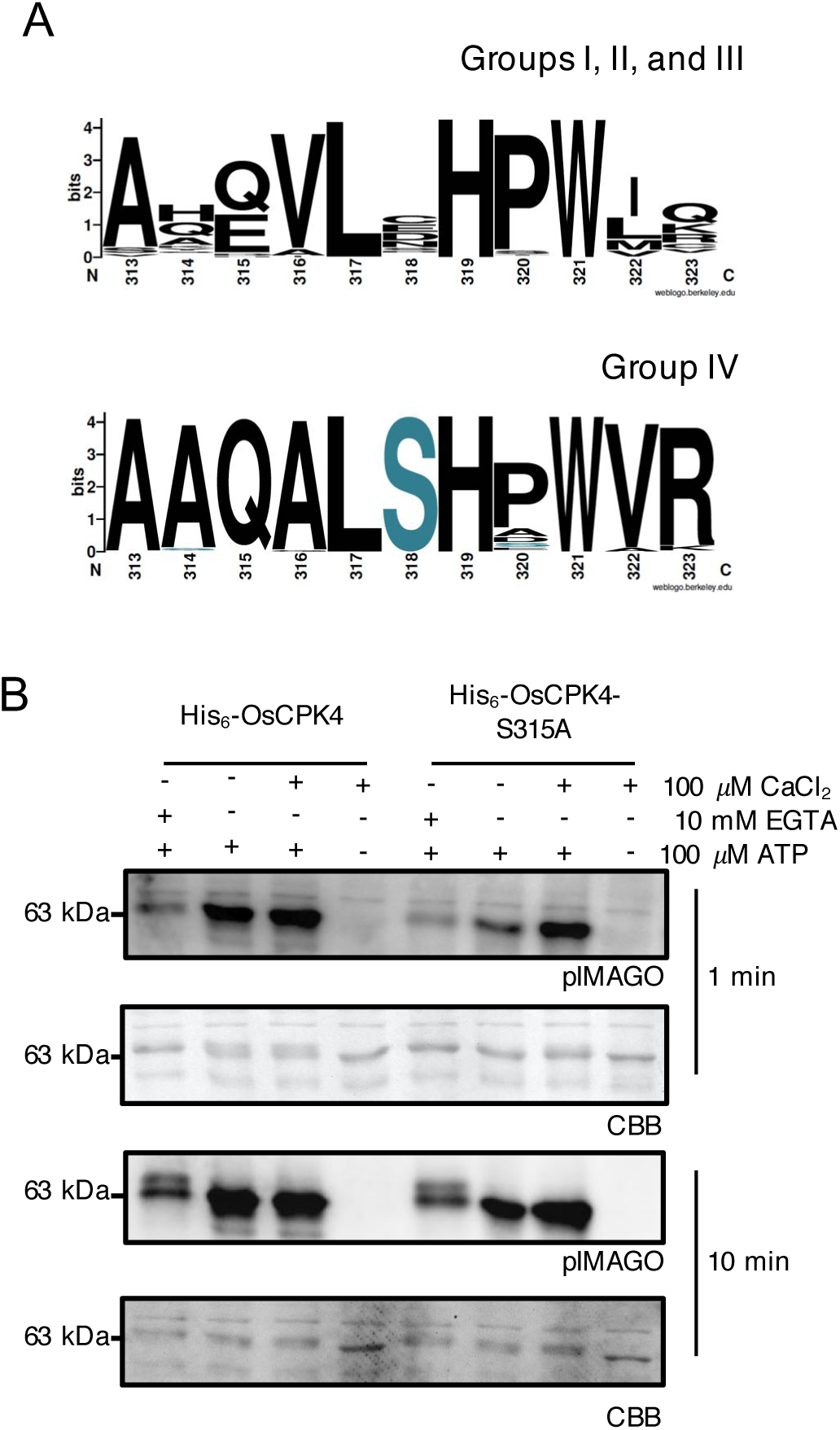
Ser318 is a conserved feature of group IV CDPKs. *(A)* Amino acid consensus at position 318 (AtCPK28) across CDPK subgroups. Sequences were retrieved from Phytozome and aligned as described in the Experimental Procedures. Logos were generated using WebLogo (97). *(B)* Autophosphorylation of recombinant His_6_-OsCPK4 and His_6_-OsCPK4^S315A^. Reactions were carried out in the absence of Ca^2+^ (+10 mM EGTA), in the presence of background Ca^2+^(-EGTA, -CaCl_2_), or with the addition of 100 µM CaCl_2_ for 1 or 10 min as indicated in figures. Phosphorylation was detected using pIMAGO. Nylons were stained with Coomassie Brilliant Blue (CBB) to assess protein loading. All experiments were conducted at least three times with similar results.

To determine if the Ca^2+^-priming function of Ser318 autophosphorylation is conserved, we generated a phospho-ablative variant of the rice CPK28 ortholog (OsCPK4^S315A^). Short 1 min autophosphorylation assays were conducted using recombinant His_6_-OsCPK4 and His_6_-OsCPK4^S315A^ produced in LamP-expressing *E. coli* cells, as described above. His_6_-OsCPK4 displayed a clear requirement for Ca^2+^ with a marked decrease in overall phosphorylation in the presence of 10 mM EGTA (Fig 5B). While wild-type His_6_-OsCPK4 displayed high levels of autophosphorylation at basal [Ca^2+^], His_6_-OsCPK4^S315A^ was comparatively less active (Fig 5B). The addition of 100 μM CaCl_2_ did not further activate His_6_-OsCPK4 but did restore His_6_-OsCPK4^S315A^ autophosphorylation levels to that observed with wild-type protein (Fig 5B). Together, these data provide evidence of a conserved biochemical function for phosphorylation of this residue in orthologous CDPKs across the plant lineage.

## DISCUSSION

Expansion of CDPKs in plants is predicted to have occurred under selective adaptation for kinases with varying Ca^2+^ sensitivities (27), however the biophysical properties underlying Ca^2+^ sensitivity are not fully understood. Analysis of protein sequences from Arabidopsis indicates that CDPKs with little or no requirement for Ca^2+^ possess one or more degenerated EF-hand motifs (37, 69). However, some CDPKs with the ability to bind Ca^2+^ do not require it for their activation (30, 70–72) pointing to additional mechanisms regulating CDPK function. Previous work has shown that *in situ* autophosphorylation “primes” Arabidopsis CPK28 for Ca^2+^ activation *in vitro* (53). Here, we demonstrated that phosphorylation at one site, CPK28-Ser318, is responsible for autophosphorylation-based priming when Ca^2+^ concentrations are limiting (Fig 4A-B). Additionally, *in vivo* phosphorylation at Ser318 was required for CPK28 function in immune homeostasis (Fig 1C-E) but not stem elongation (Fig 1B), suggesting a role in stimulus-specific activation of a multifunctional protein kinase through Ca^2+^-sensitivity priming.

Autophosphorylation has been correlated with the activation (15, 53, 73–75) or inhibition (72, 75, 76) of several CDPKs, although the mechanisms of regulation remain largely unknown. Phosphorylation could influence interactions with protein substrates (75, 77) or induce changes in secondary protein structure causing transitions between functional enzyme states (78, 79). A complete crystal structure for a plant CDPK has not yet been resolved; however, a mechanism for Ca^2+^ activation has been proposed based on the structures of apicomplexan CDPKs (31–33). Experiments using *Toxoplasma gondii* TgCDPK1/2 and *Cytosporidium parvum* CpCDPK1 demonstrate that Ca^2+^ activation is reversible with contact sites between the CaM-like domain and kinase domain stabilizing both active and inactive forms (31). Many of the residues that stabilize these conformations are conserved between plants and protists (32) suggesting similar contact sites may exist in plants.

CPK28-Ser318 resides in the C-terminal portion of the kinase domain in close proximity to the AIJ (Fig 1A). Although we could not generate a high confidence structural model of CPK28 using the crystal structures of Ca^2+^-bound CDPKs, modeling of CPK28 using inactive TgCDPK1 (31) indicated that Ser318 is likely surface-localized, directed away from the active site of the kinase domain (Fig S6). In this position, phosphorylated Ser318 would not interact with established contact sites, such as the autoinhibitory triad (31), or other interactions between the pseudosubstrate region and the active site of the kinase domain. We rather propose that phosphorylation of Ser318 could induce a structural change in the AIJ that prohibits stabilization of the inactive conformation. This could cause the protein to adopt an “intermediate” conformation that can more readily move to the active state upon Ca^2+^ binding (Fig 6). In support of this idea, our analysis of intrinsic Trp fluorescence of CPK28 and CPK28^S318A^ suggests that Ca^2+^-dependent conformational changes can occur at a lower concentration of Ca^2+^ in the hyper- compared to de-phosphorylated protein and that the conformational change at low Ca^2+^ requires Ser318 phosphorylation (Fig 4C). Phosphorylation of Ser318 could also stabilize the active conformation, serving a similar function to autophosphorylation of a residue in the autoinhibitory region (Thr286) of CaMKII from rat brain, which renders CaMKII substrate phosphorylation independent of both Ca^2+^ and CaM (80, 81).

**Figure 6.**
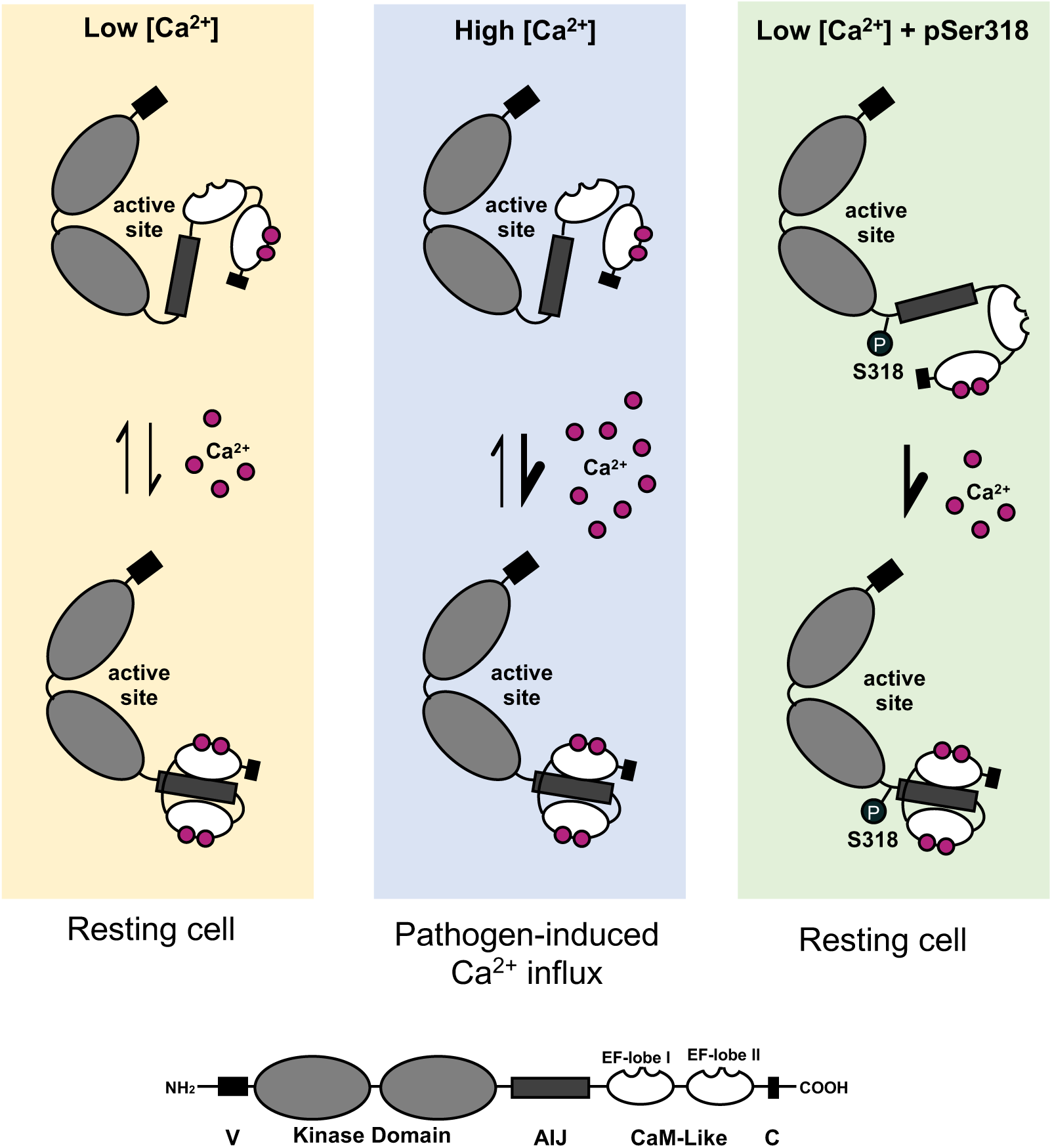
Model for CPK28 activation. Proposed mechanism of Ca^2+^-sensitivity priming of CPK28 by phosphorylation of Ser318. In a resting cell, CPK28 transitions between active and inactive conformations, binding and releasing Ca^2+^ from the amino-terminal EF-hand lobe of the calmodulin (CaM)-like domain. Ca^2+^ elevation during signaling events shifts this equilibrium towards the activated state via Ca^2+^-induced conformational changes. When Ser318 is phosphorylated, the transition of CPK28 from an inactive to an active state occurs at lower Ca^2+^ concentrations possible via stabilizing a conformation in which the AIJ is excluded from the active site.

Overall, our analysis of CPK28 autophosphorylation, peptide kinase activity, and conformational changes collectively suggest that phosphorylation of Ser318 near the AIJ promotes an open, active conformation of CPK28 at low Ca^2+^ concentrations (Fig 6). Currently, it is not clear whether this conformational state results in enhanced affinity of the CaM-like domain for Ca^2+^, or whether pSer318 exerts allosteric effects promoting release of the AIJ from the kinase domain. Notably, we recently demonstrated that CaM binds to an area of the CPK28 AIJ to inhibit both *in vitro* auto- and trans-phosphorylation activity (53), and that CPK28 autophosphorylation relieves this inhibition (53). Ser318 phosphorylation could conceivably block CaM binding or, alternatively, CaM could prevent Ser318 phosphorylation. Resolving an intact crystal structure for CPK28, and other plant CDPKs, will provide valuable insight into the activation of these kinases and allow for investigation into the possible structural roles of phosphorylation.

From a physiological perspective, phosphorylation of Ser318 would allow CPK28 to be active in a cell under resting conditions. Indeed, CPK28 promotes the degradation of BIK1 prior to the activation of immune signaling (49, 50). Limiting BIK1 accumulation is critical to prevent mounting an immune response in the absence of pathogen invasion. Ser318 was identified as an *in vivo* phosphorylation site in unstimulated cells (51), indicating no requirement for immune activation. A 25-fold increase in phosphorylated peptides corresponding to pSer318 were identified in *cpk28-1* protoplasts expressing CPK28-YFP compared to the kinase inactive variant (CPK28^D188A^-YFP), suggesting that Ser318 is an autophosphorylation site *in vivo* (51). Our *in vitro* data indicates that CPK28 undergoes both intra- and inter-molecular autophosphorylation at Ser318 (Fig 2) and can also be transphosphorylated by BIK1 (Fig 3A and 3B), although a higher level of phosphorylation was detected on autophosphorylated CPK28 in our assays (Fig 3B). Whether Ser318 phosphorylation occurs by autophosphorylation and/or BIK1-mediated transphosphorylation *in vivo* is unknown, but our *in vitro* analyses clearly establish the potential for both mechanisms as drivers of Ser318 phosphorylation *in vivo*. How these distinct events would contribute to CPK28-mediated BIK1 proteostasis remains an open and challenging question.

Nevertheless, it is tempting to speculate that BIK1 transphosphorylation could contribute to CPK28 regulation. Our epistasis analysis indicates that CPK28-mediated immune signaling is dependent on BIK1 and its homolog, PBL1 (Fig 3D). However, loss of *bik1 pbl1* in *cpk28-1* mutants restored *cpk28-1* growth inhibition to levels observed in Col-0 seedlings (Fig 3D), suggesting the involvement of additional RLCKs in response to endogenous immune peptide elicitation. In rice, OsRLCK176 phosphorylates OsCPK4 at three sites: Thr73, Ser210 and Ser381 (64), corresponding to CPK28-Thr76, -Ser213, and -Val384. Phosphorylation at these sites activates OsCPK4 as part of a regulatory feedback loop to control immune output through continual degradation of OsRLCK176 (64). Our *in vitro* kinase assays suggest that BIK1 phosphorylates CPK28 at Ser318 and additional, currently unknown, residues (Fig 3A). Given the high conservation of this immune signalling network (82), Thr76 and Ser213 are likely candidates for future analysis. In Arabidopsis, BIK1 turnover is mediated by CPK28-dependent phosphorylation and activation of the E3 ubiquitin ligases PUB25 and PUB26, which target BIK1 for 26S proteasomal degradation (49). Previous work indicates that CPK28 is also capable of phosphorylating BIK1 *in vitro* (50), although the biochemical and biological consequences of this transphosphorylation are not known. It is therefore plausible that BIK1 accumulation could be modulated by an interplay of BIK1 ubiquitination and trans-phosphorylation events between CPK28 and BIK1, which is a current area of investigation.

Our cumulative data indicate that phosphorylation of Ser318 would render CPK28 highly responsive to slight increases in Ca^2+^ following immune activation, preventing initiation of a robust immune response. Ser318 phosphorylation could additionally slow CPK28 deactivation following a primary pathogen attack, making plants more susceptible to secondary infection. In accordance with its role in BIK1 turnover, overexpression of CPK28 dampens the MAMP-triggered Ca^2+^ (48) and oxidative (50) bursts, which are dependent on BIK1 and PBL1 (48, 50). This implies that CPK28 overactivation could additionally prevent the establishment of systemic acquired resistance which is reliant on ROS/Ca^2+^ signal propagation (34, 83, 84). In order to buffer these attenuation mechanisms the plant would need to adopt a safeguard against sustained activation of CPK28. Recent work has demonstrated that following cell stimulation with AtPep1 (85) or the bacterial flagellin immunogenic epitope flg22 (86), alternative *CPK28* transcripts are produced that generate a truncated isoform that lacks the C-terminal EF-hand lobe. This CPK28 variant could act to outcompete Ca^2+^ “competent” proteins for downstream substrates, alleviating immune attenuation (87). Thus, CPK28 signaling appears to be intricately regulated at both the post-transcriptional and post-translational levels.

Despite the clear requirement for Ser318 phosphorylation in immune signaling (Fig 1C-E), *cpk28-1* plants expressing CPK28^S318A^ displayed normal stem elongation (Fig 1B). The catalytic activity of CPK28 is indispensable for all known biological functions (50, 51); however, the lower kinase activity of CPK28^S318A^ at physiologically relevant Ca^2+^ concentrations (Fig 4A-B) did not impair stem elongation (Fig 1B), suggesting pathway-specific requirements for full CPK28 catalytic activity. We speculate that more processive genetic programs, such as those that occur in developmental and reproductive processes, would not require CDPKs to be as responsive to Ca^2+^ as stress-induced signals. In support of this hypothesis, prolonged exposure to low levels of Ca^2+^ relieved the requirement of Ser318 phosphorylation for full kinase activity in autophosphorylation assays (Fig 4A). In a physiological context, the phosphorylation status of CPK28 is likely dictated by a combination of kinase and phosphatase activities to precisely control CPK28 function. Although pathway-specific CPK28 binding partners have not yet been identified in the reproductive phase transition, it is possible that such associations could increase kinase activity or lower Ca^2+^ requirements.

The development of pathogen resistant crops can be complicated by growth trade-offs associated with the overactivation of immune signalling (88, 89). For example, prolonged activation of immune receptors by MAMP stimulation causes seedling growth inhibition (90, 91) and the formation of lesions (92, 93). Ablation of Ser318 phosphorylation allowed us to generate a *CPK28* allele that displays enhanced resistance to *Pto* DC3000 (Fig 1E) with no consequences to plant growth (Fig 1B). Group IV CDPKs are highly conserved across all land plants (26) and fulfill conserved roles as regulators of immune signaling (82) and reproductive development (51, 94–96) in multiple plant species. Accordingly, our *in vitro* analysis of OsCPK4 indicated that, on a biochemical level, phosphorylation of Ser315 (analogous to CPK28-Ser318) has a conserved function in rice (Fig 4D). Additionally, a Ser residue at this site was identified as a unique feature of group IV CDPKs across all surveyed land plants (Fig 4C). Cumulatively, our data suggests that ablation of “Ser318” phosphorylation in species with agricultural value could serve as an effective tool for the development of disease resistance without associated costs to fitness or yield.

## EXPERIMENTAL PROCEDURES

### Plant growth conditions

*Arabidopsis thaliana* plants were grown either in soil or sterile media depending on the assay. For soil assays, seeds were stratified for 2-3 days at 4°C, sown on soil, and transplanted as one plant per pot in Sunshine Mix 1 soil (Sungro) for 5 weeks in temperature-controlled growth chambers in the Queen’s University Phytotron at 22°C with 10 h light (150-160 μE m^2^ s^-1^) and no humidity control. Oxidative burst and pathogen infection assays were conducted on soil-grown plants 3-4 weeks post germination (wpg), prior to reproductive stage transition. For assessment of stem elongation, plants were subsequently transferred to a growth chamber maintained at 22°C, with a 16 h photoperiod (150-160 μE m^2^ s^-1^), and 30% relative humidity for 1-2 weeks or until all plants had produced reproductive bolts. For sterile assays, seeds were surface-sterilized using 50% bleach, stratified for 2-3 days at 4°C, and germinated on 0.8% agar plates with 0.5x Murashige and Skoog (MS) media for 3-4 days and then transplanted into liquid 0.5x MS supplemented with 1% sucrose. Sterile seedlings were grown at ambient temperature with 10 h light 150-160 μE m^2^ s^-1^ and assays were conducted at 2 wpg. Soil-grown *A. thaliana* plants were fertilized biweekly with 1.5 g L^-1^ 20-20-20 NPK. Predatory *Amblyseius swirskii* mites (Koppert Biological Systems) were released into growth chambers biweekly as a precautionary measure against greenhouse pests, according to manufacturer’s instructions.

### Plant materials

Stable *A. thaliana* transgenics were generated via *Agrobacterium tumefaciens* (GV3101)-mediated floral dip transformation (97). T_1_ plants were selected on MS agar plates containing 50 μg/mL hygromycin. Only lines displaying a 3:1 segregation ratio on selective media in the T_2_ generation were bred to homozygosity and used in complementation experiments. Stable *cpk28-1/35S:CPK28-YFP*, *cpk28-1/35S:CPK28^S318A^-YFP*, and *cpk28-1/35S:CPK28^D188A^-YFP Arabidopsis* lines were previously described (51). Higher-order mutants were generated by crossing *bik1* or *bik1 pbl1* (65) mutants with *cpk28-1* (51) and bred to homozygosity using PCR-based genotyping. Table S1 includes a list of all germplasm used and generated in this study.

### Molecular cloning

The *pCPK28:CPK28-FLAG* construct was cloned by fusing the coding sequence of *CPK28* downstream of its native promoter (1742 bp upstream of the start codon) and in frame with a C-terminal FLAG peptide using digestion-ligation cloning into a pGREENII-based binary vector carrying the aminoglycoside phosphotransferase gene from *E. coli* for hygromycin B resistance in plants (98). Both the *pT7:His_6_-CPK28* construct in pET28a+ (EMD Biosciences), and the *pT7:MBP-His_6_-CPK28* construct in pOPINM (Novagen), have been described previously (50, 53). Site-directed mutagenesis was used to generate CPK28 mutant constructs using overlapping complementary primers as described previously (50), using either pGREENII-based binary plasmids or pET28a+ clones as the template. The coding region of *BIK1* was PCR-amplified from previously described pENTR-BIK1 or pENTR-BIK1^K105A/K106A^ vectors (65) and cloned into bacterial expression vector pGex6.1 (GE Healthcare) by Gibson Assembly (NEB) to generate *pT7:GST-BIK1* and catalytically-inactive *pT7:GST-BIK1^K105A/K106A^* constructs. *pET100:His_6_-OsCPK4* and *pET100:His_6_-OsCPK4^S315A^* constructs were synthesized by GeneArt^TM^ (Fisher Scientific). The coding region of PUB25 was PCR-amplified from previously described pET28a:PUB25 vectors (49) and cloned into bacterial expression vector pMAL-c2x (GE Healthcare) by digestion and ligation cloning using *Xba*I and *Pst*I-HF (NEB) to produce pMAL-c2x:MBP-PUB25. PUB26 was PCR amplified from pCAMBIA1300-35S:PUB26-FLAG (49) and cloned into pMAL-c2x using *Xba*I and *Pst*I-HF to produce pMAL-c2x:MBP-PUB26. All clones were confirmed by Sanger Sequencing using plasmid- and/or gene-specific primers (The Center for Applied Genomics, Hospital for Sick Children, Toronto Canada, or Eurofins Genomics, Ebersberg, Germany). All primers used for cloning are listed in Table S1.

### Confocal Microscopy

Leaf discs were sampled from 4-6 week-old Arabidopsis soil grown plants using a 4 mm biopsy punch (Integra Miltex) and were wet-mounted in water with the abaxial surface facing upwards prior to confocal imaging. Imaging was performed using a LSM 710 (Zeiss) confocal microscope with excitation at 488 nM for YFP and a range of 510-540 nM for measuring emission. To detect chlorophyll autofluorescence, an excitation wavelength of 543 nM and a range of 680-760 nM for detecting emission was used.

### Immune assays

AtPep1(99) and elf18 (100) used for immune assays were synthesized by EZBiolab (USA). AtPep1-induced SGI and ROS burst assays were performed as previously described (101). Infection assays were conducted using virulent *Pseudomonas syringae* pv. *tomato* (*Pto* DC3000) on soil-grown *A. thaliana* plants using a needless syringe as outlined previously (50).

### Generation of the anti-pSer318 antibody

The CPK28 pSer318 antibody was generated and purified by LifeTein (New Jersey, USA). Briefly, rabbits were immunized with KLH-coupled synthetic peptide corresponding to the region of CPK28 surrounding Ser318 (NH_2_- CKDPRARLTAAQAL**pS**HAWV-COO-). KLH coupling was facilitated by the addition of Cys to the N-terminus of the peptide. Antibody was purified first by enrichment against the pSer318 phospho-peptide and then by negative enrichment against an unphosphorylated version of the peptide to remove non-phospho-specific IgGs. Purified antibodies were validated by immunoblotting against wildtype, kinase-dead (K91E), or S318A recombinant CPK28 proteins. Purified antibodies were determined to be phosphorylation and site-specific.

### Recombinant protein expression and purification

All recombinant CPK28 clones were transformed into lambda phosphatase-expressing BL21 (DE3) *E.coli* cells for the production of dephosphorylated proteins or into T7 Express cells (New England Biolabs) for production of hyperphosphorylated protein (53). Cultures were grown in Luria-Burtani (LB) broth at 37 °C to an OD_600_ of ∼0.6-0.8. Expression was induced using 1 mM of β-D-1-thiogalactopyranoside (IPTG) for 16-18 h at room temperature with gentle shaking. Bacterial cells were harvested at 3,500 x *g* for 25 min at 4 °C and resuspended in 50 mL of extraction buffer containing 50 mM Tris-HCl (pH 7.5), 100 mM NaCl and 1 protease inhibitor cocktail tablet (SigmaAldrich). Cells were lysed by passing the resuspended culture through a French press (Glen Mills^®^High Pressure Cell Disruption) 3 times. Lysates were clarified by centrifugation at 35,000 x *g* for 40 min at 4 °C. His_6_-CPK28 proteins were immobilized on a nickel-nitrilotriacetic acid (NTA) gravity flow column (ThermoFisher) as described previously (53). Elution fractions were dialyzed against two exchanges of 2,500 volumes of 25 mM Tris-HCl (pH7.5), 50 mM NaCl and 1 mM DTT at overnight 4 °C. Recombinant production of GST-BIK1 and GST-BIK1-KD in BL21(DE3)-VR2-pACYC-LamP *E. coli* cells was conducted as described above in a phosphate buffered saline (PBS) solution (ThermoFisher) containing 1 mM DTT, 1 mM PMSF, and 6 mM MgCl_2_. Proteins were immobilized on a glutathione agarose gravity flow column (Qiagen) washed 5 times with PBS solution and eluted in 50 mM Tris-HCl (pH 8.0), 5 mM DTT, and 10 mM reduced glutathione. All proteins were concentrated using Pierce^TM^ 3000 MWCO concentration columns (ThermoFisher) to a final concentration of approximately 1.5 mg/mL, as determined by Bradford analysis (Bio-Rad) against bovine serum albumin standards. Recombinant production of MBP-PUB25 and MBP-PUB26 were conducted as described above with the following modifications. pMAL-c2x:MBP-PUB25/26 were transformed into BL21(DE3) *E. coli* cells. Cultures were grown in a baffled flask with LB broth at 37°C to an OD_600_ of ∼0.6-0.8 then expression was induced using 0.1 mM IPTG for 3 hours at room temperature with gentle shaking. Pelleted cells were resuspended in MBP column buffer containing 20 mM Tris HCl pH 7.4, 200 mM NaCl, 1 mM EDTA, 1 mM DTT, and 1 mM PMSF. MBP-PUB25/26 were purified using Amylose Resin (NEB) by batch purification according to manufacturer’s instructions. Protein was eluted using 100 µL elution buffer and used for assays on the same day as the purification. Purity was assessed by SDS-PAGE analysis followed by staining with Coomassie Brilliant Blue total protein stain. Protein aliquots were flash frozen in liquid N_2_ and stored at -80 °C until use.

### *In vitro* autophosphorylation assays

Autophosphorylation assays were conducted by incubating 5 μg of purified His_6_- CPK28/His_6_-OsCPK4 or mutant variants in a 50 μL reaction containing 25 mM Tris-HCl (pH7.5), 10 mM DTT, 100 μM ATP, and 100 μM CaCl_2_ or 10 mM EGTA, where specified. Proteins were allowed to autophosphorylate at room temperature for 1-60 min, as specified in figures. Reactions were stopped by the addition of 6x Laemmli sample buffer (LSB) and heating at 80 °C for 5 min. Reactions were analysed immediately or stored at -20 °C before SDS-PAGE and immunoblotting. Autophosphorylation was detected using the pIMAGO kit (Tymora Analytical) according to manufacturer’s instructions.

### *In vitro* trans-phosphorylation assays

Trans-autophosphorylation assays were performed using 2 μg of His_6_-MBP-CPK28 or His_6_-MBP-CPK28^D188A^ and 4 μg of His_6_-CPK28^D188A^ in the same reaction buffer as in the autophosphorylation assays at 30 °C for 30 min. Trans-phosphorylation assays were conducted using 2 μg of purified GST-BIK1 or GST-BIK1^K105A/K106A^ and 4 μg of His_6_-CPK28^D188A^ or His_6_-CPK28^D188A/S318A^ in a 20 μL total reaction volume of 25 mM Tris-HCl (pH 7.5), 10 mM MgCl_2_, 1 mM DTT and 100 μM ATP at 30 °C for 30 min. Reactions were stopped by adding 6x LSB buffer and heating at 80 °C for 5 min. Reactions were analysed immediately or stored at -20 °C before visualizing phosphorylated proteins using Phospho-Tag gel stain (APB Bio) according to the manufacturer’s instructions.

### Ca^2+^-activation assays

Analysis of CPK28 peptide kinase activity was carried out as exactly as previously described (53) using the ACSM+1 peptide (NH_2_-NNLRLSMGKR-COO^-^) as substrate. Briefly, reactions contained 40 mM Tris-HCl, pH 7.5, 1 mM DTT, 10 mM MgCl_2_, 100 μM ATP, 0.1 μCi/μl [γ-^32^P]ATP (150 cpm/pmol), 500 ng purified of His_6_-CPK28 or the S318A site directed mutant as indicated in the figures, and 10 μM peptide substrate. Reactions were initiated by addition of an ATP/[γ-^32^P]ATP mixture. Reactions contained combinations of CaCl_2_, or EGTA as indicated in the appropriate figures. For experiments at physiological Ca^2+^, final free Ca^2+^ concentrations were achieved by buffering CaCl_2_ with EGTA, calculated using the online WEBMAXC Extended calculator (102). Final reaction volumes were 40 μl. After addition of the ATP/[γ-^32^P]ATP mixture, reactions were allowed to proceed for 10 min at room temperature and were stopped by spotting 35 μl of each reaction onto P81 phosphocellulose cation exchange paper followed by washing three times for 5 min each in 0.45% (v/v) *o-*phosphoric acid. Incorporation of ^32^P was assessed by liquid scintillation counting.

### Intrinsic Trp fluorescence measurements

Intrinsic Trp fluorescence of recombinant purified dephosphorylated or hyperphosphorylated His_6_-CPK28 or His_6_-CPK28^S318A^ was performed in a PTI QuantaMaster steady-state spectrofluorimeter (Horiba Scientific) in a quartz cuvette with a 1 cm pathlength (Hellma USA Inc.). Data acquisition and background subtraction were performed using the FeliXGX software package (Horiba Scientific). Measurements were carried out on samples of CPK28 at a concentration of 200 nM in a buffer containing 20 mM HEPES-NaOH pH 7.2, 100 mM KCl, and 1 mM DTT. Trp fluorescence was measured at background Ca^2+^ levels (‘non-treated’) and after the titration of 100 µM CaCl_2_ into the same sample. Samples were measured at an excitation wavelength of 288 nm and fluorescence was collected between 300-450 nm in 1 nm steps with a dwell time of 1 second at each step. Each curve represents an average of two replicate scans of the same sample after appropriate background subtraction of buffer alone or buffer titrated with 100 µM CaCl_2_. Experiments were performed twice on independent preparations of all recombinant proteins.

### Phylogenetic analysis

To determine the conservation of Ser318 in group IV CDPKs, the full-length AtCPK28 protein sequence was used as a query in the Phytozome 12 BLAST tool, which identified a total of 114 amino acid sequences from 53 species (File S1). Similarly, the full-length sequences of all group I, II, and III AtCPKs were queried in Phytozome 12 BLAST limited to twelve species spanning the plant lineage (*M. polymorpha, P. patens, S. fallax, S. moellendorffi, A. trichopoda, O. sativa, A. thaliana, V. vinifera, R. comunis, B. rapa, T. cacao,* and *M. truncatula*) and a total of 327 amino acid sequences were retrieved (File S2). FASTA sequences were aligned using MUSCLE in MEGAX and the 11-amino acid window spanning Ser318 was extracted for visualization of conservation using WebLogo (103).

### Protein modeling

CPK28 was modeled using PHYRE2.0 Protein Fold Recognition Server (104) on Intensive Mode and visualized using PyMol Molecular Graphics System Version 2.4.0. The crystal structure of inactive TgCDPK1 (PDB:3KU2) was used as a template.

### Statistical analysis

Statistical significance was determined by a Student’s T-test or one-way ANOVA followed by Tukey’s post hoc test using GraphPad Prism version 8, as indicated.

## ACKNOWLEDGMENTS

We gratefully acknowledge Ray Zielinski for thoughtful guidance on aspects of this work related to Ca^2+^ signaling and for assistance with intrinsic fluorescence experiments. We also extend heartfelt thanks to Cyril Zipfel (The Sainsbury Laboratory and the University of Zurich) for critical discussions regarding early observations of the transphosphorylation between BIK1 and CPK28, and for critically reading this work prior to submission. We thank Tina Romeis (Halle) for insightful discussions and sharing published materials (Table S1), as well as Tony Papanicolaou, Ruxandra Bogdan and Andrew Ji (Queen’s University) for their technical assistance, and Darrel Desveaux (University of Toronto) for providing *Pseudomonas syringae* pv. *tomato* DC3000. We additionally thank all members of the Monaghan Laboratory for their helpful and critical feedback of this work.

## SUPPLEMENTARY DATA

**Figure S1.**
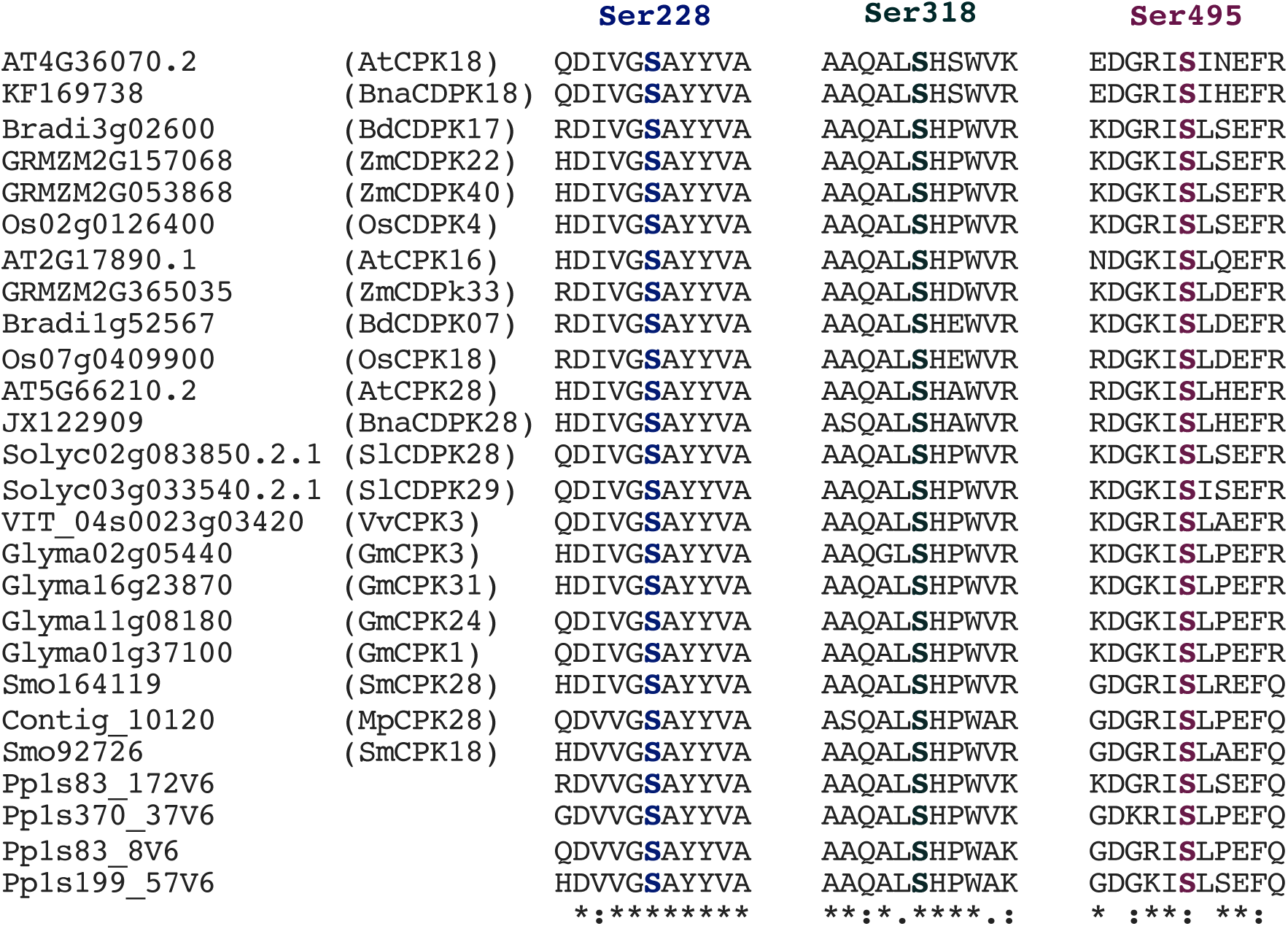
Ser228, Ser318 and Ser495 are conserved across group IV CDPKs. Amino acid sequences of representative group IV CDPKs from eudicots (*Arabidopsis thaliana, Glycine max*, *Solanum lyopersicum*, *Vitis vinifera* and *Brassica napus)*, monocots *(Brachypodium distachyon*, *Zea mays*, and *Oryza sativa*), bryophytes (*Psychomiteralla patens)*, liverworts (*Marchantia polymorpha*), and pteridophytes (*Selaginella moellendorffii)* were aligned using Clustal Omega Multiple Sequence Alignment Tool and the residues corresponding to positions 228, 318, and 495 of *Arabidopsis thaliana* CPK28 were compared. “*”=perfect alignment; “:”=strong similarity; “.”=weak similarity.

**Figure S2.**
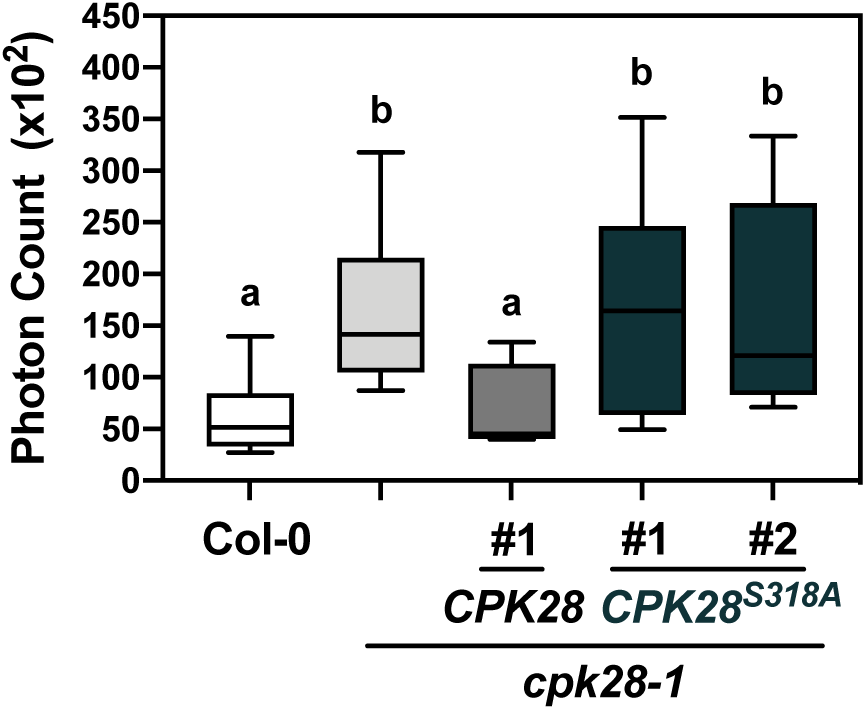
CPK28^S318A^ does not complement *cpk28-1* in response to elf18 treatment. Oxidative bursts of Col-0, *cpk28-1*, *cpk28-1/pCPK28:CPK28-FLAG*, and *cpk28-1/pCPK28:CPK28^S318A^-FLAG* lines following continual growth in elf18 (100 nM) for 12 days (n=6). Values are presented as boxplots indicating first and third quartiles, split by a median line, and whiskers representing maximum and minimum values. Statistically different groups (p<0.005) are indicated with lowercase letters, as determined by ANOVA analysis followed by Tukey’s posthoc test. Experiments were conducted three times with similar results.

**Figure S3.**
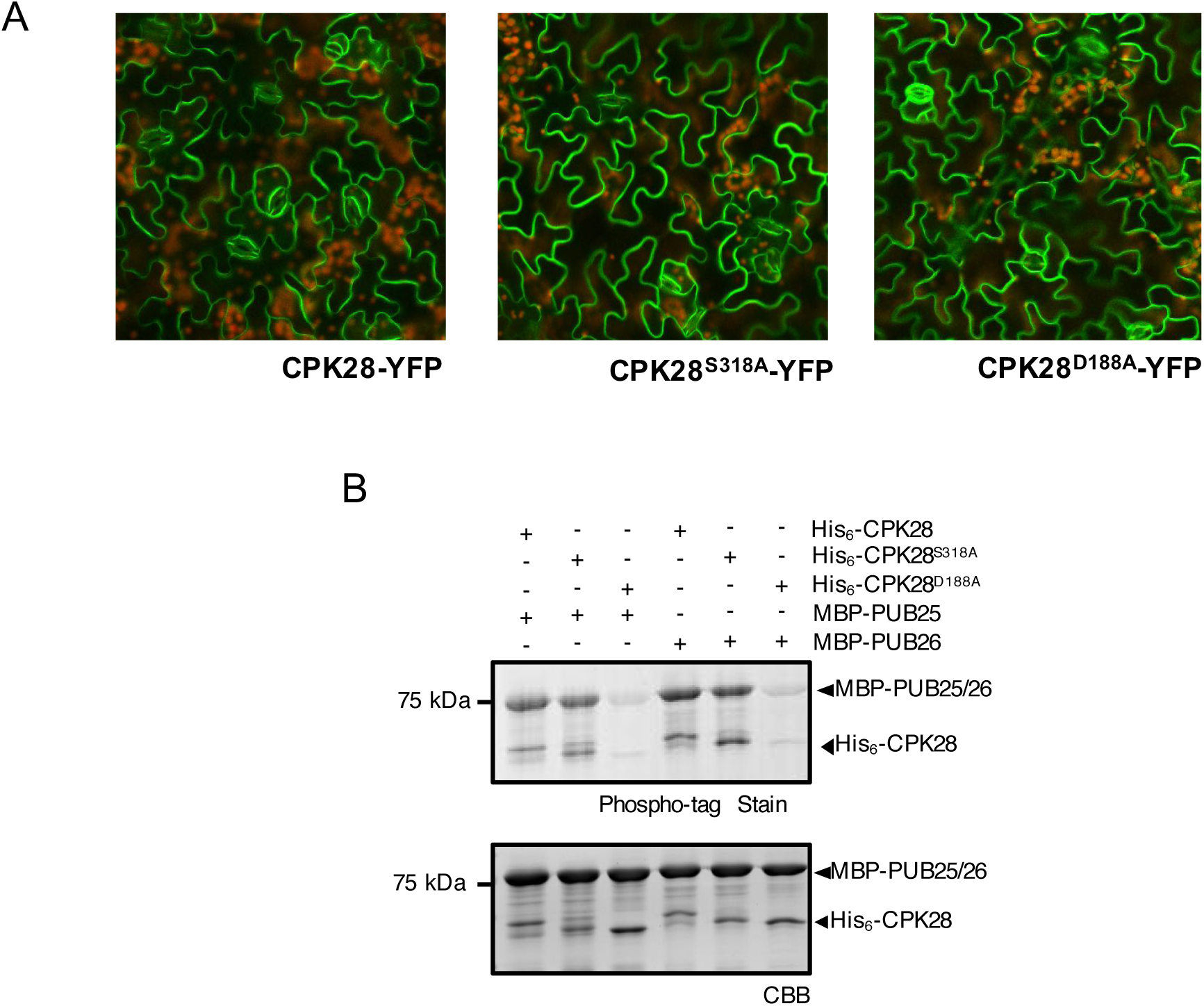
CPK28^S318A^ localizes to the plasma membrane and phosphorylates PUB25 and PUB26. *(A)* Subcellular localization of CPK28-YFP, CPK28^S318A^-YFP, and CPK28^D188A^-YFP stably expressed under the CaMV *35S* promoter in *cpk28-1* mutants. Imaging was performed using a LSM 710 (Zeiss) confocal microscope with excitation at 488 nm for yellow fluorescent protein (YFP; coloured green) and a range of 510-540 nm for measuring emission. Chlorophyll autofluorescence (coloured red) was detected with an excitation wavelength of 543 nm and an emission wavelength range of 680-760 nm. *(B)* Phospho-tag gel stain of *in vitro* kinase assay using recombinantly produced His_6_-CPK28, His_6_-CPK28^S318A^, or His_6_-CPK28^D188A^ and MBP-PUB25 or MBP-PUB26. Gels were stained with Coomassie Brilliant Blue (CBB) to assess loading. Experiments were conducted at least three times with similar results.

**Figure S4.**
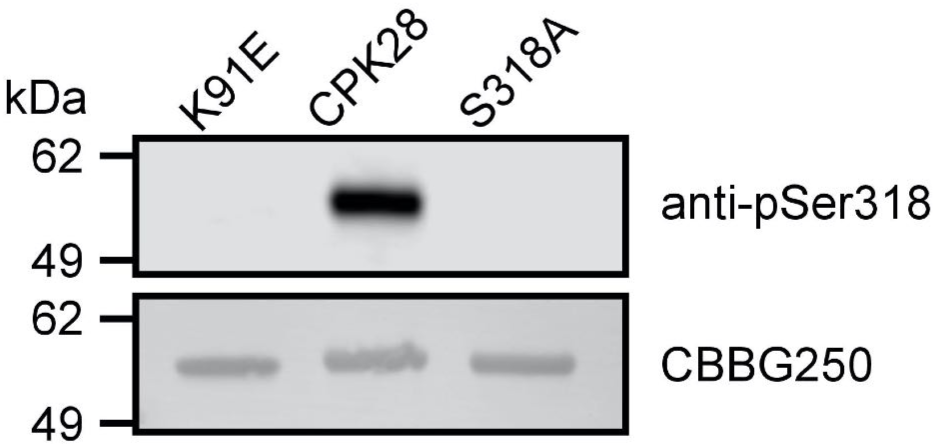
Specificity of the *α*-pSer318 antibody. Specificity of the CPK28 pSer318 antibody was determined by immunoblotting against wildtype (CPK28), kinase-dead (CPK28^K91E^), or CPK28^S318A^ . 200 ng of *in situ* phosphorylated purified recombinant protein was separated by gel electrophoresis and blotted to a PVDF membrane before probing with 2 µg/ml anti-CPK28 pSer318 IgGs. Only the wildtype protein was detected, demonstrating specificity of the antibody for the pSer318 site. Gels were stained with Coomassie Brilliant Blue (CBBG250) to assess loading. Experiments were conducted twice with similar results.

**Figure S5.**
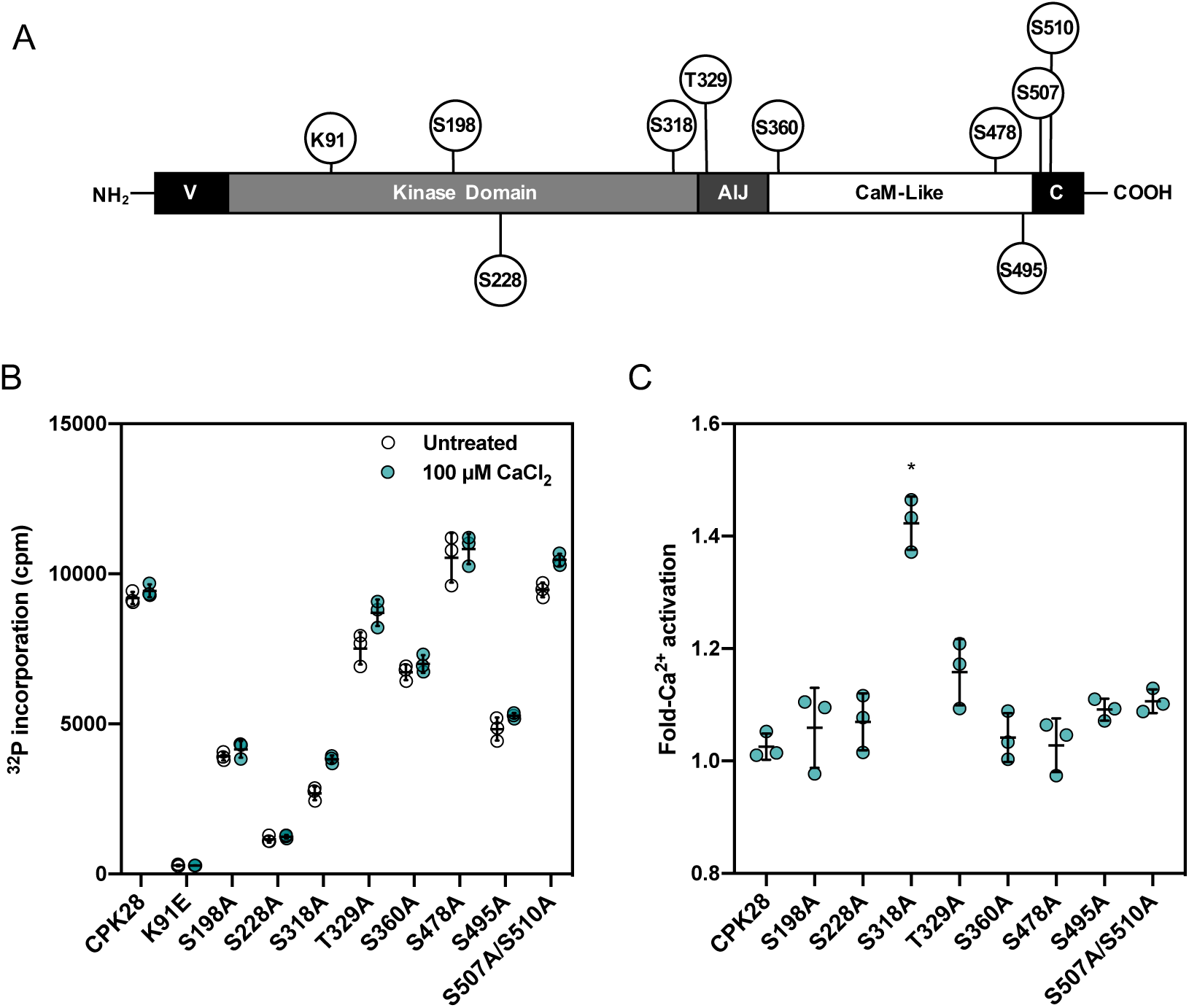
Ser318 phosphorylation uniquely primes CPK28 for Ca^2+^-activation. *(A)* Position of tested phosphorylation sites across CPK28 protein domains. *(B)* Biochemical screen of CPK28 phospho-null mutants for Ca^2+^ activation of peptide kinase activity using the ACSM+1 peptide as substrate. Activity was assessed using *in situ* phosphorylated purified recombinant proteins at either background (open circles) or 100 µM CaCl_2_ (teal circles) and is shown as ^32^P incorporation in cpm. Individual data points (three technical replicates) are shown with mean and standard deviation. *(C)* Fold-activation of CPK28 phospho-null mutants by the addition of excess Ca^2+^ derived from data shown in *(B)*. No difference is observed for wildtype CPK28 between these two conditions and only the S318A site-directed mutants showed statistically significant activation by Ca^2+^ (Kruskal-Wallis ANOVA, *p* = 0.024361, n = 3 technical replicates).The screen with all phospho-site mutants was performed once and S318A was selected for confirmation of altered calcium sensitivity.

**Figure S6.**
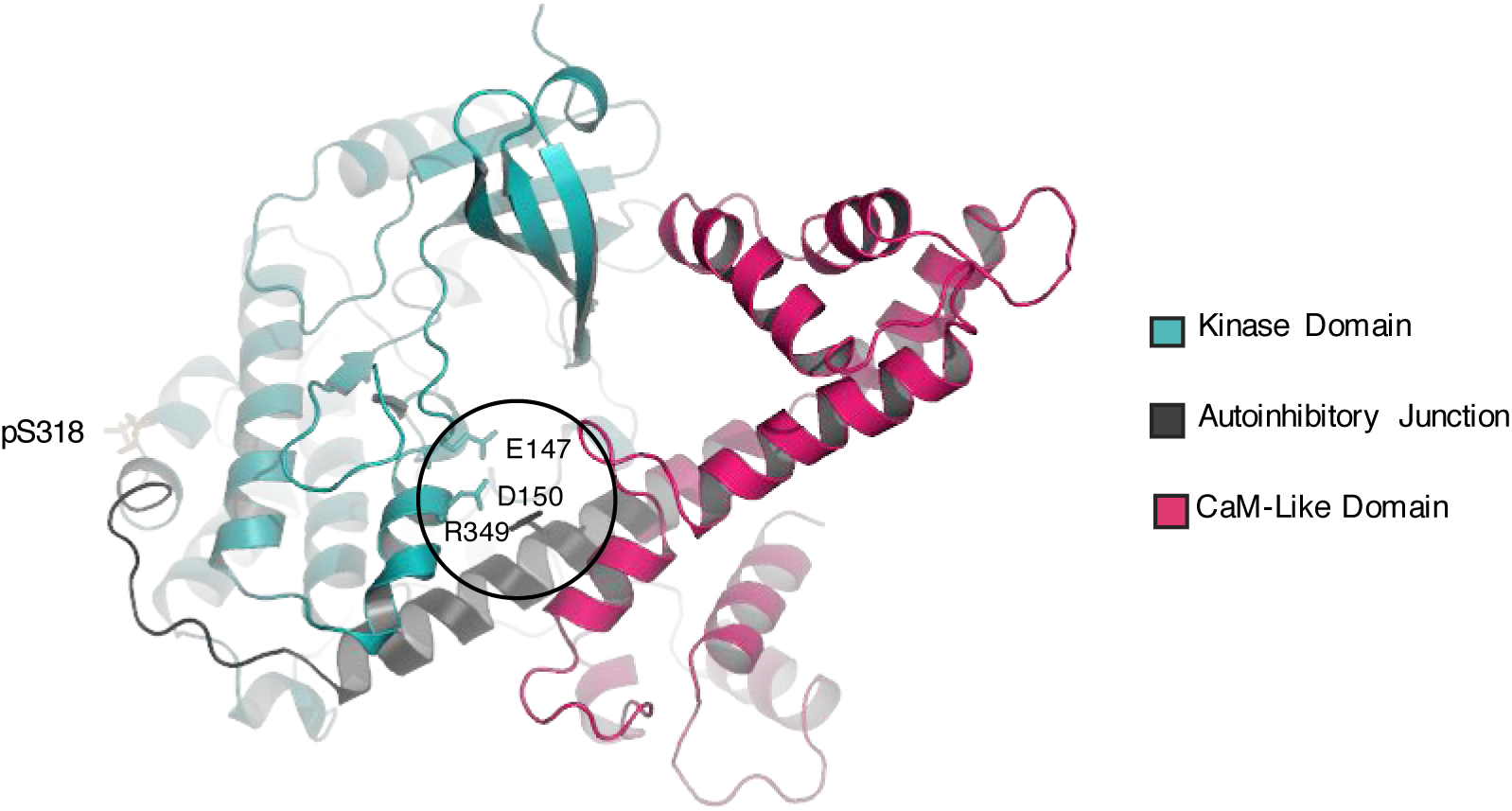
Structural modeling of CPK28. A model for CPK28 was generated using the PHYRE2.0 Protein Recognition Server (104) with inactive TgCDPK1 (iTgCDPK1; 3KU2) as a template (95% confidence score). Phosphorylated Ser318 is indicated in orange. The residues in the autoinhibitory triad that stabilizes TgCDPK1 (Lys338-Glu135-Asp138) (31), corresponding to Arg349-Glu147-Asp150 in CPK28, are labelled. CPK28 was visualized using the PyMOL Molecular Graphics System, Version 2.0 Schrödinger, LLC with PyTMs legacy add-on for post-translational modifications.

**Table S1.**
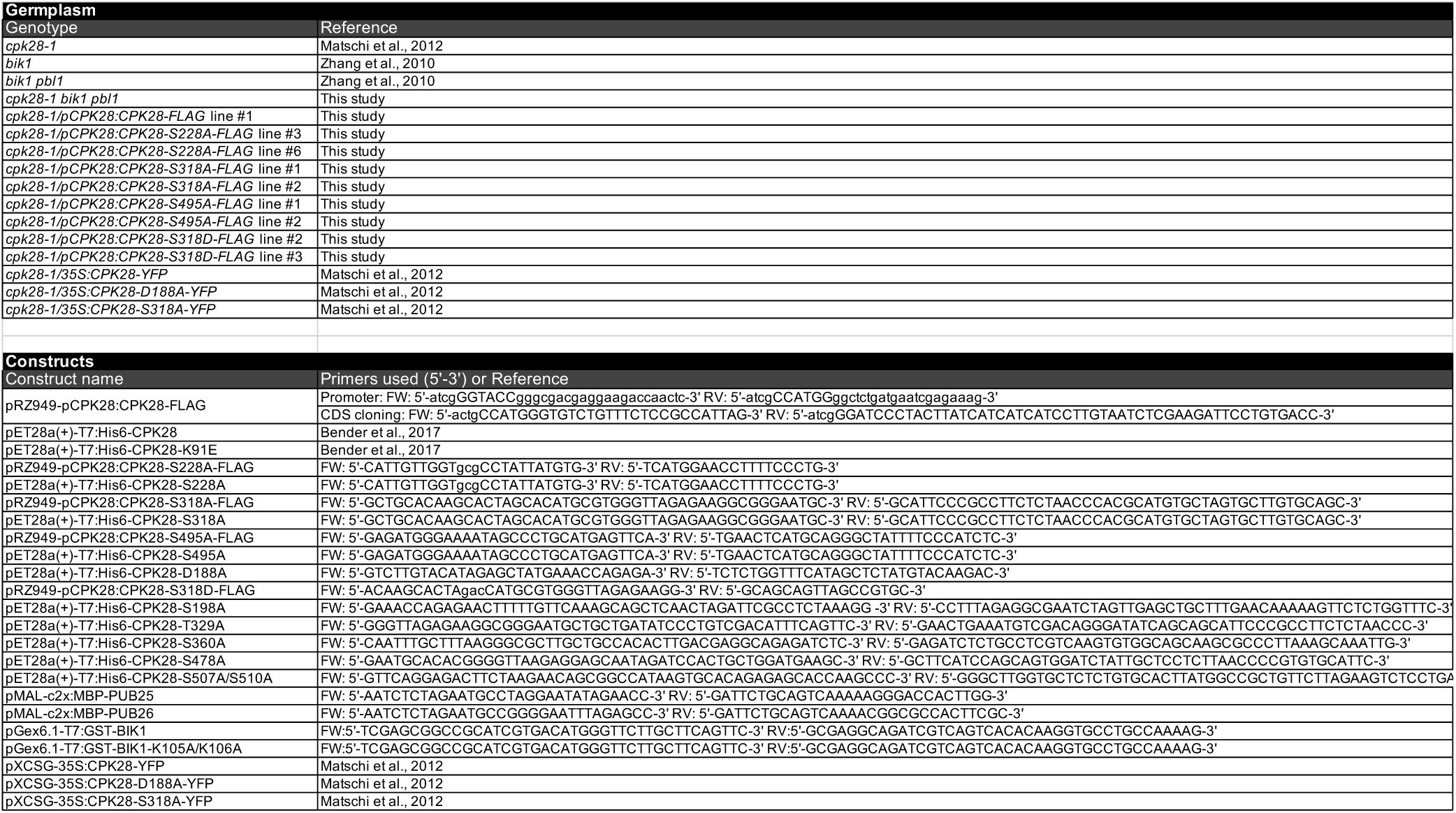
List of germplasm, constructs, and primers used in this study.

**File S1. Amino acid sequences of group IV CDPKs across the plant lineage.**

This text file includes all FASTA-formatted sequences retrieved following a query for group IV CDPKs using the Phytozome 12 BLAST tool (114 sequences from 53 species).

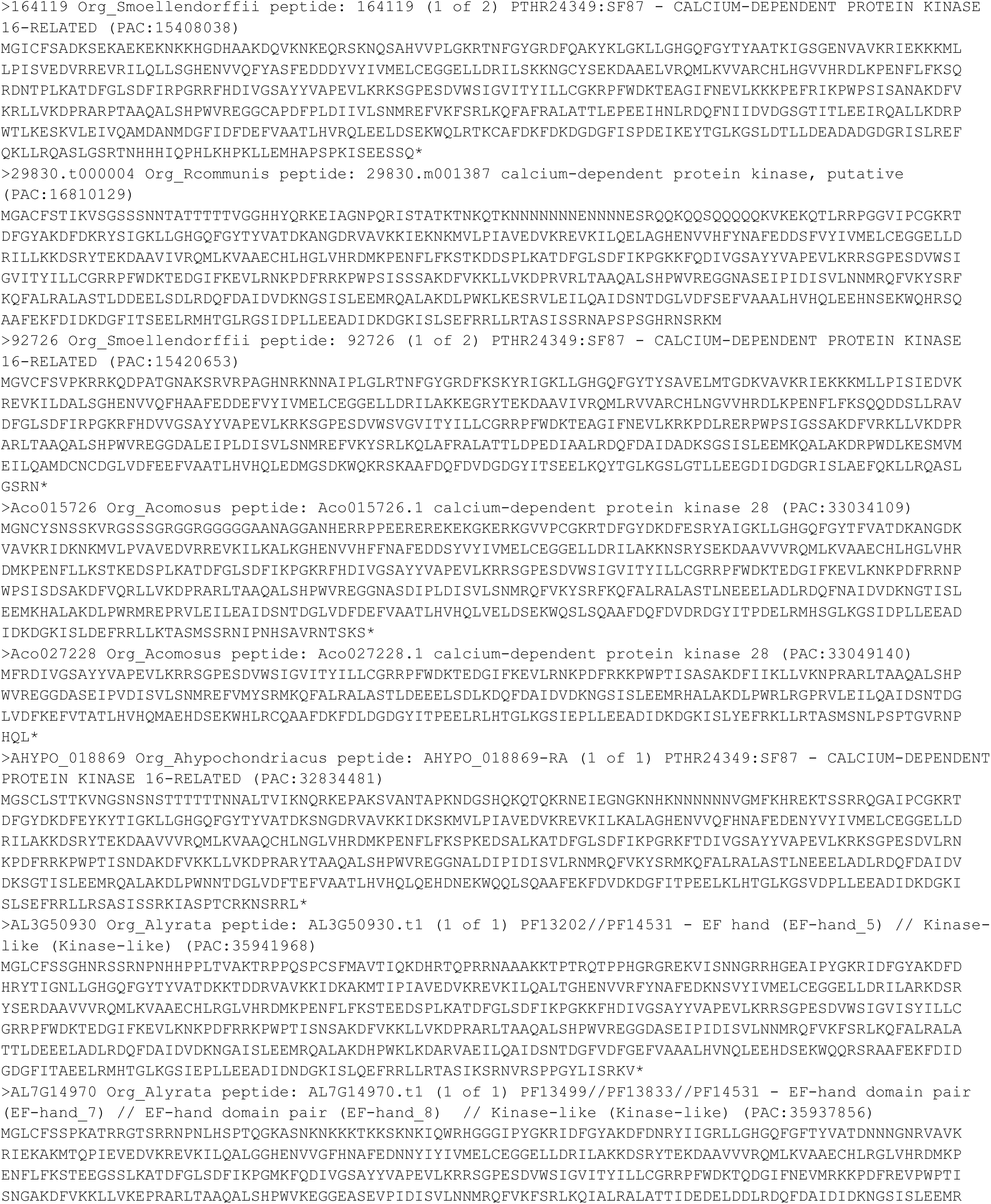
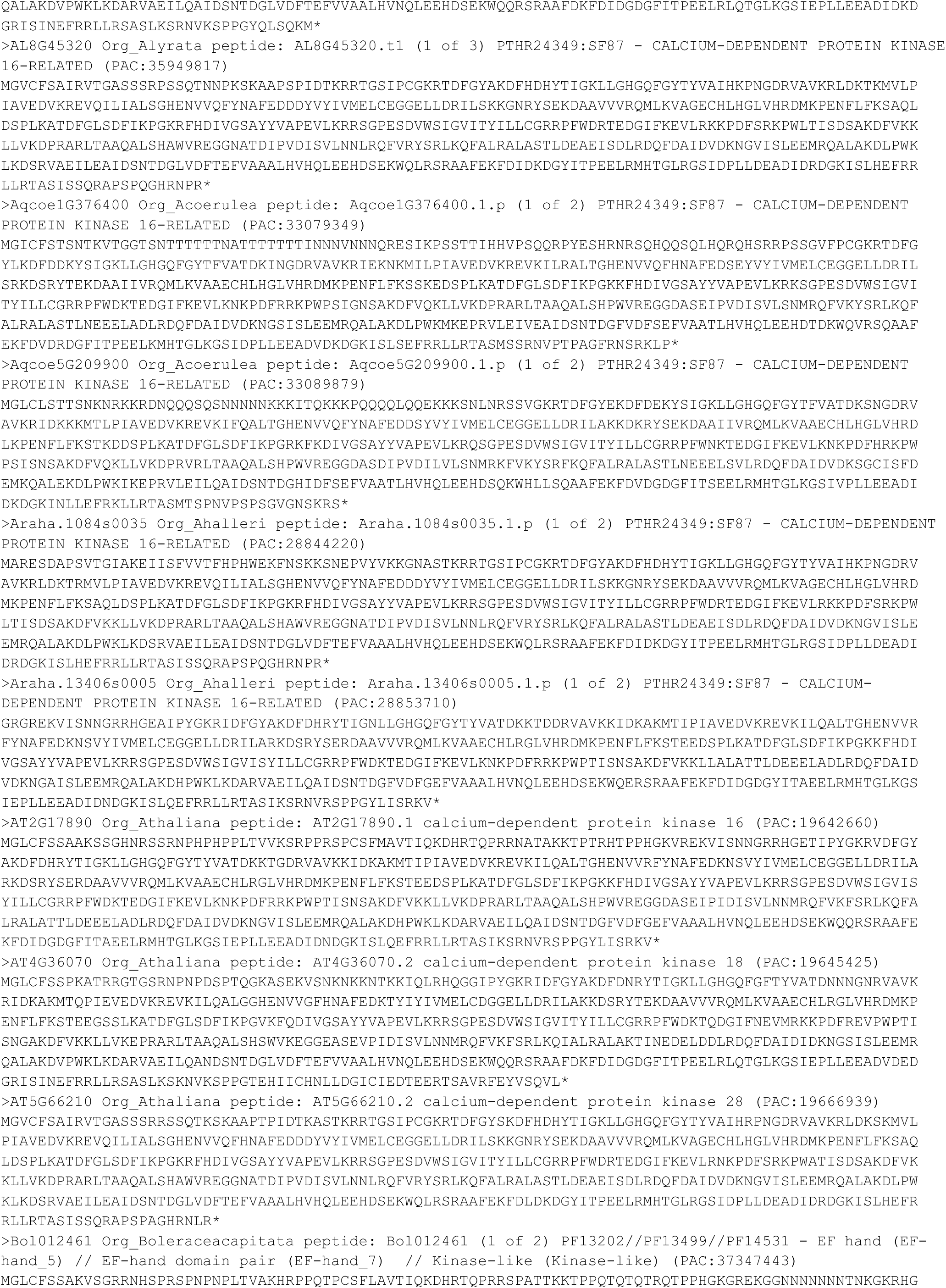
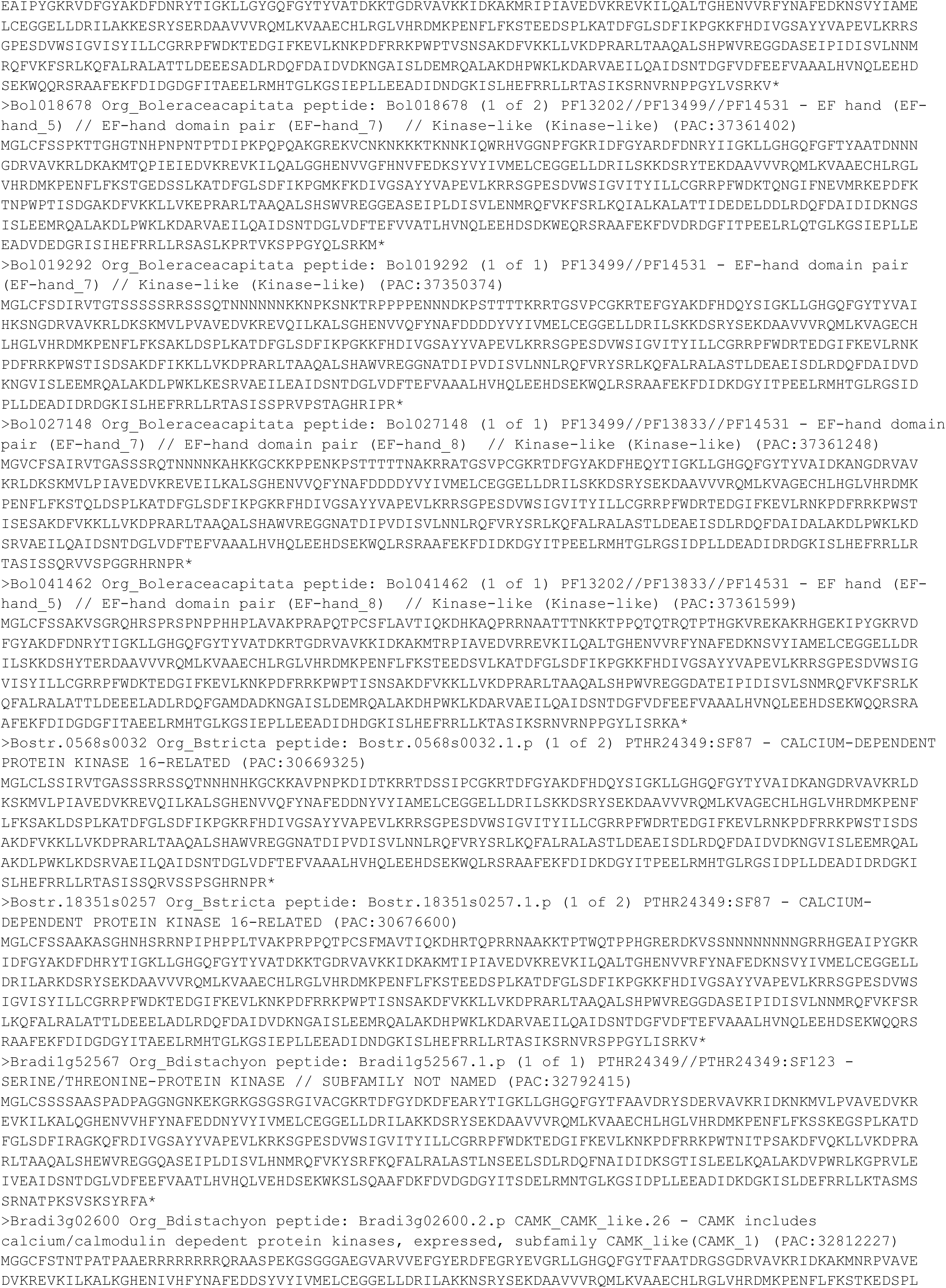
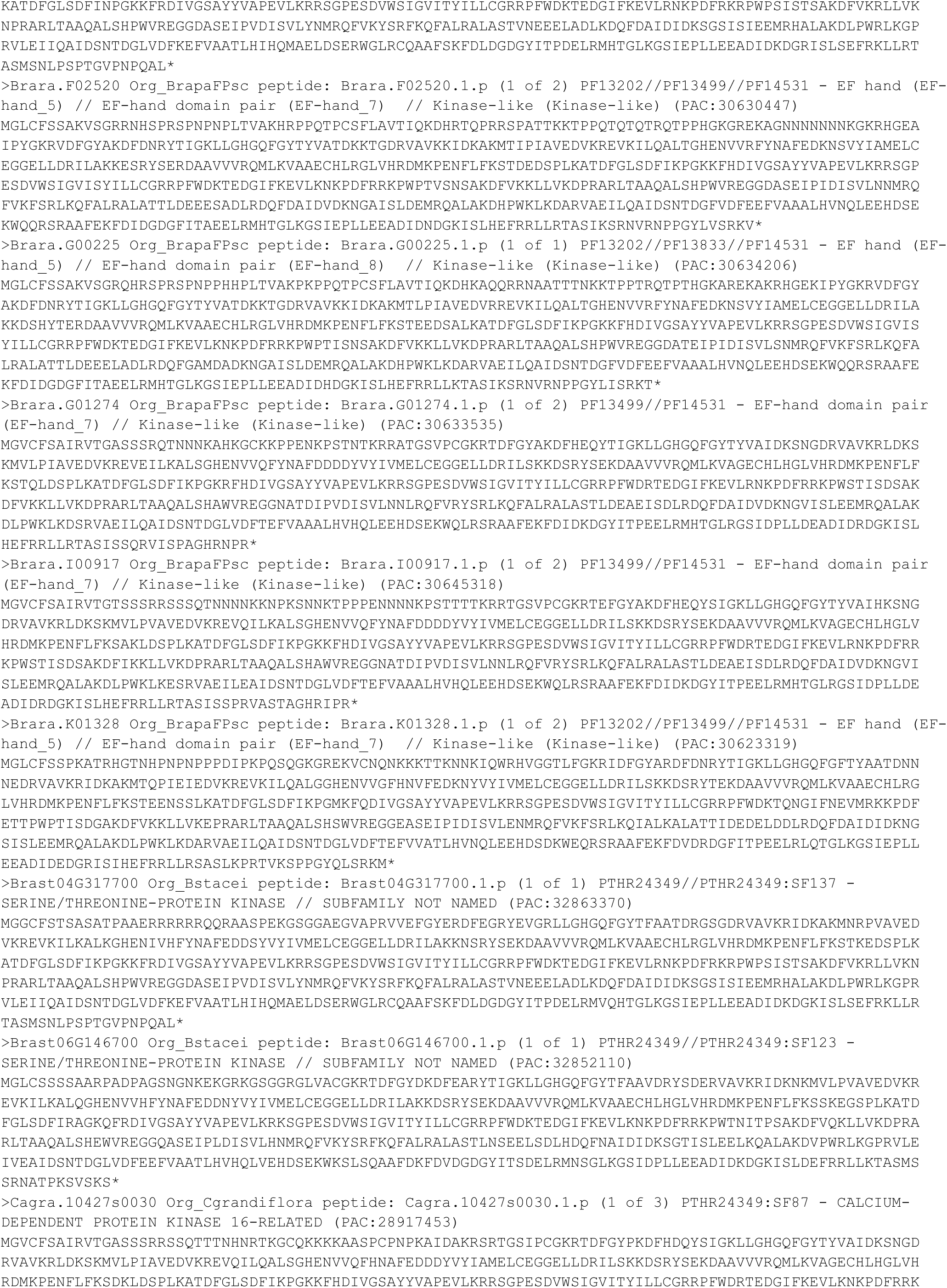
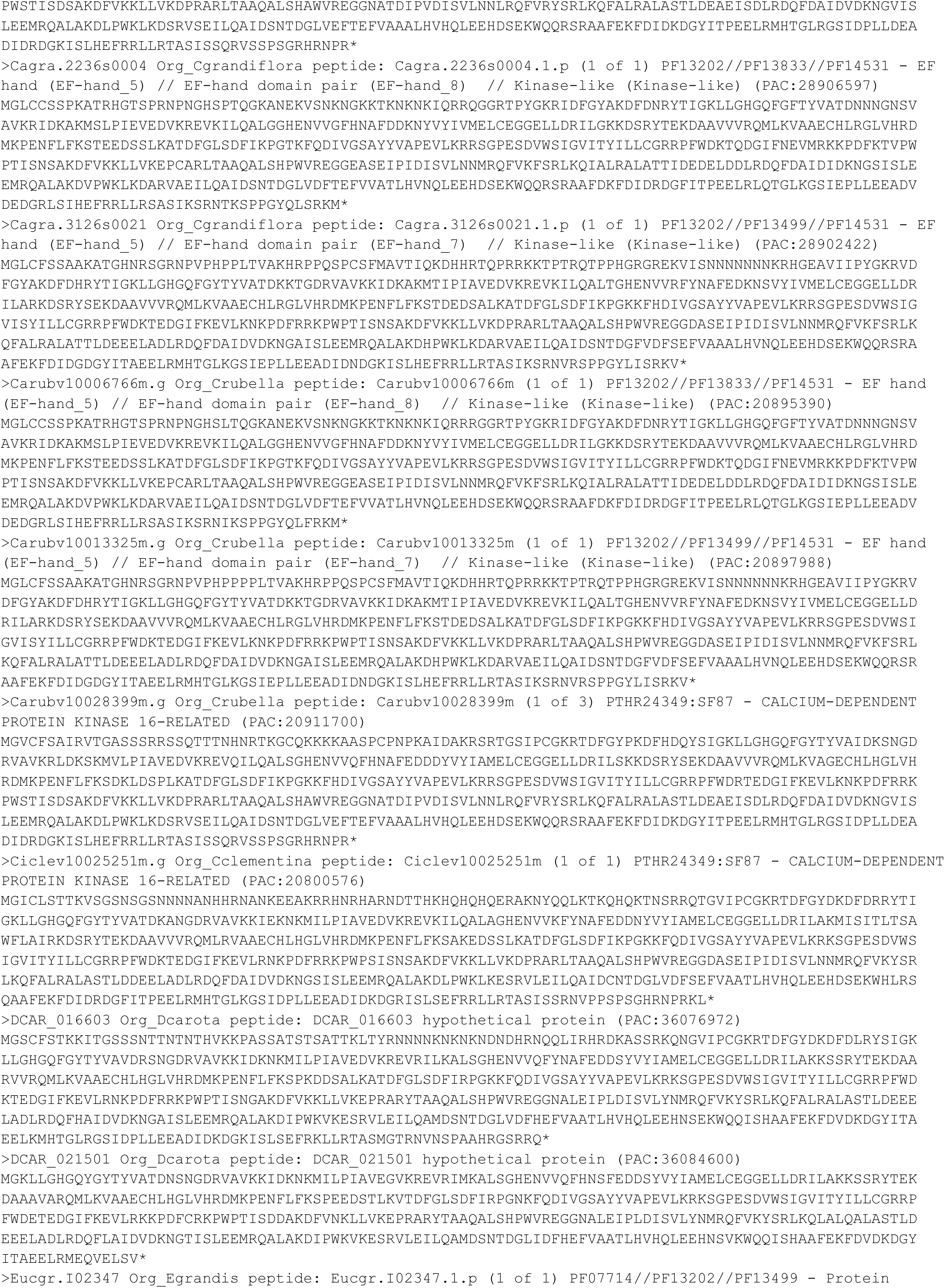
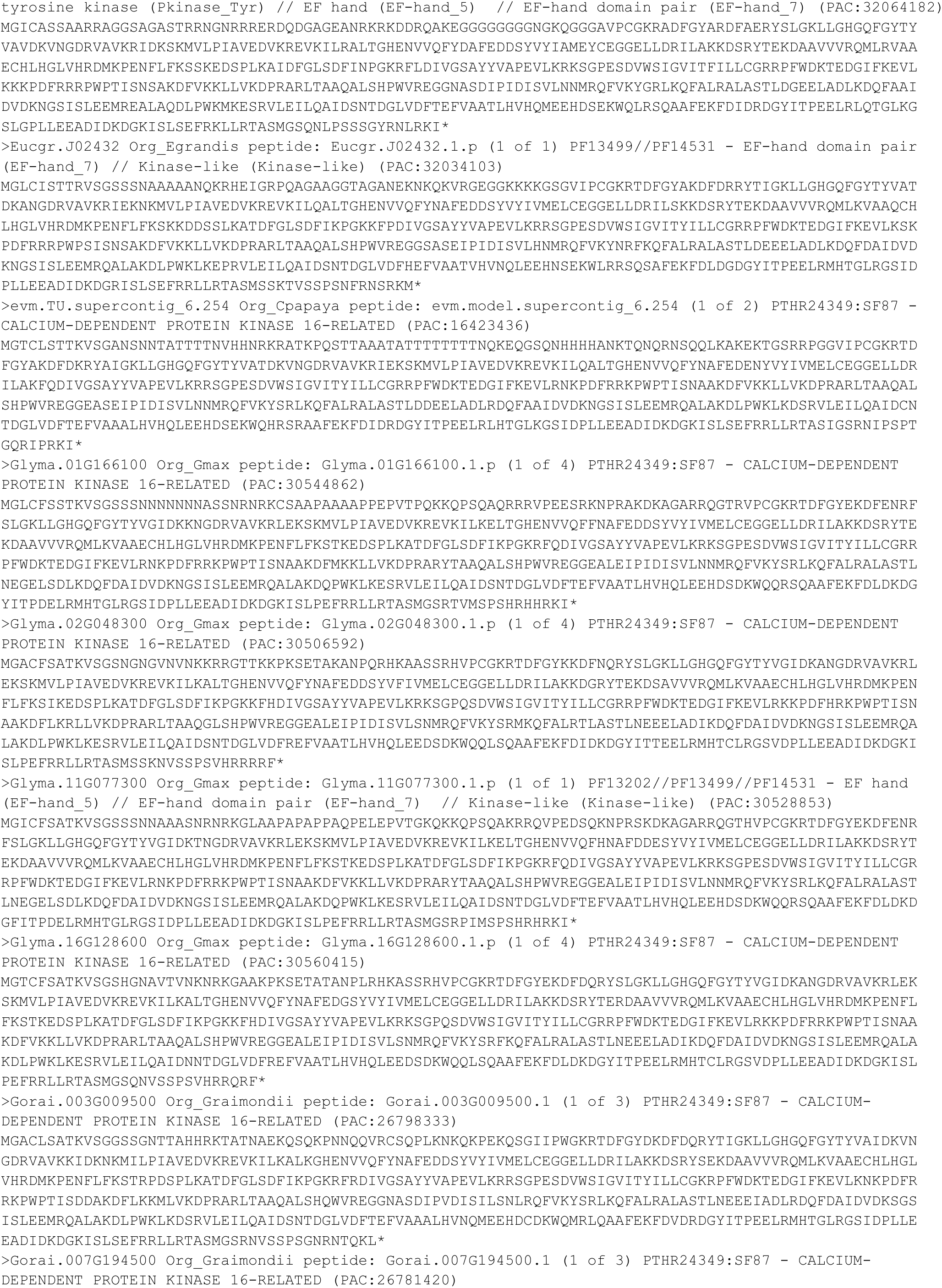
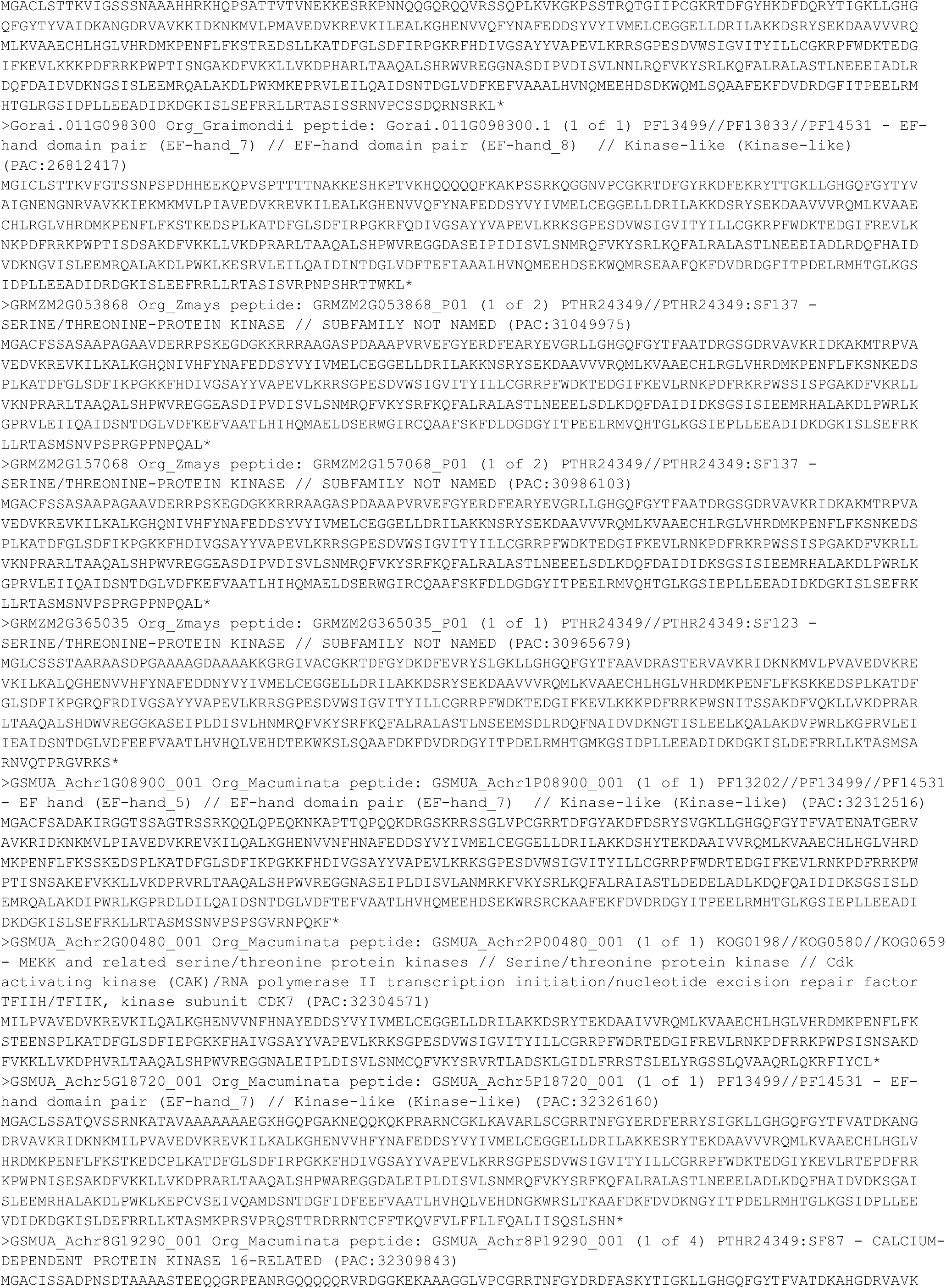
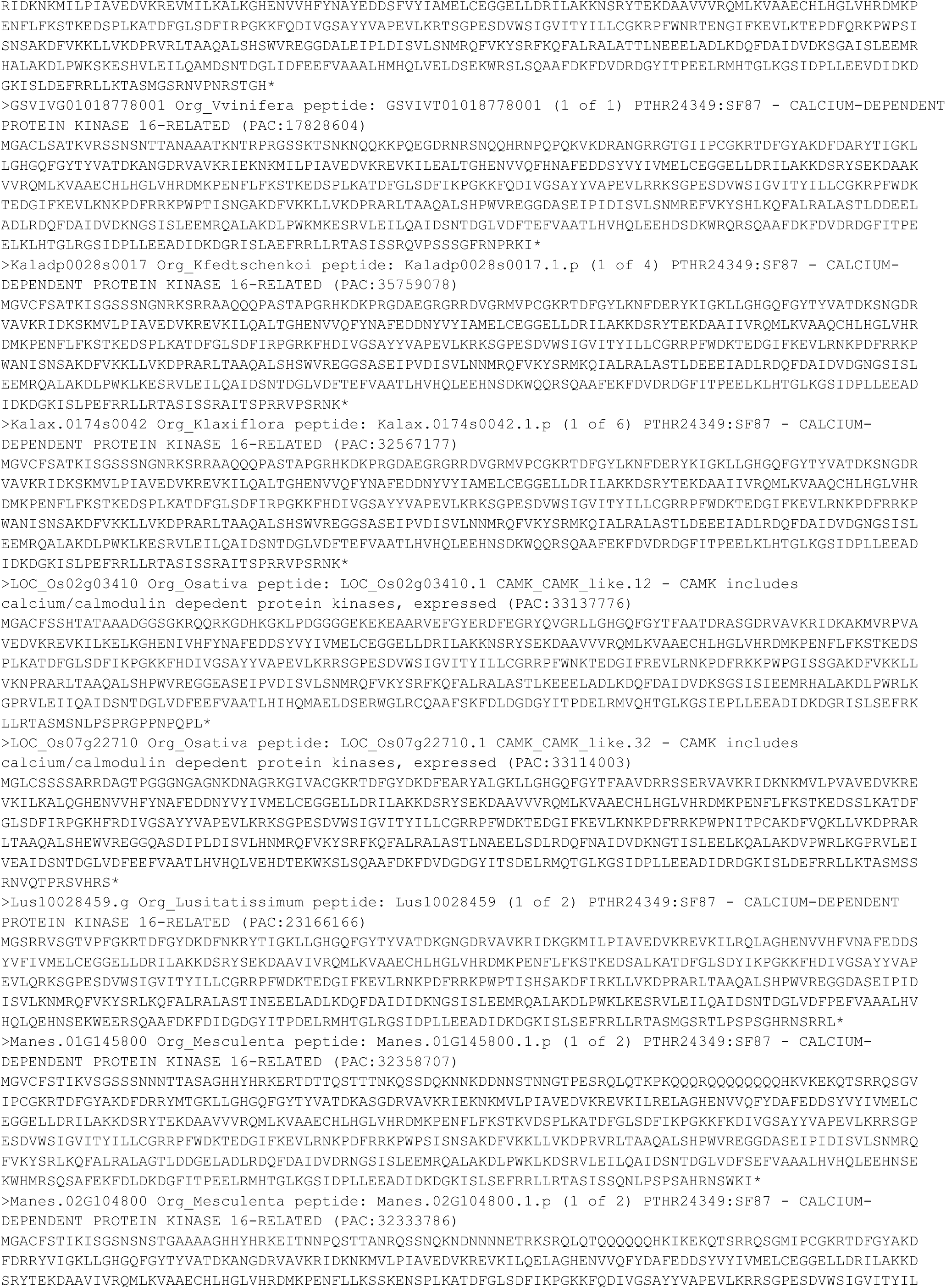
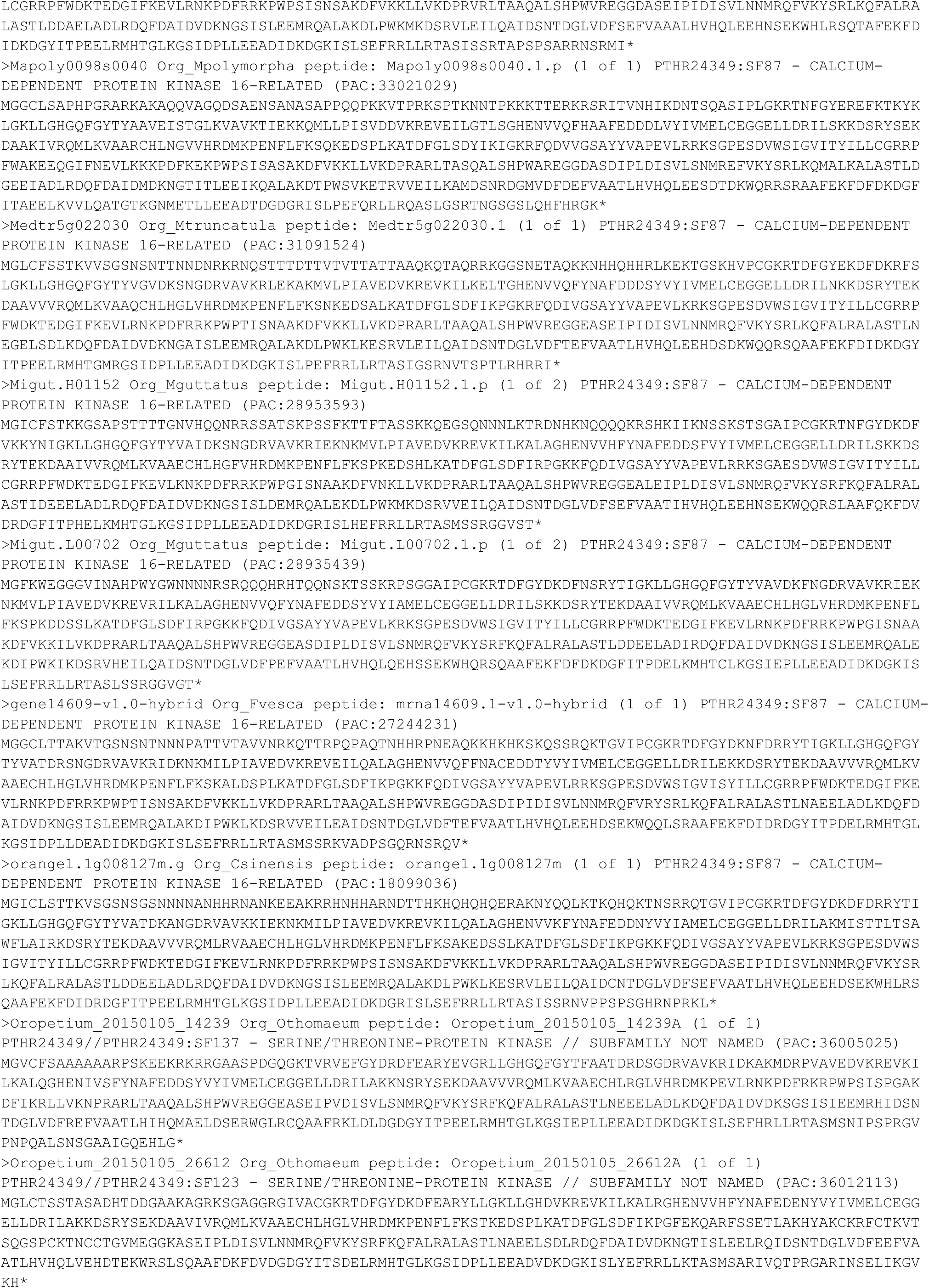
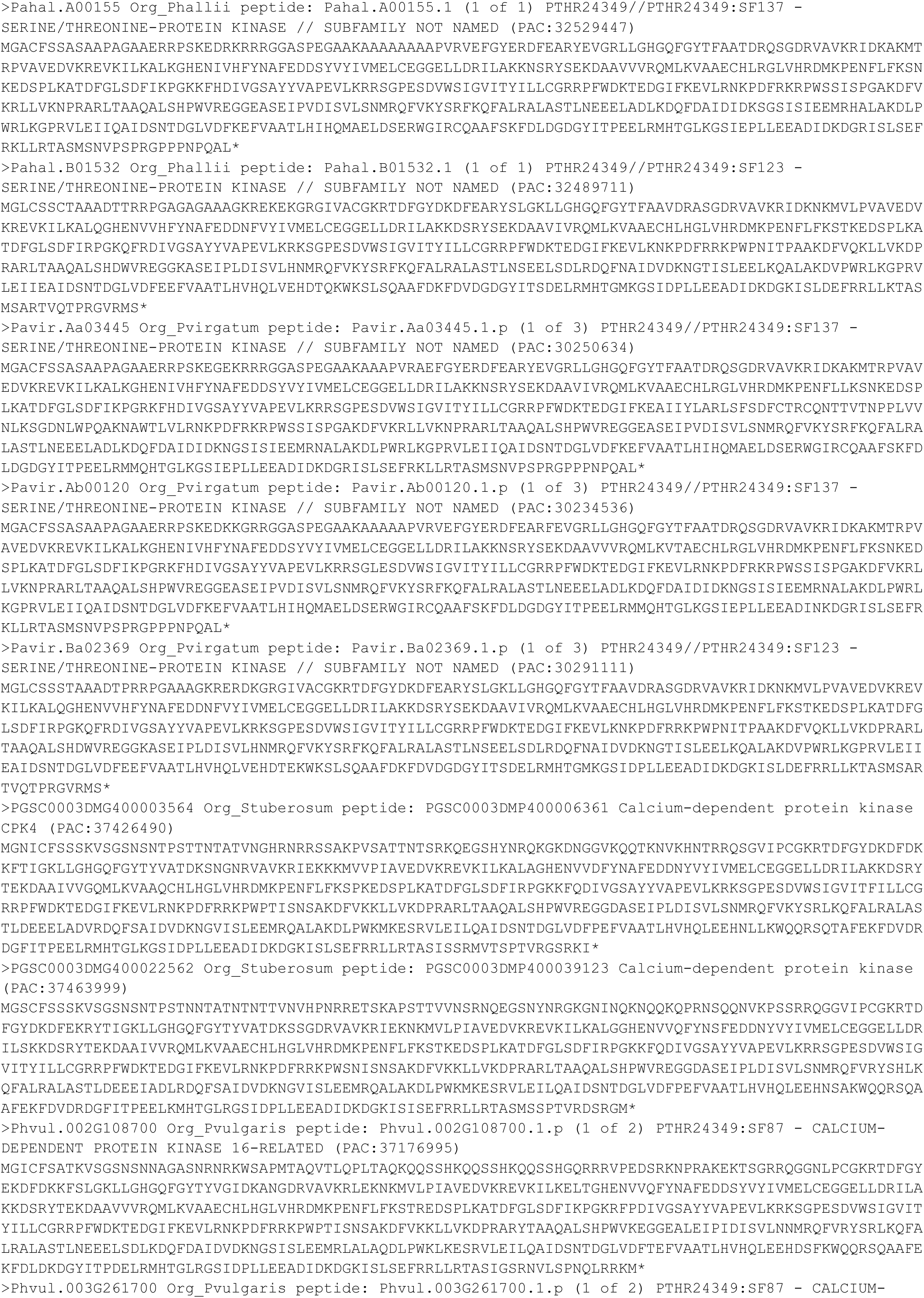
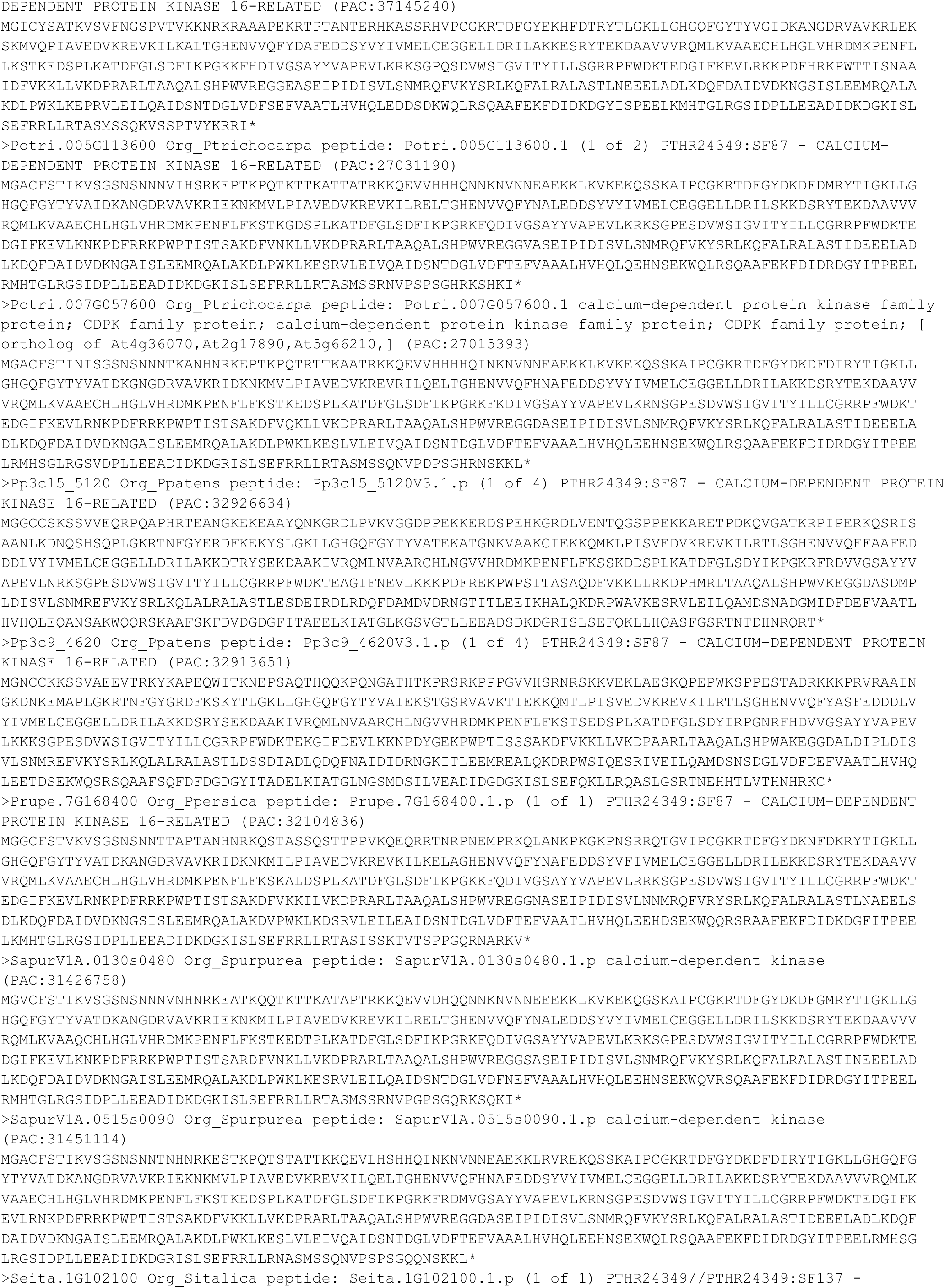
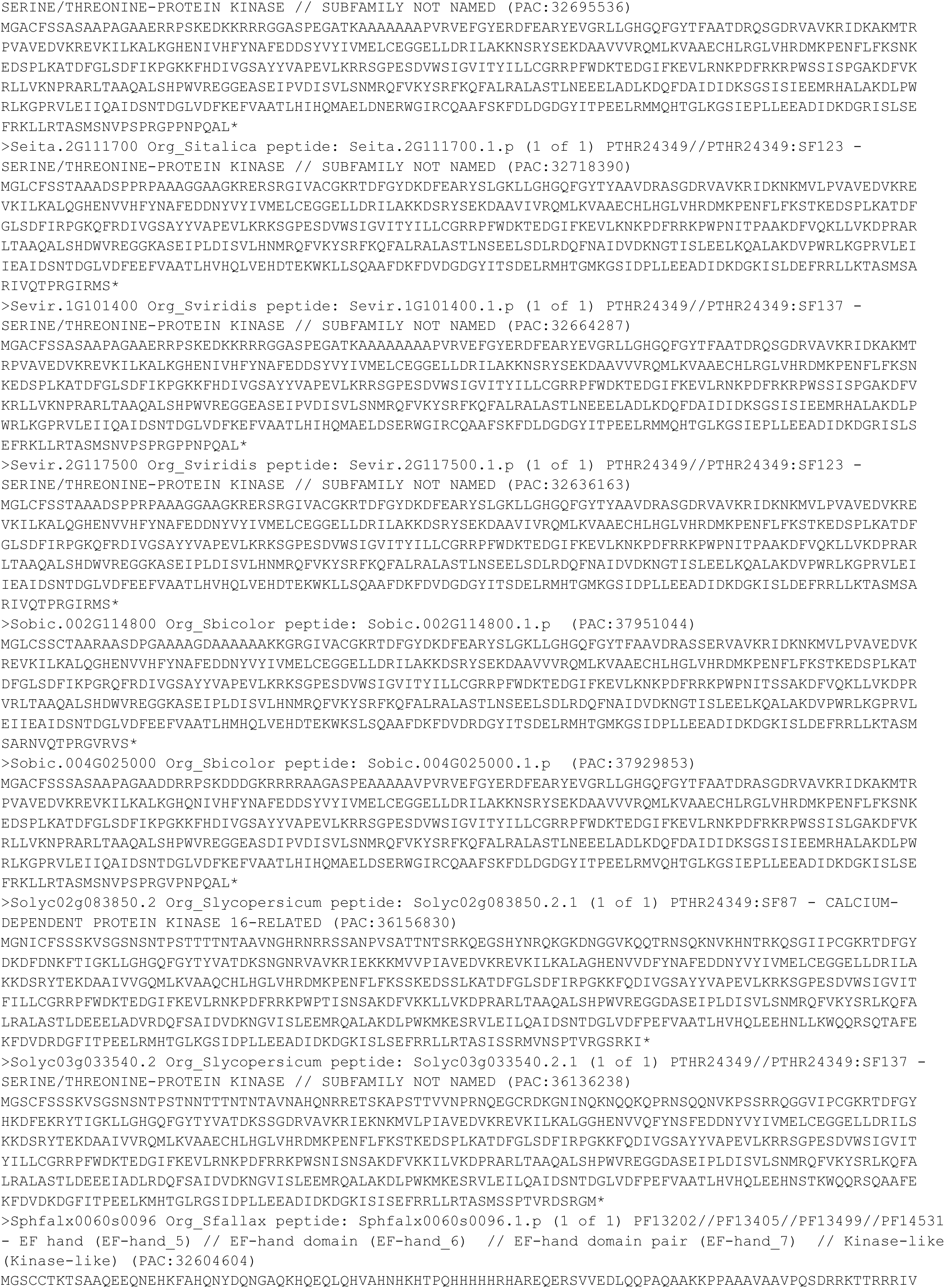
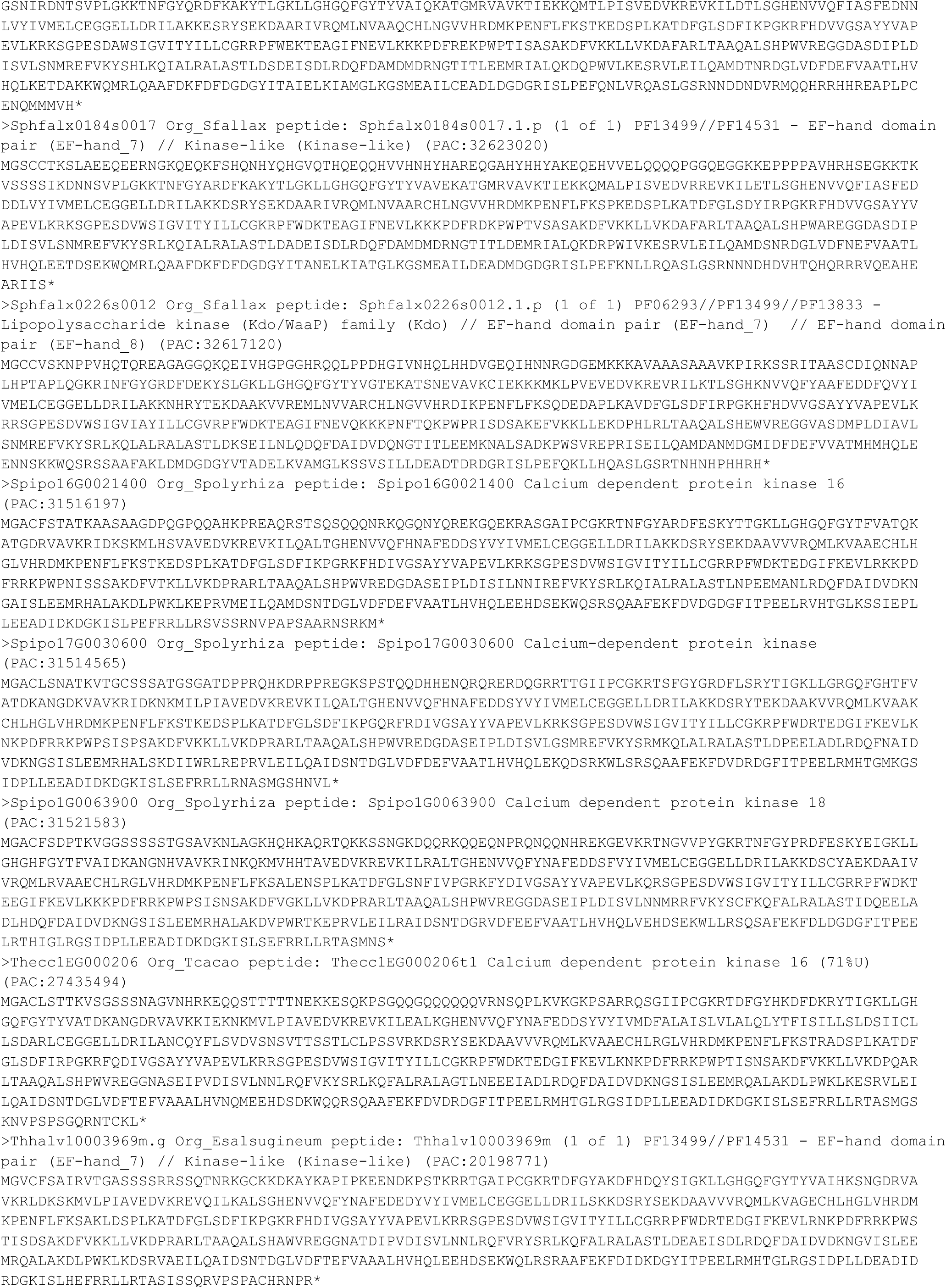
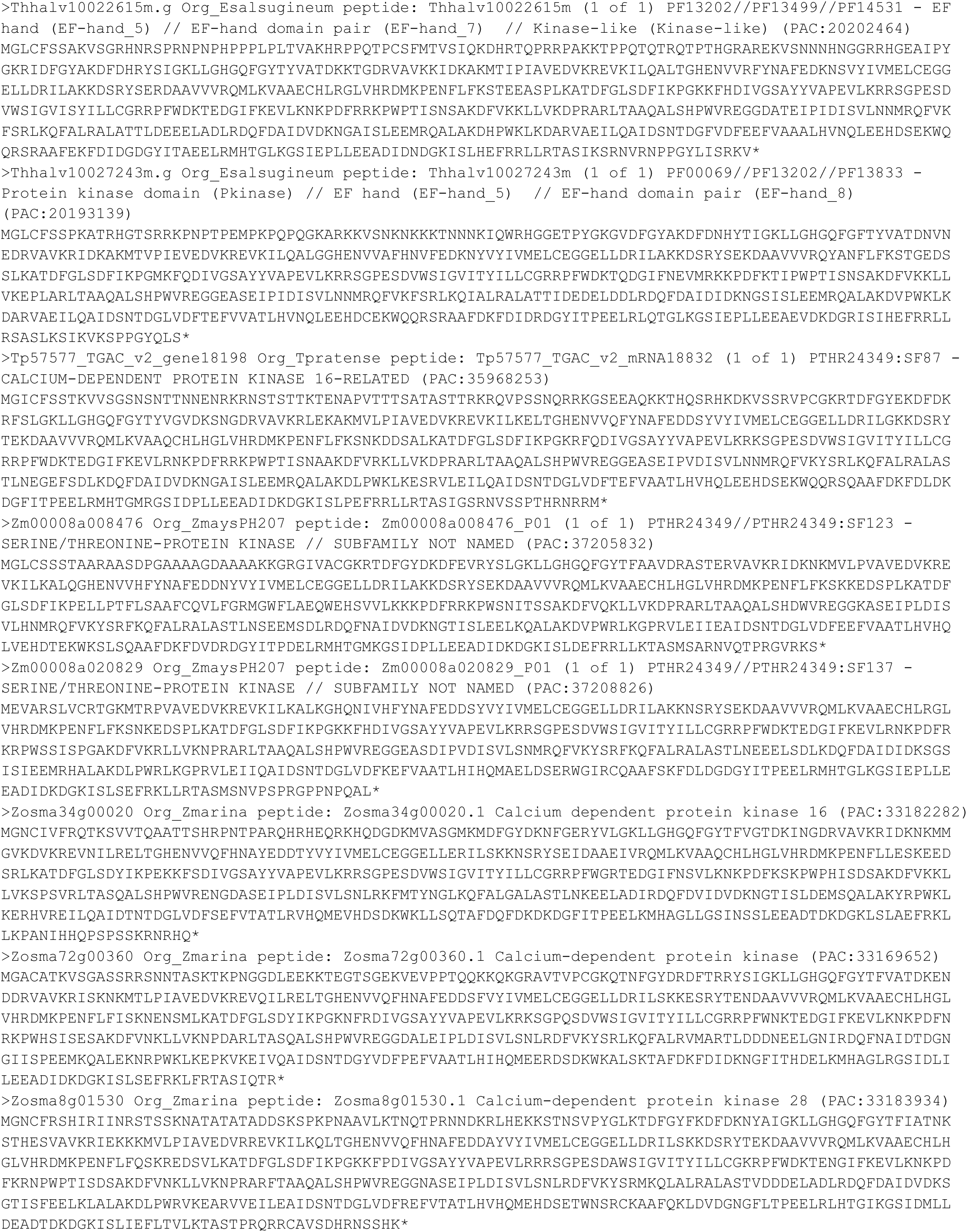

**File S2. Amino acid sequences of representative group I, II, and III CDPKs across the plant lineage.**

This text file includes 327 FASTA-formatted sequences retrieved from the Phytozome 12 BLAST tool following a query for group I, II, and III CDPKs in 12 species spanning the plant lineage (*M. polymorpha, P. patens, S. fallax, S. moellendorffi, A. trichopoda, O. sativa, A. thaliana, V. vinifera, R. comunis, B. rapa, T. cacao,* and *M. truncatula*).

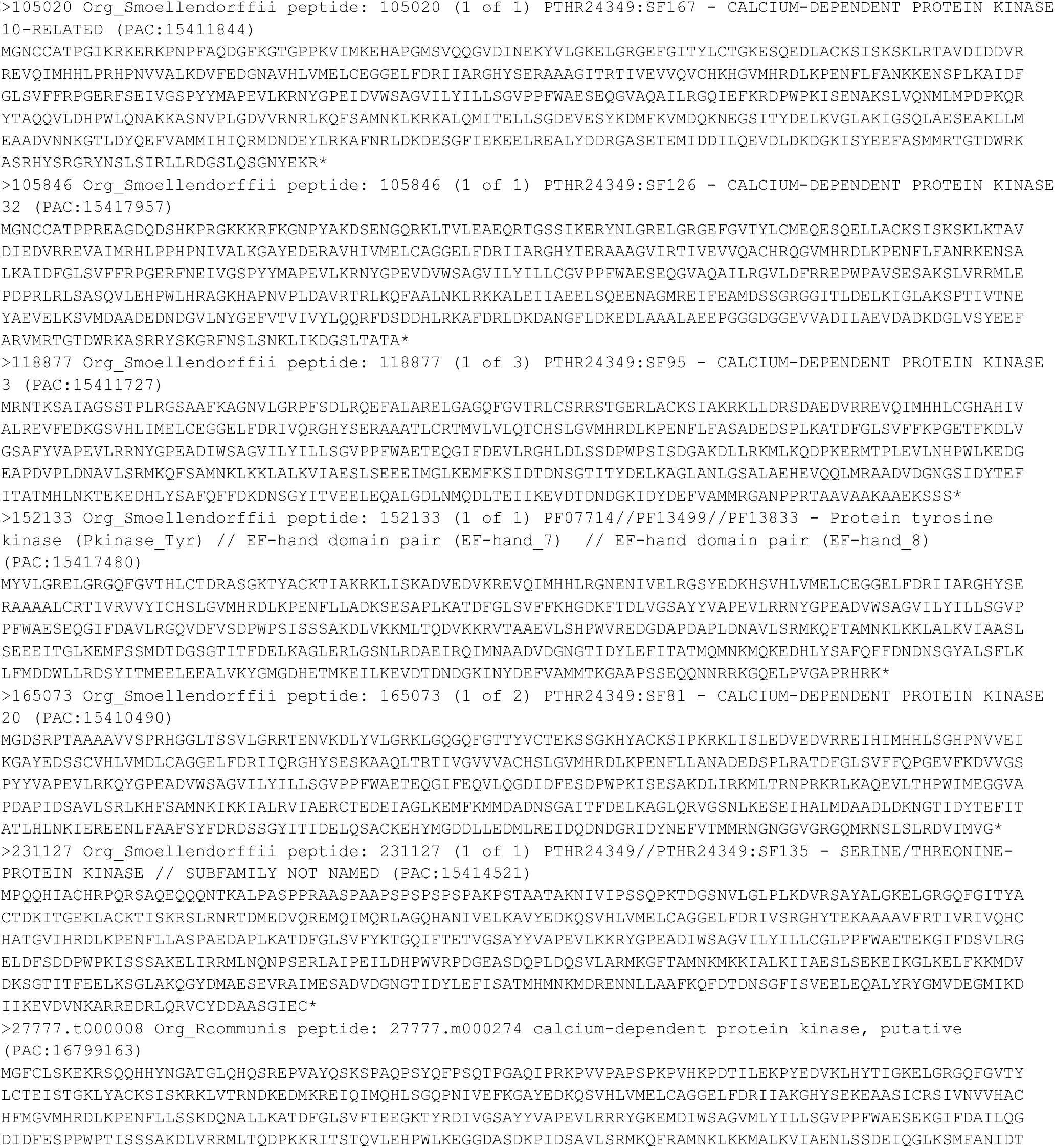
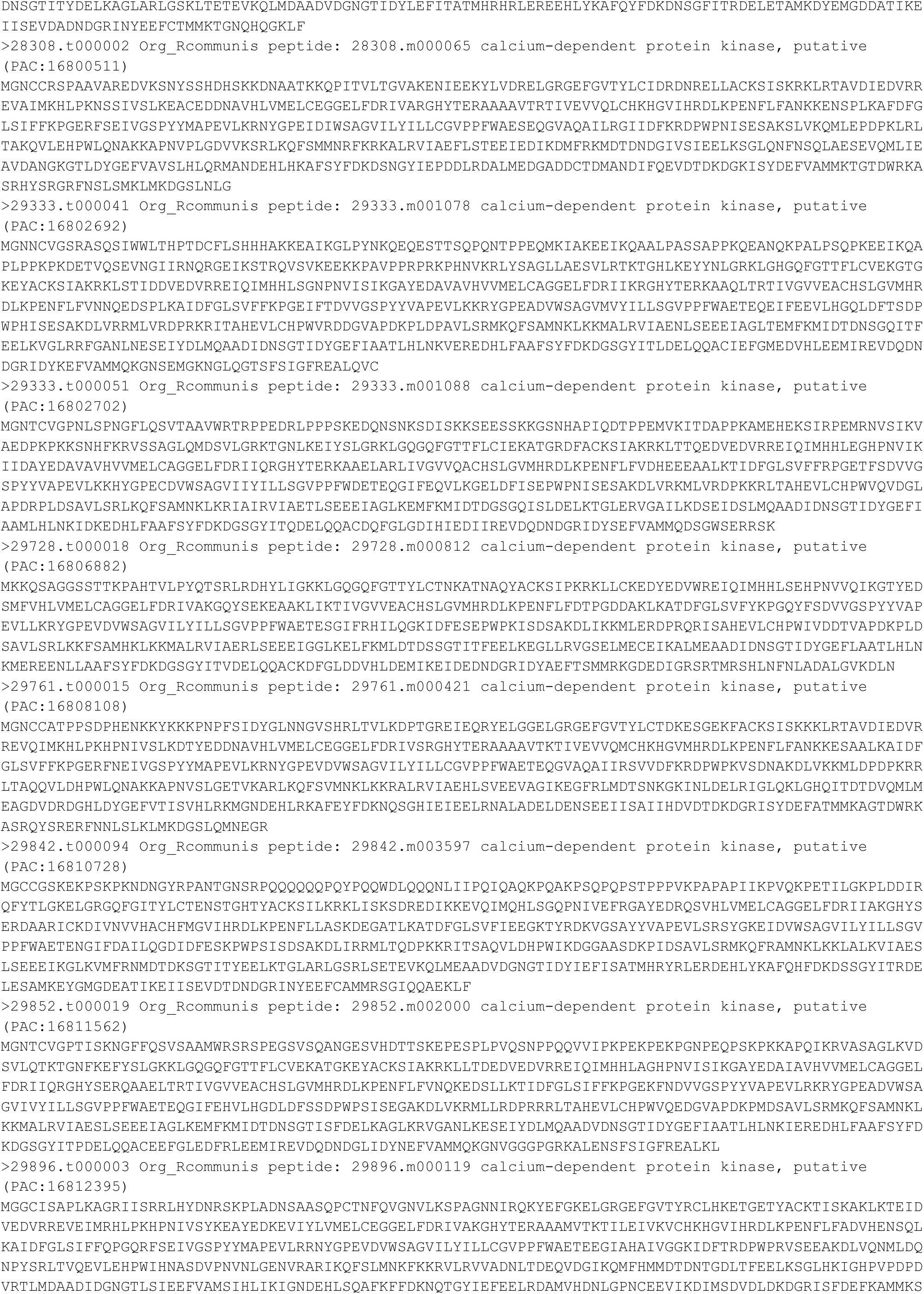
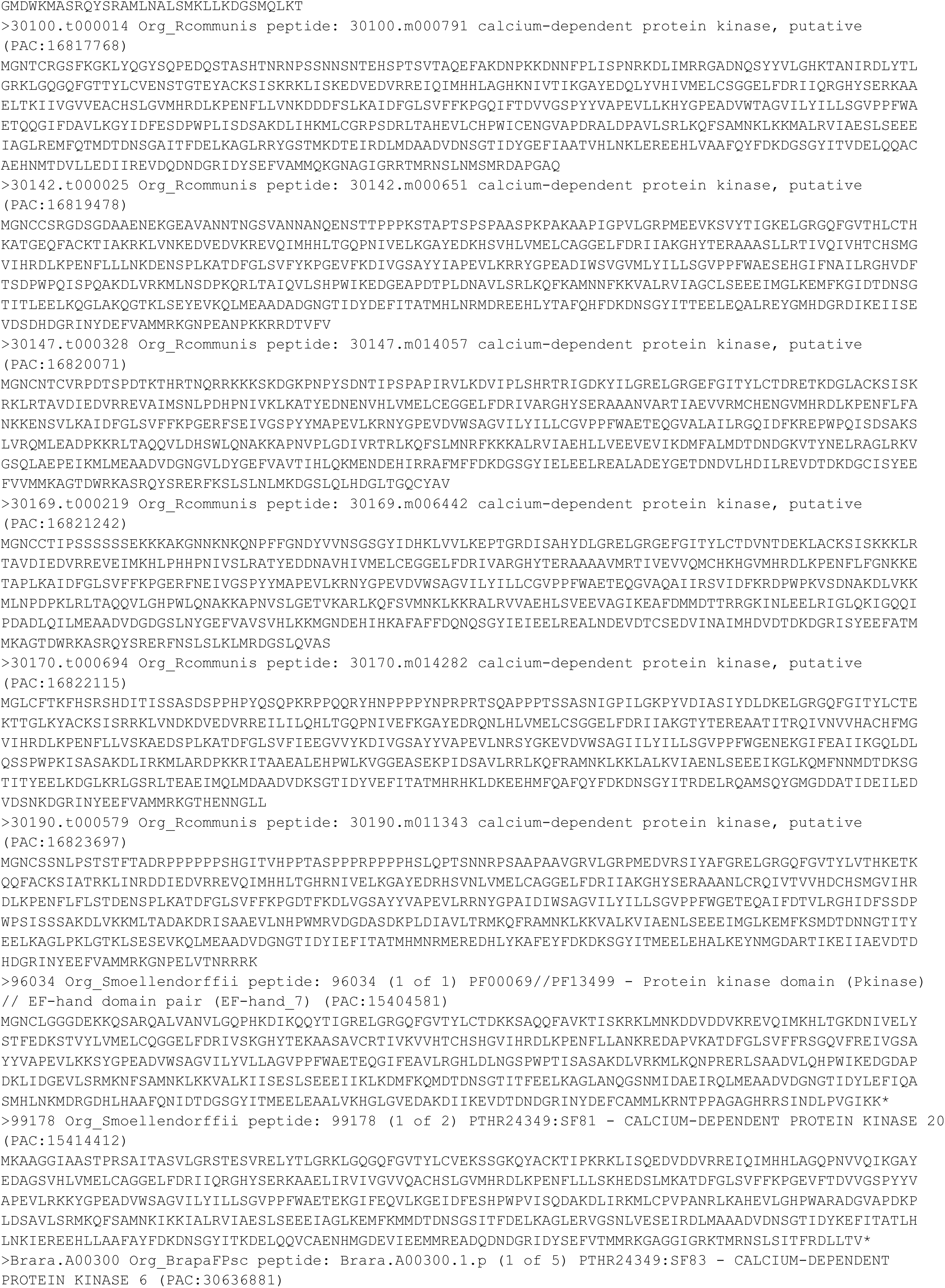
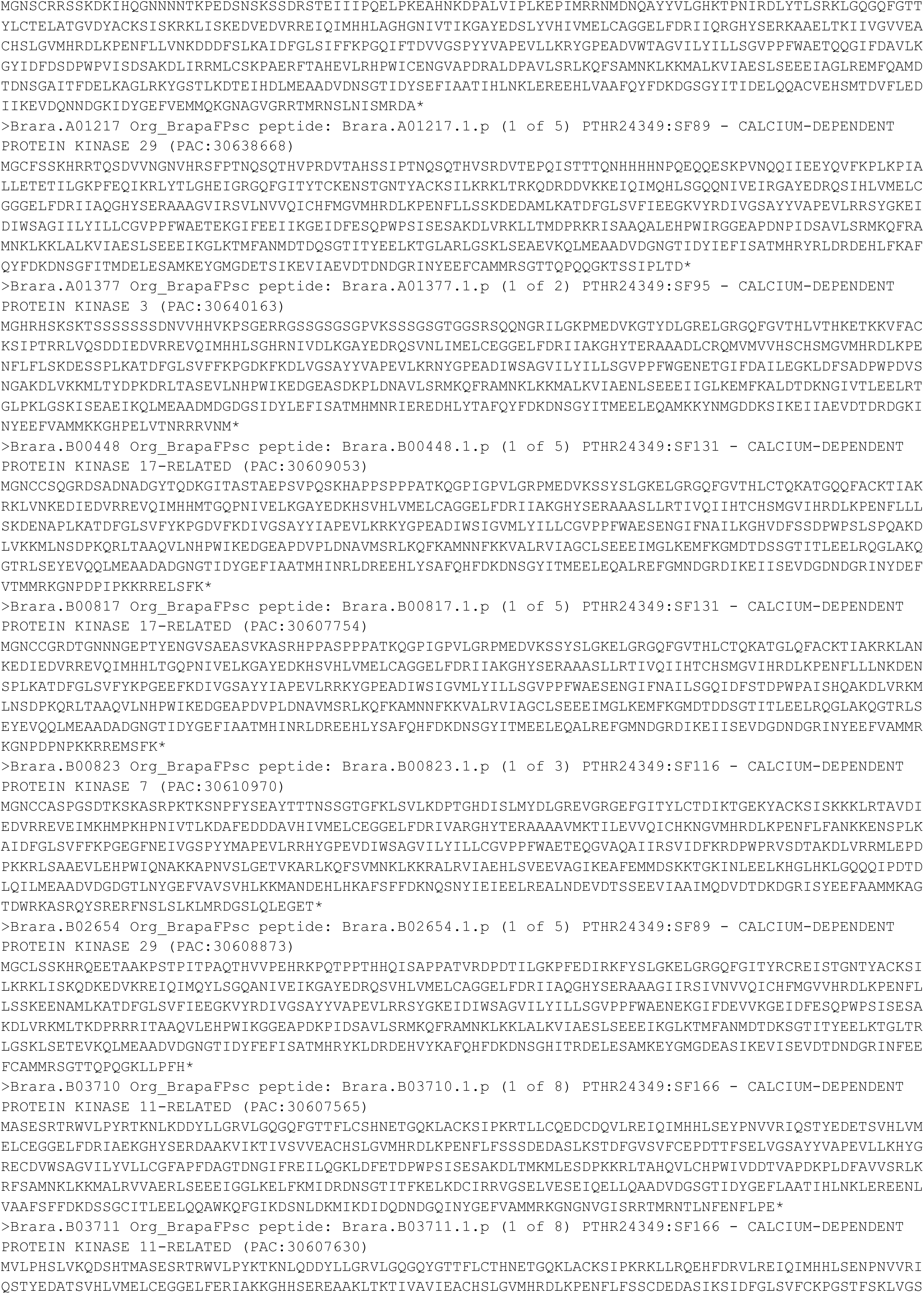
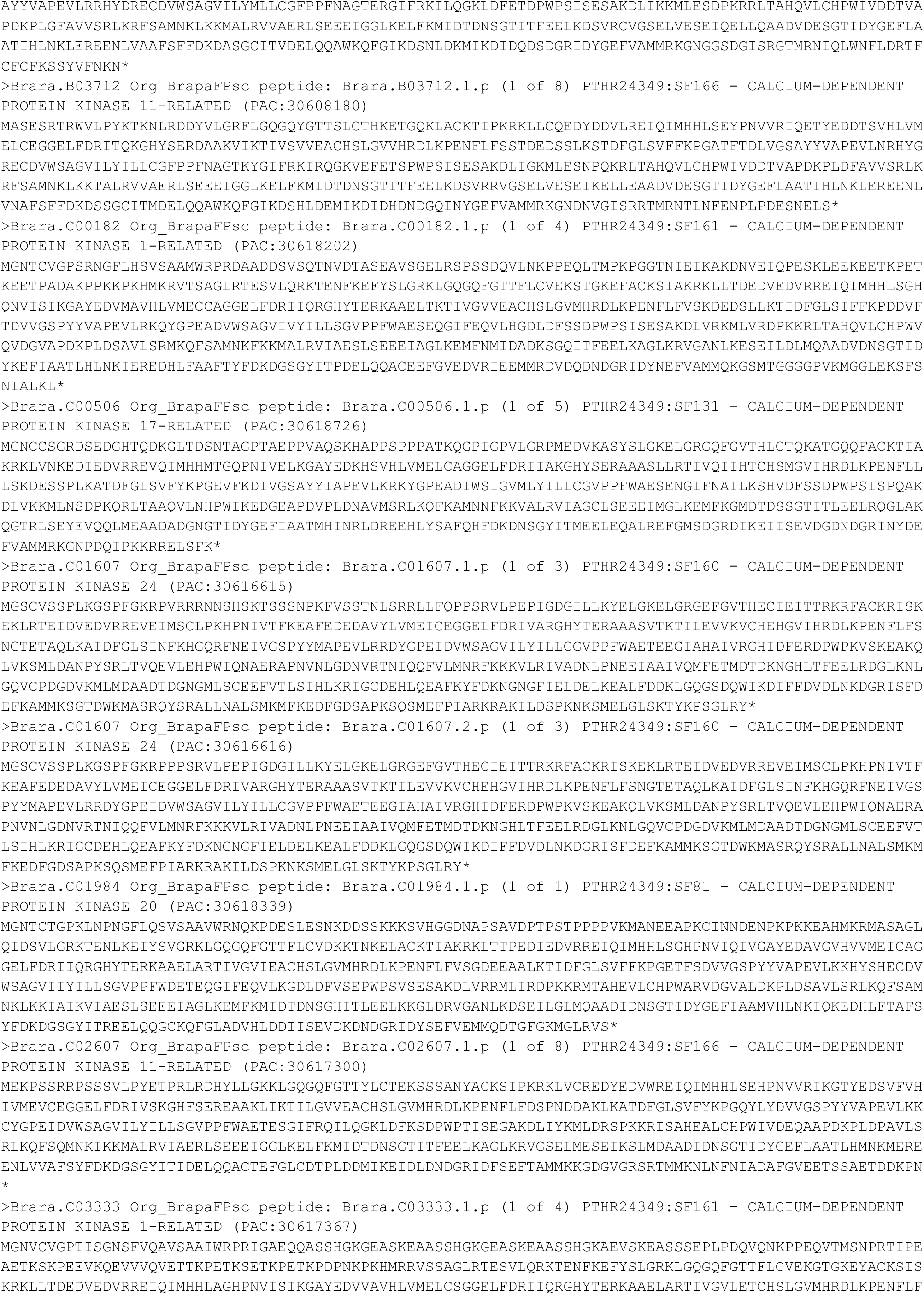
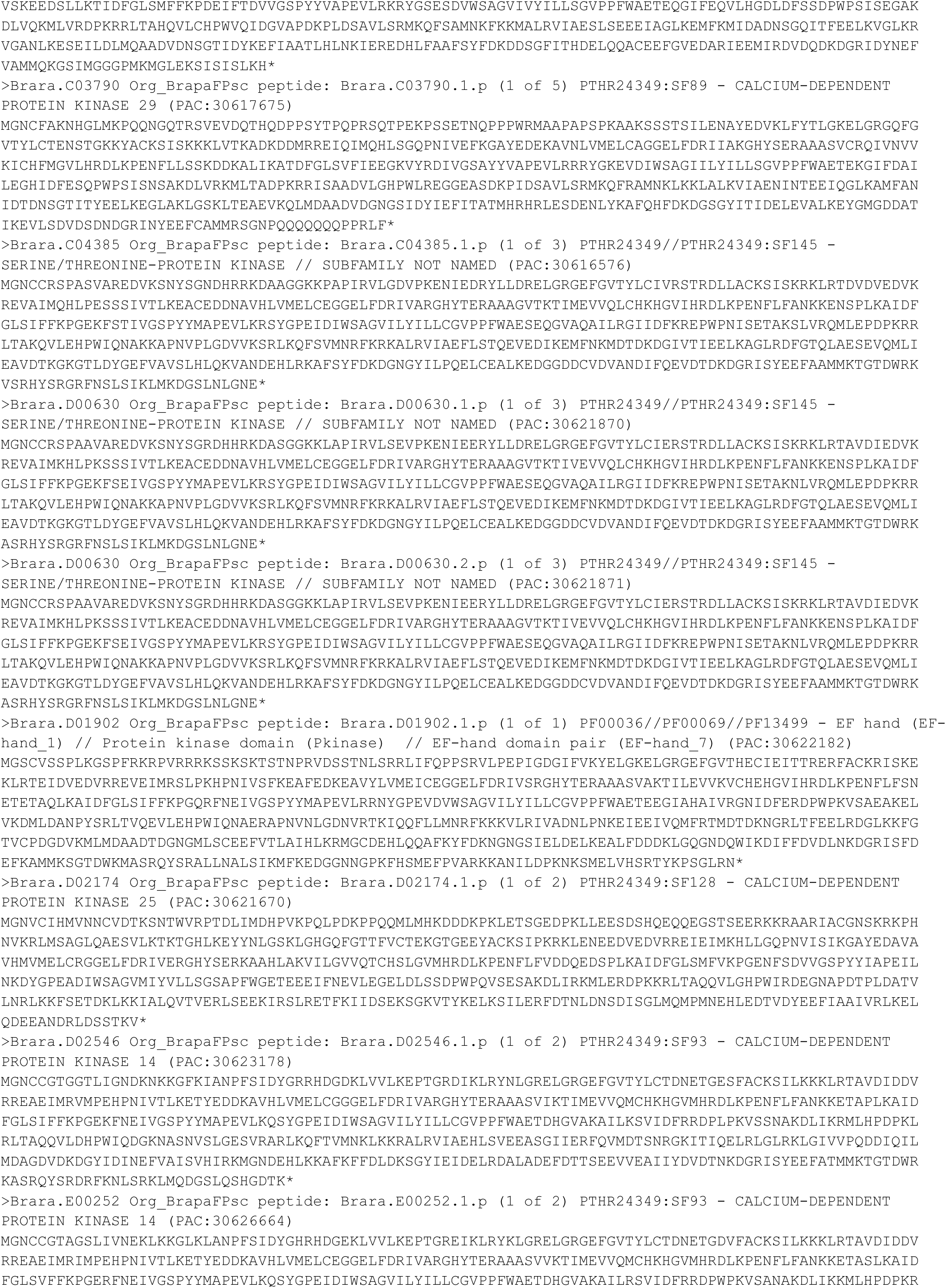
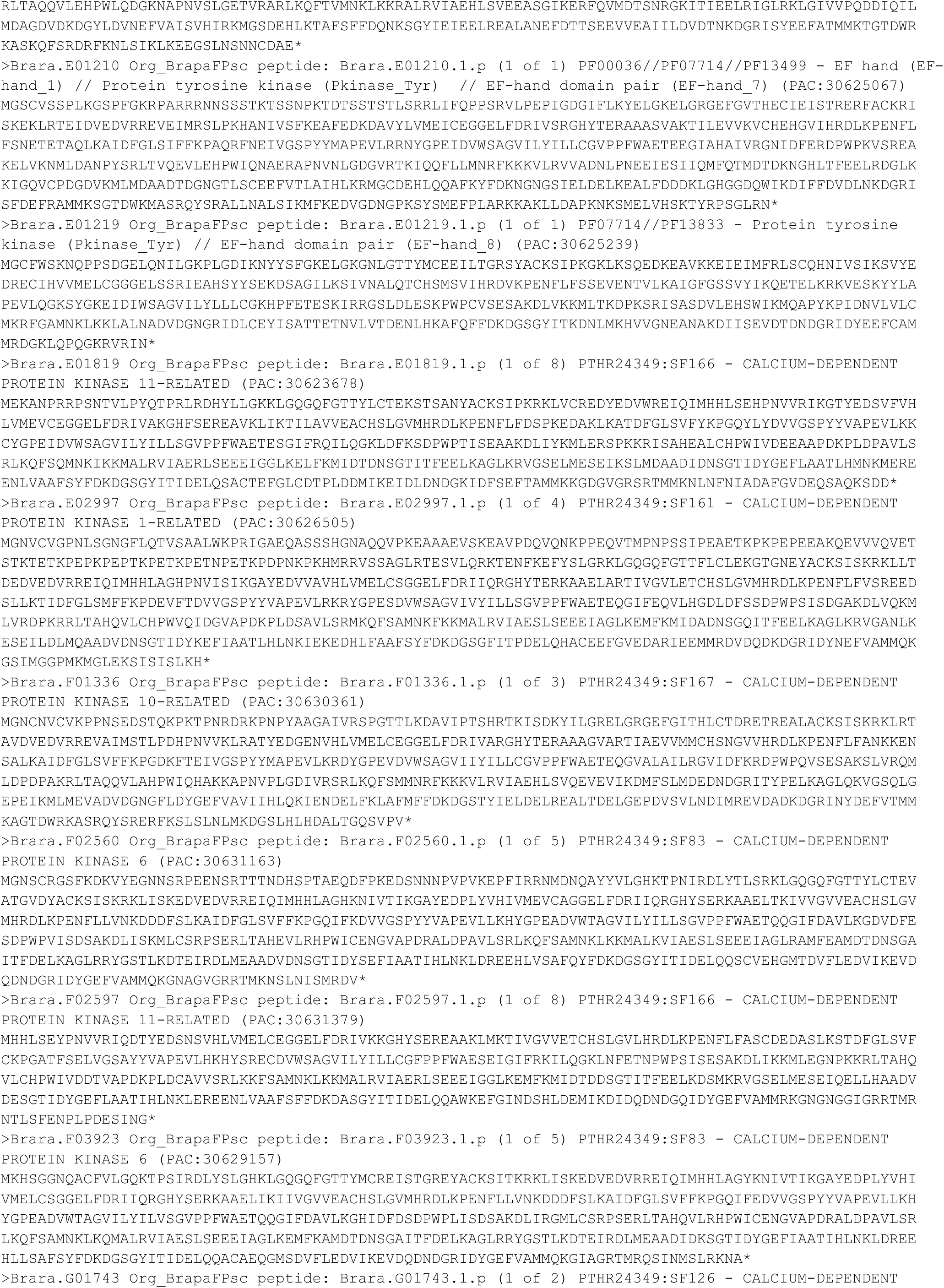
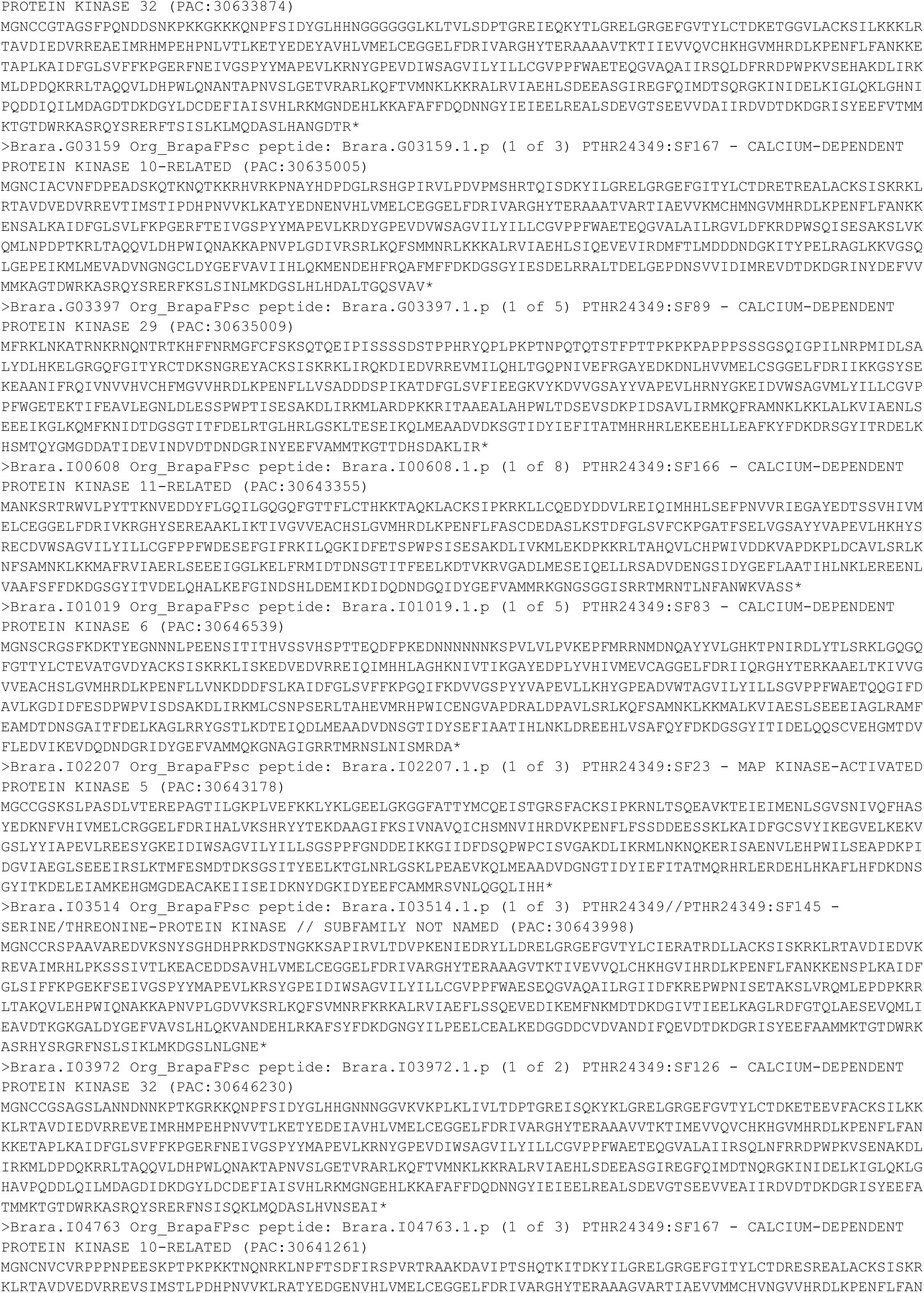
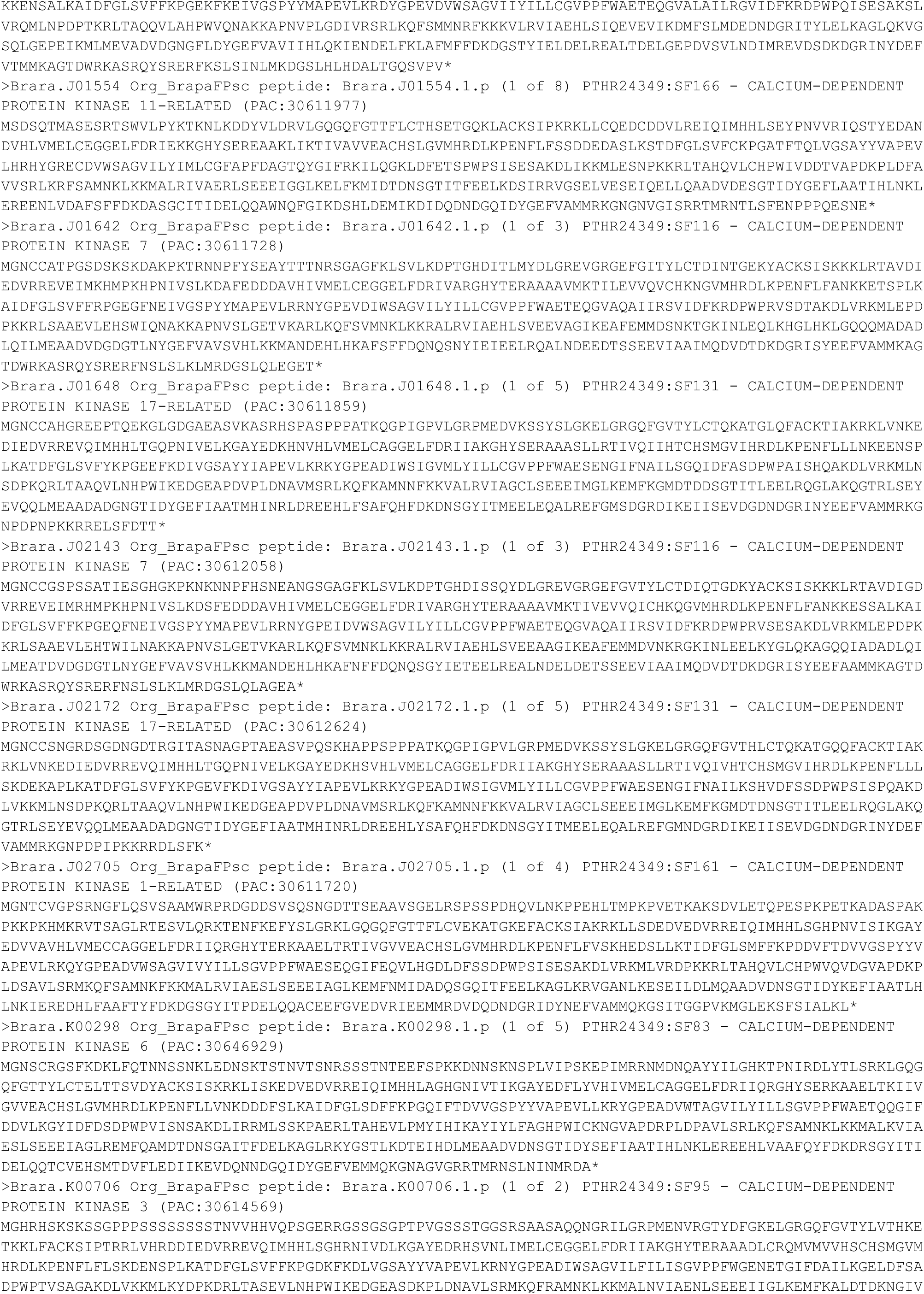
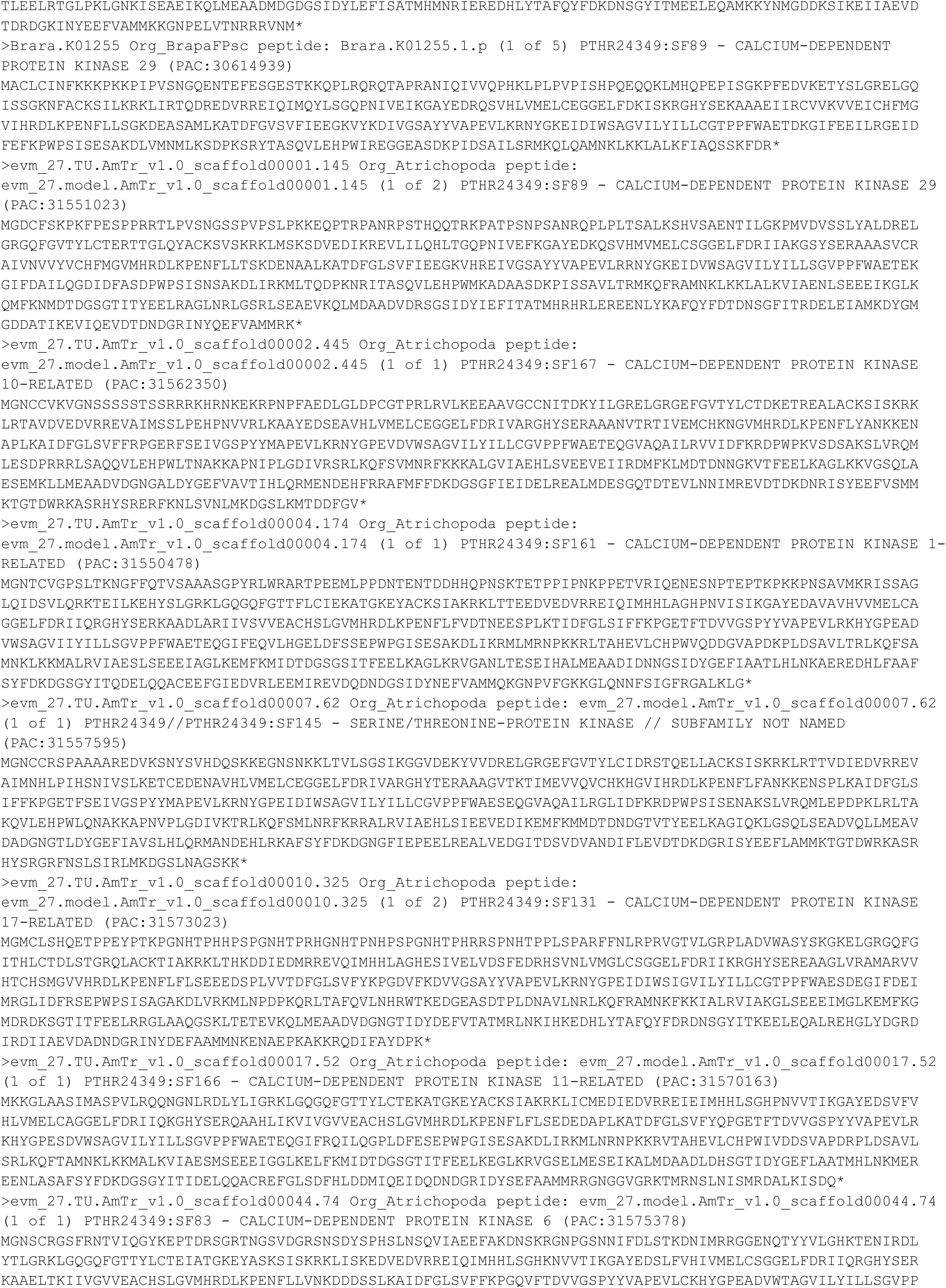
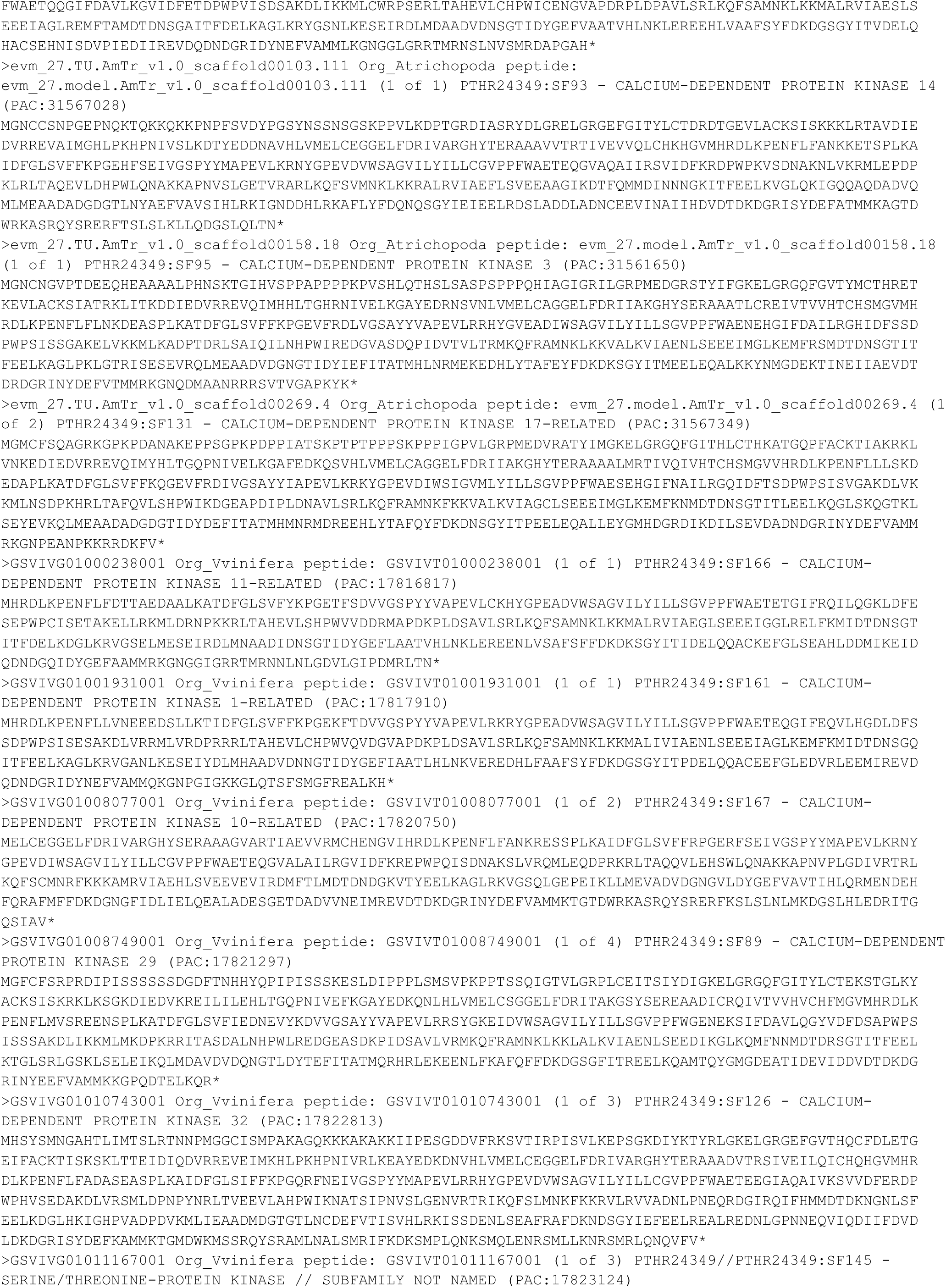
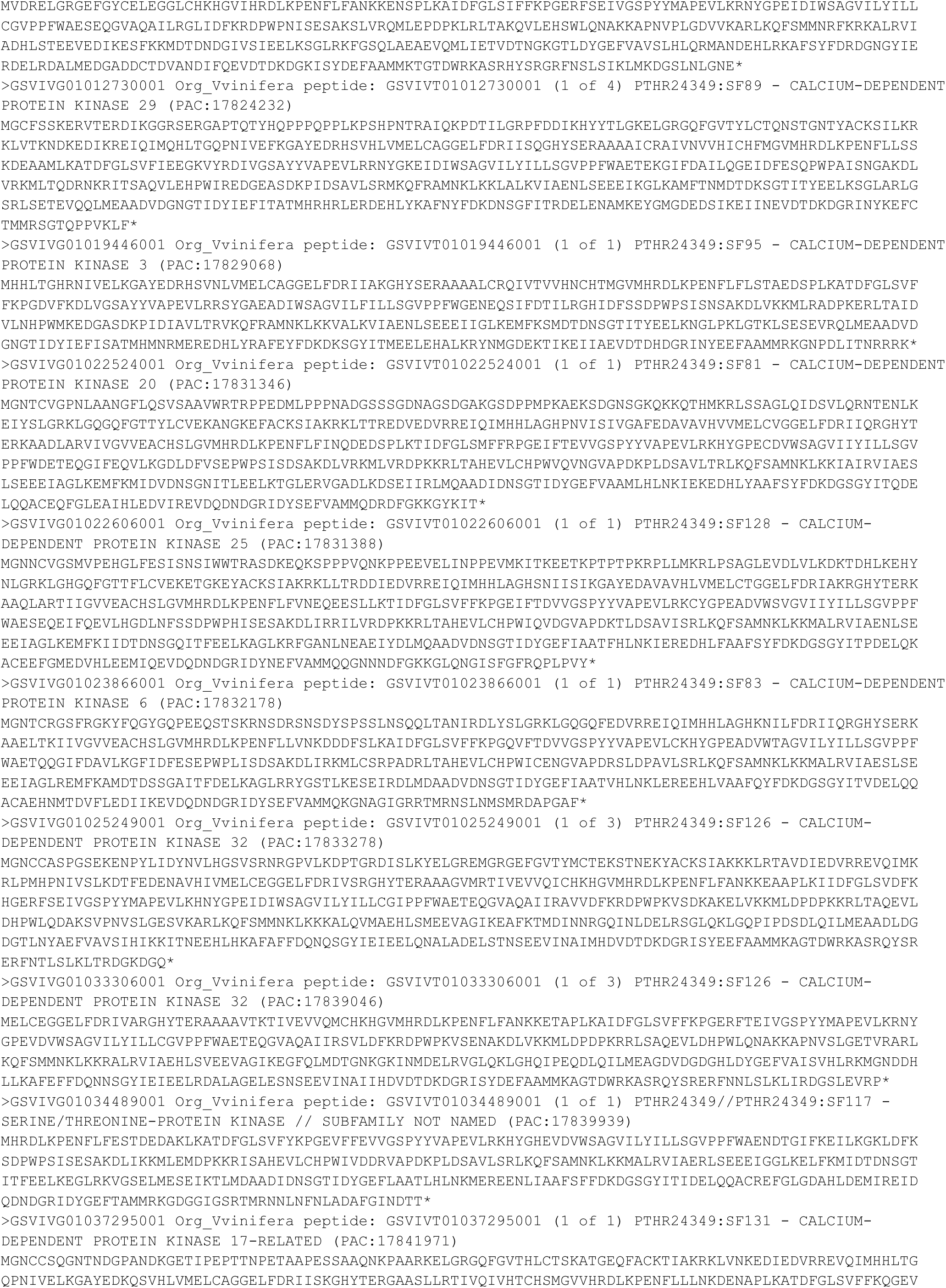
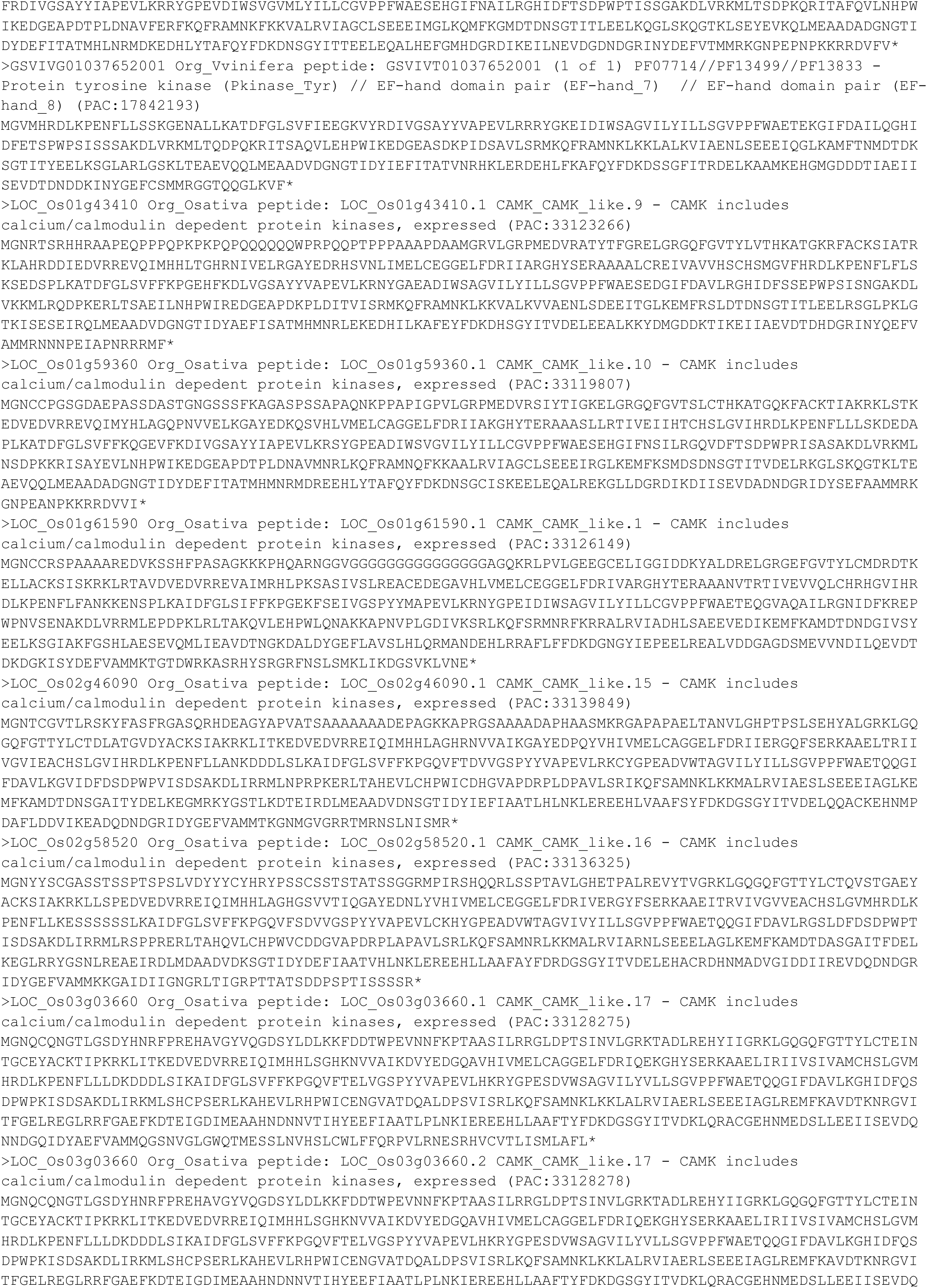
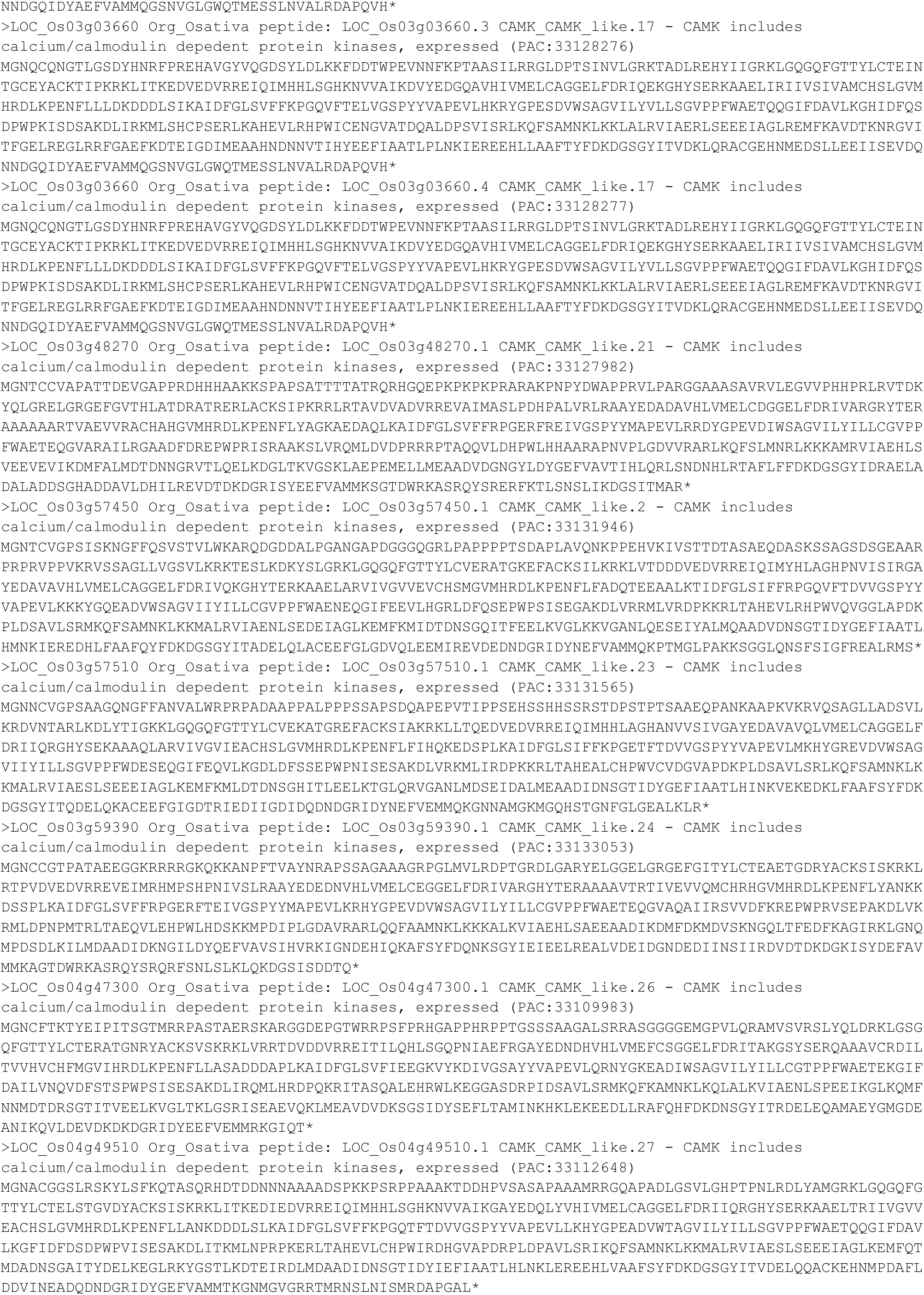
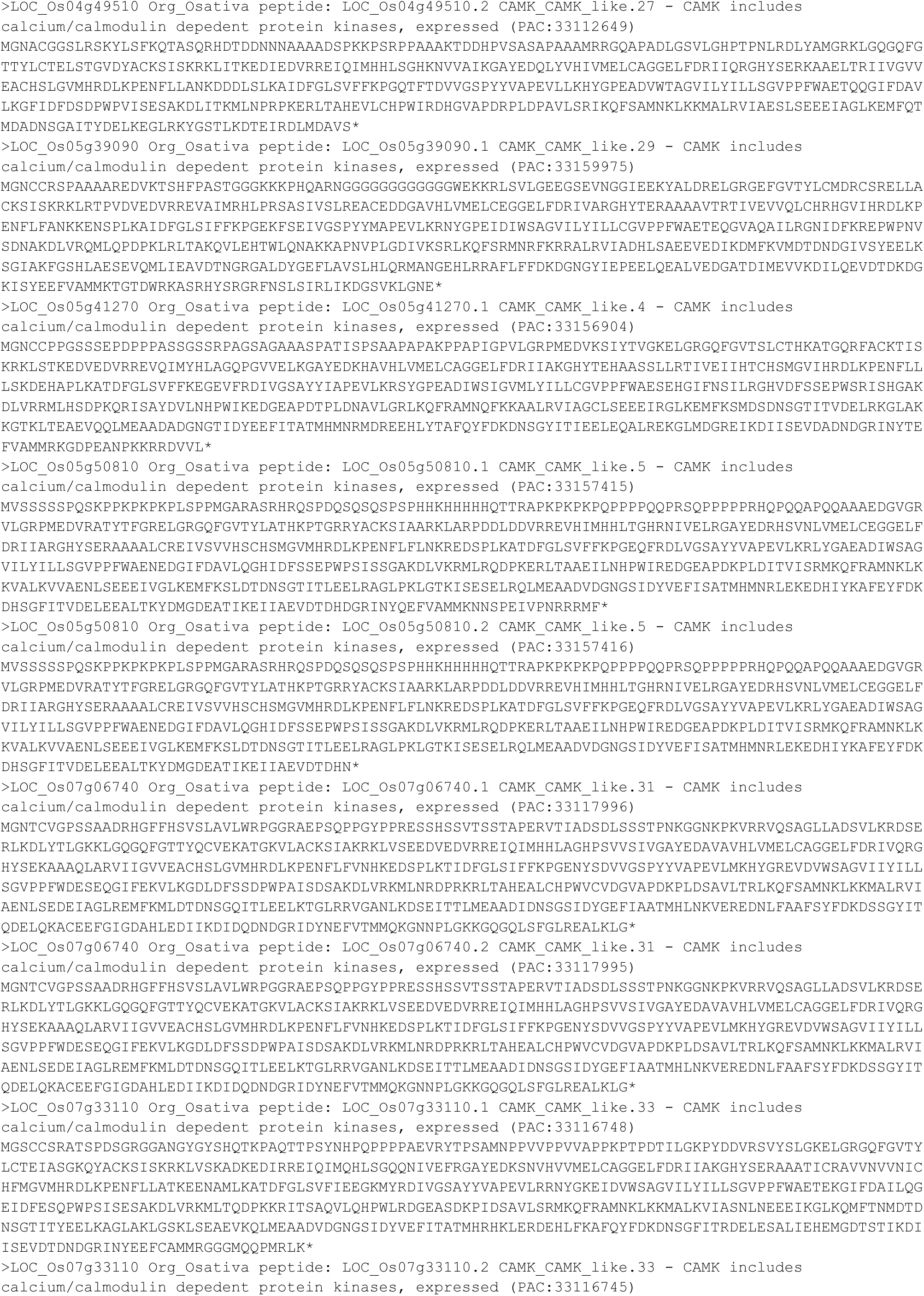
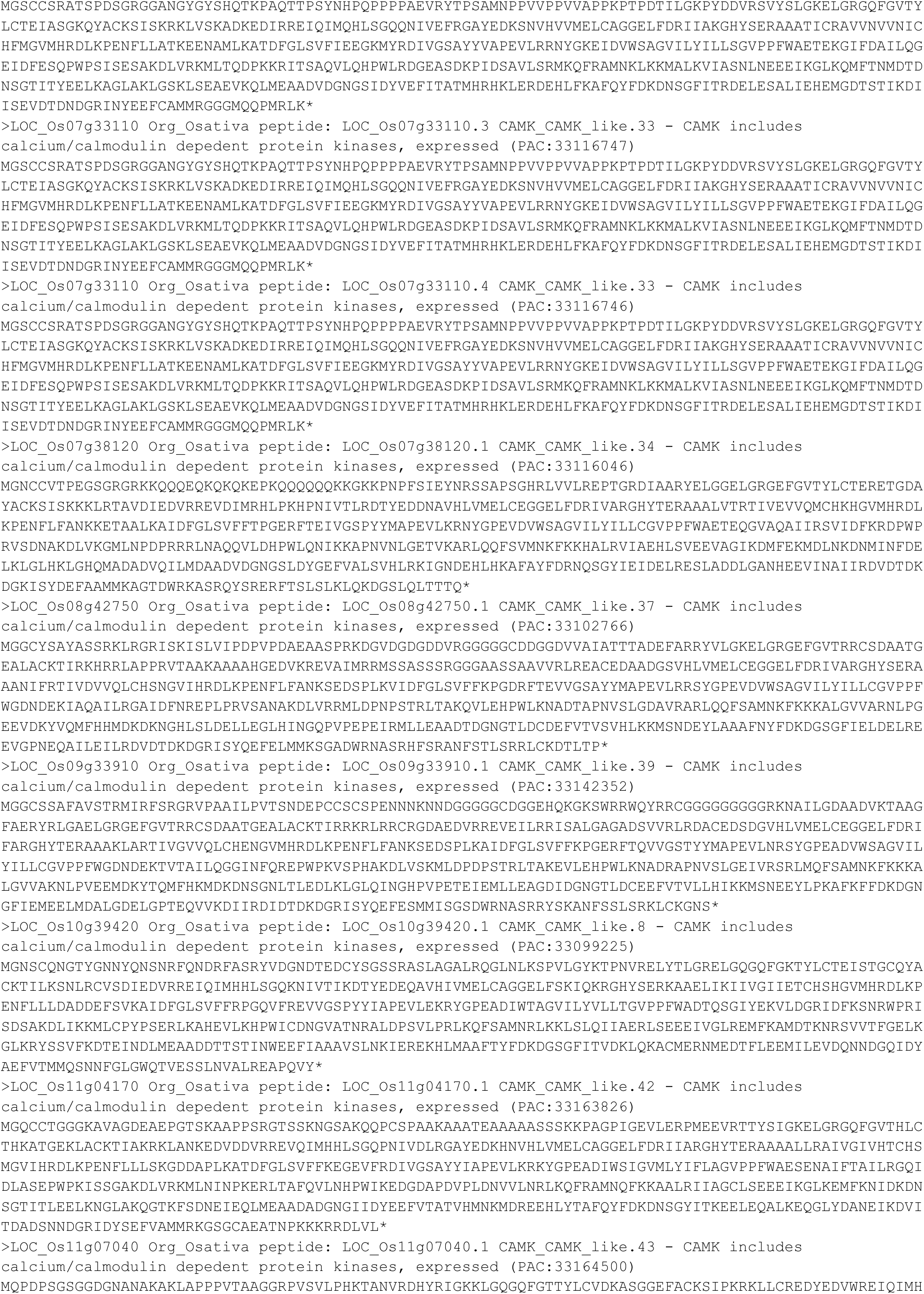
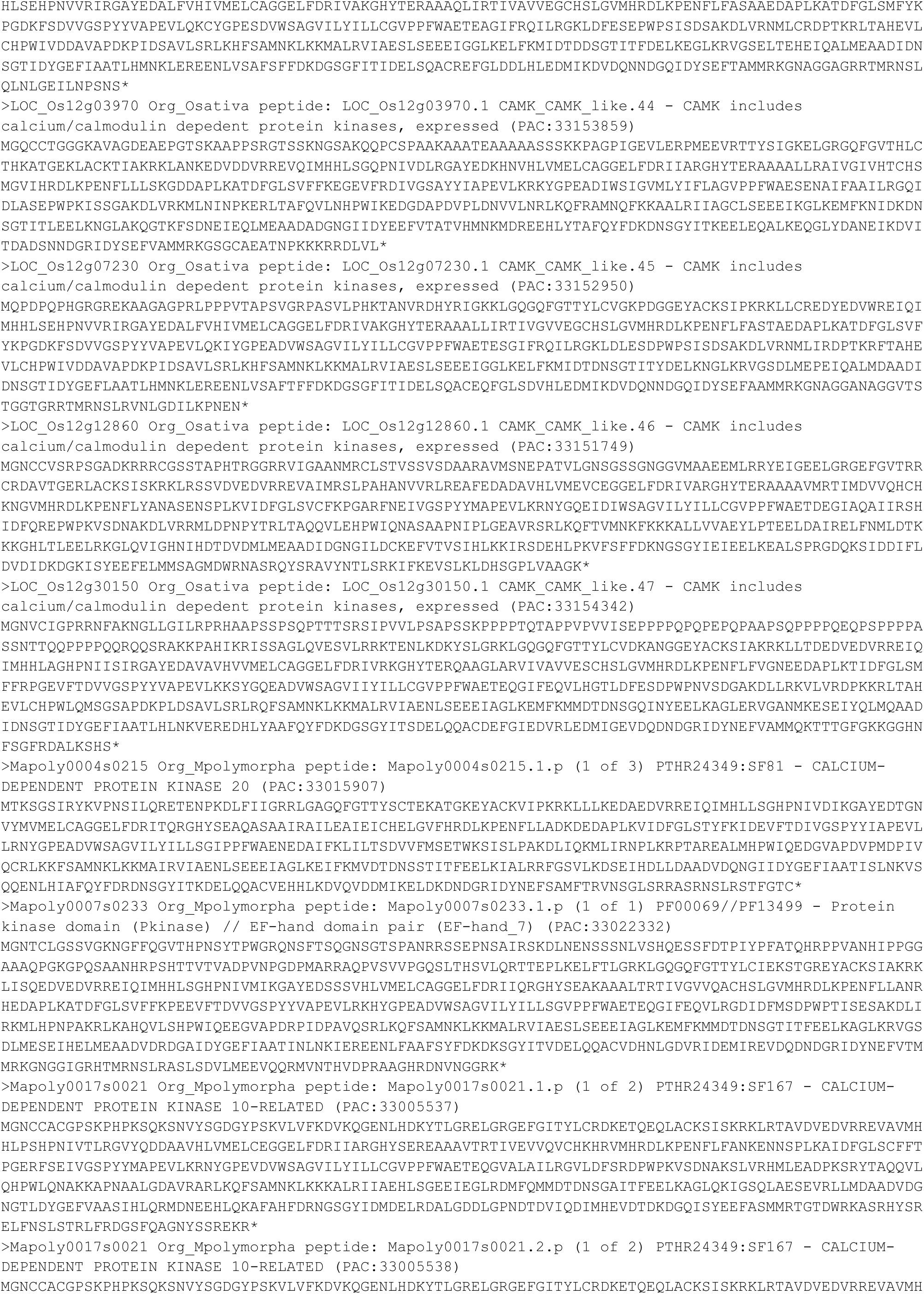
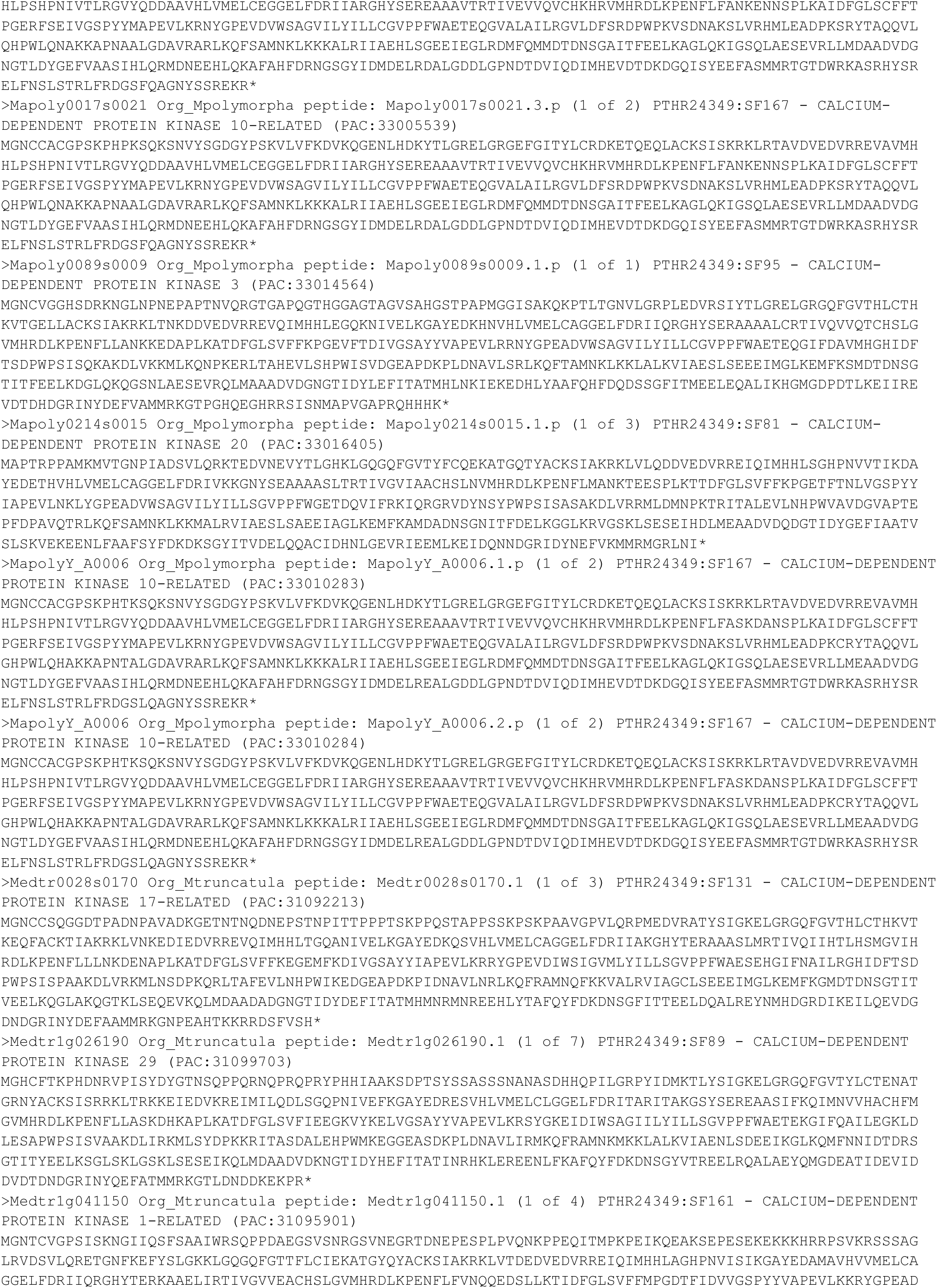
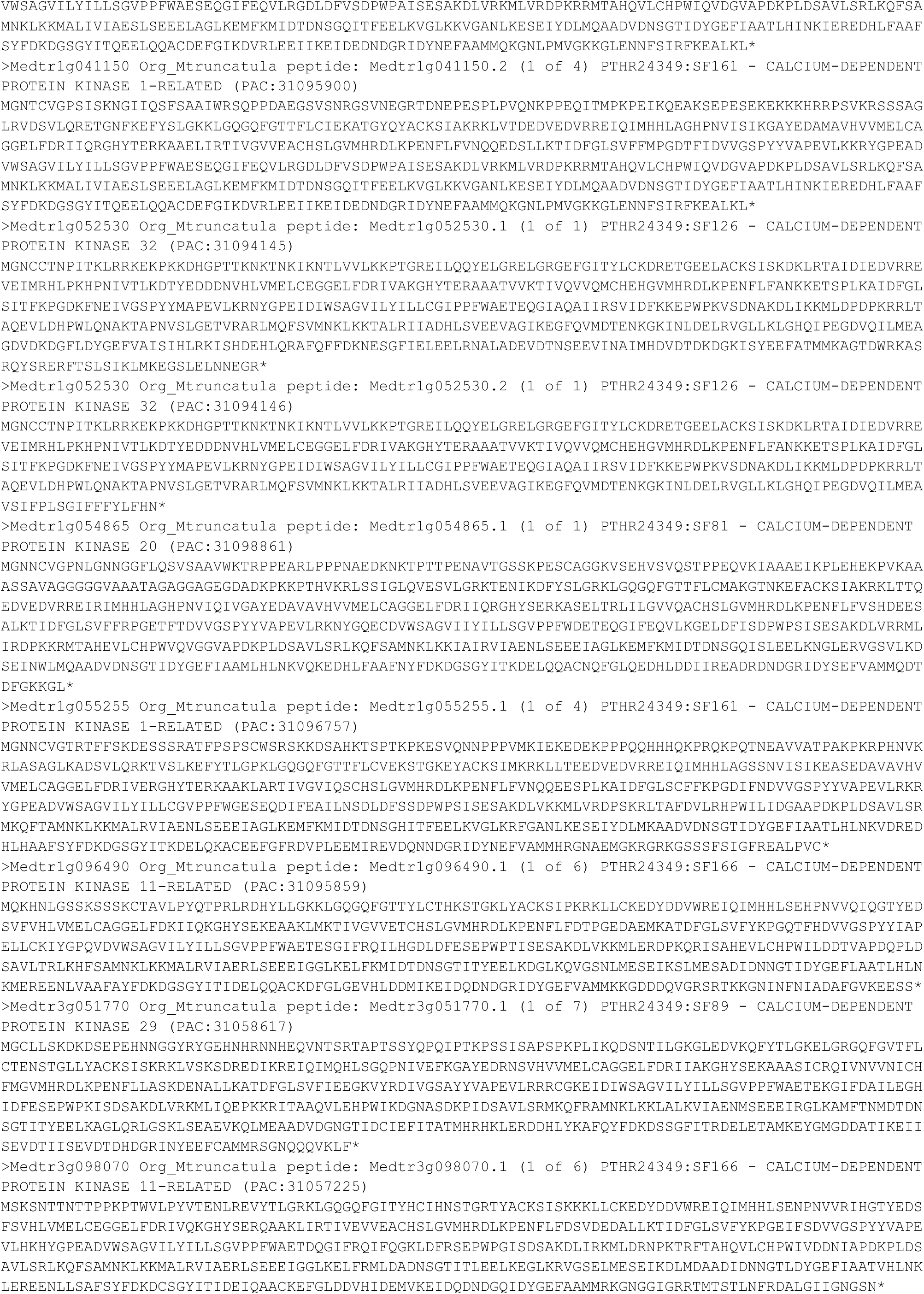
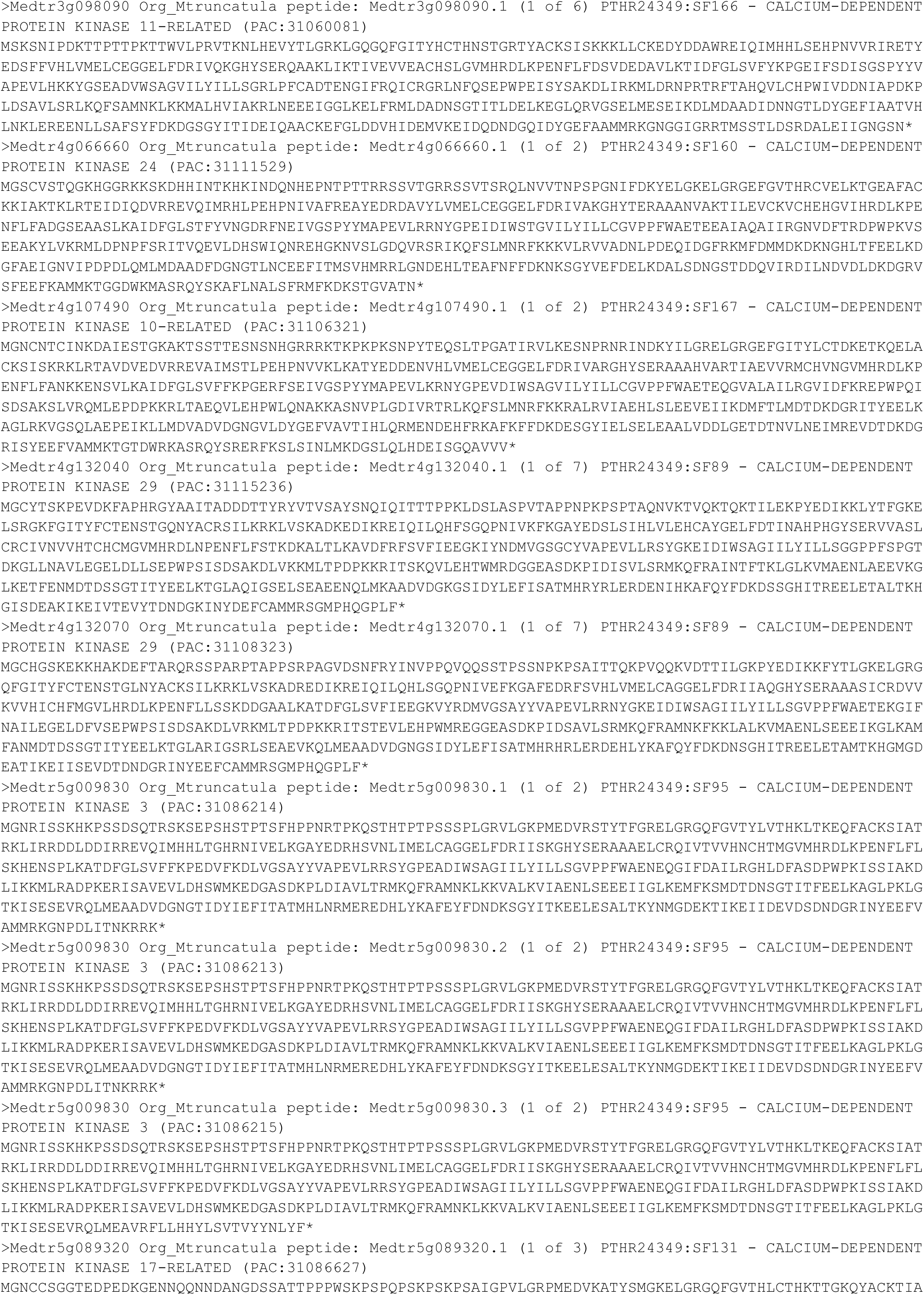
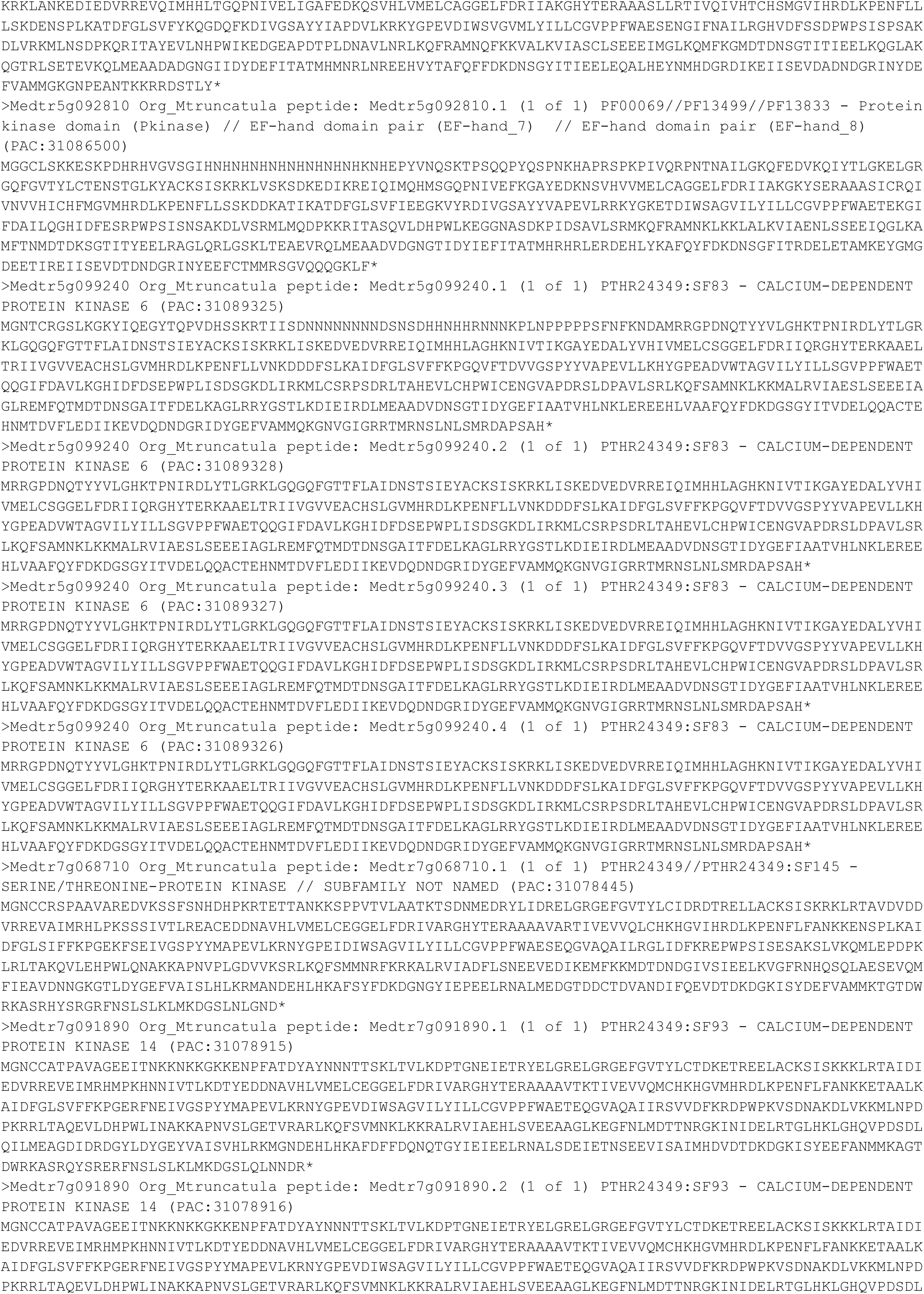
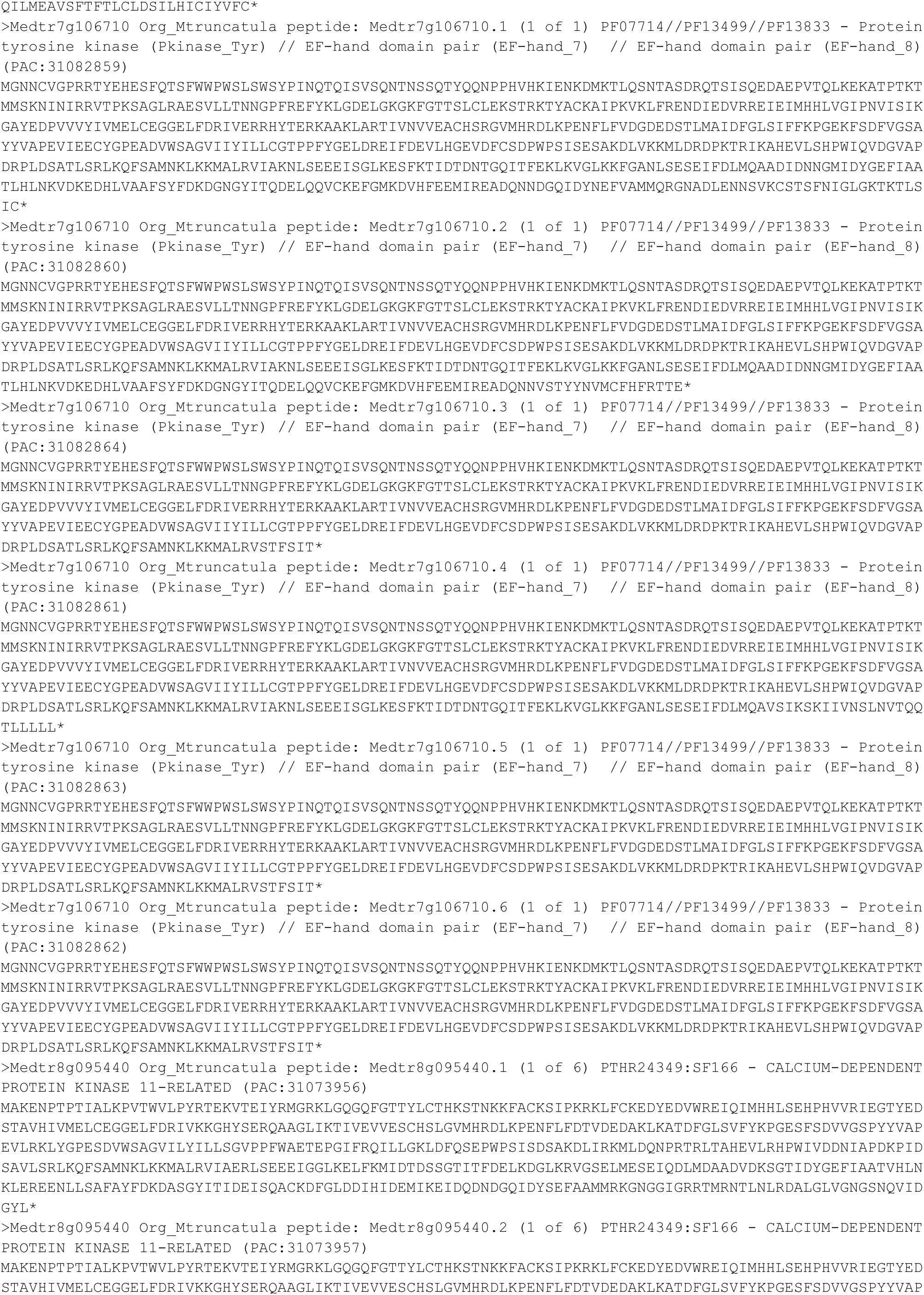
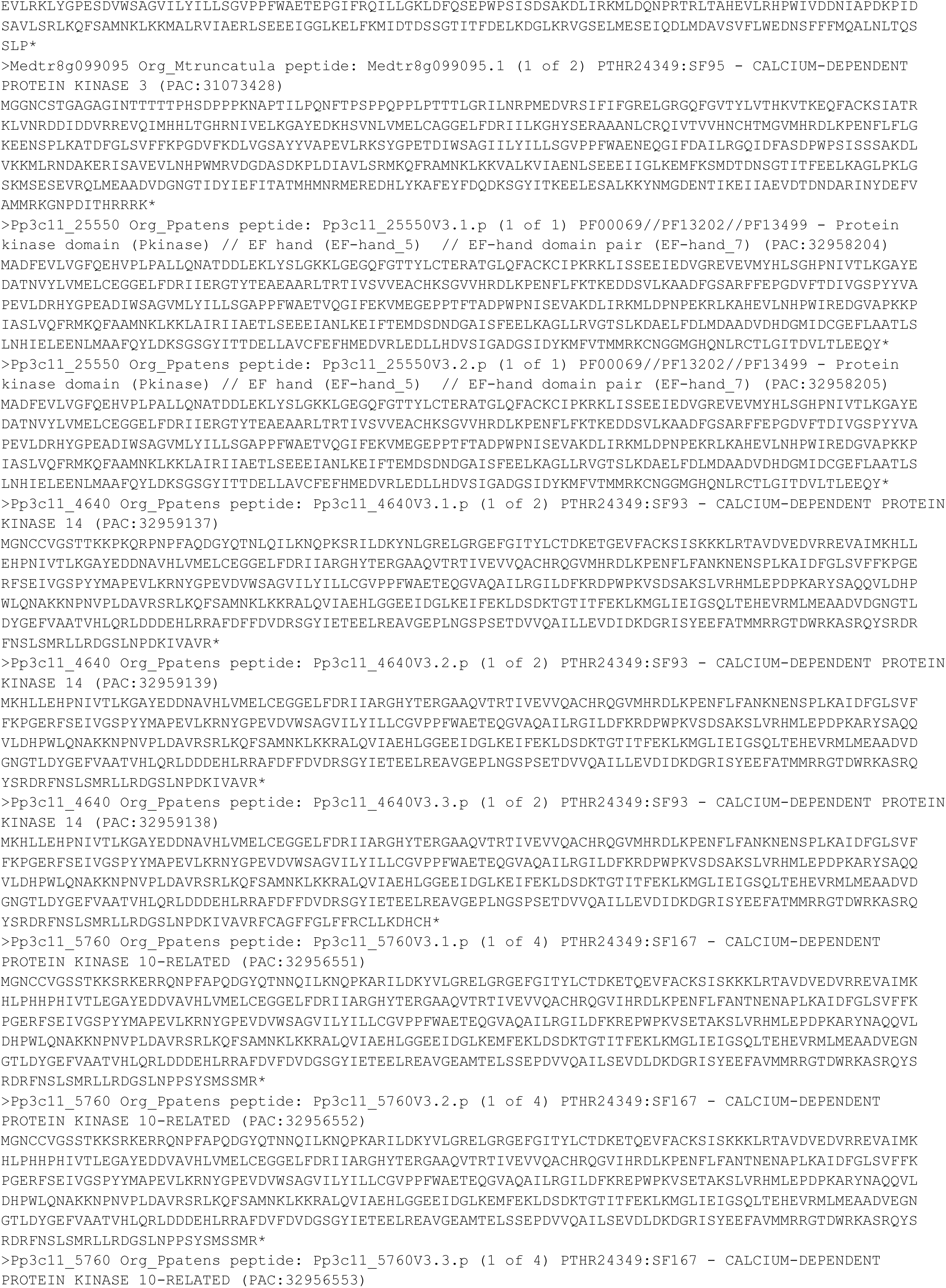
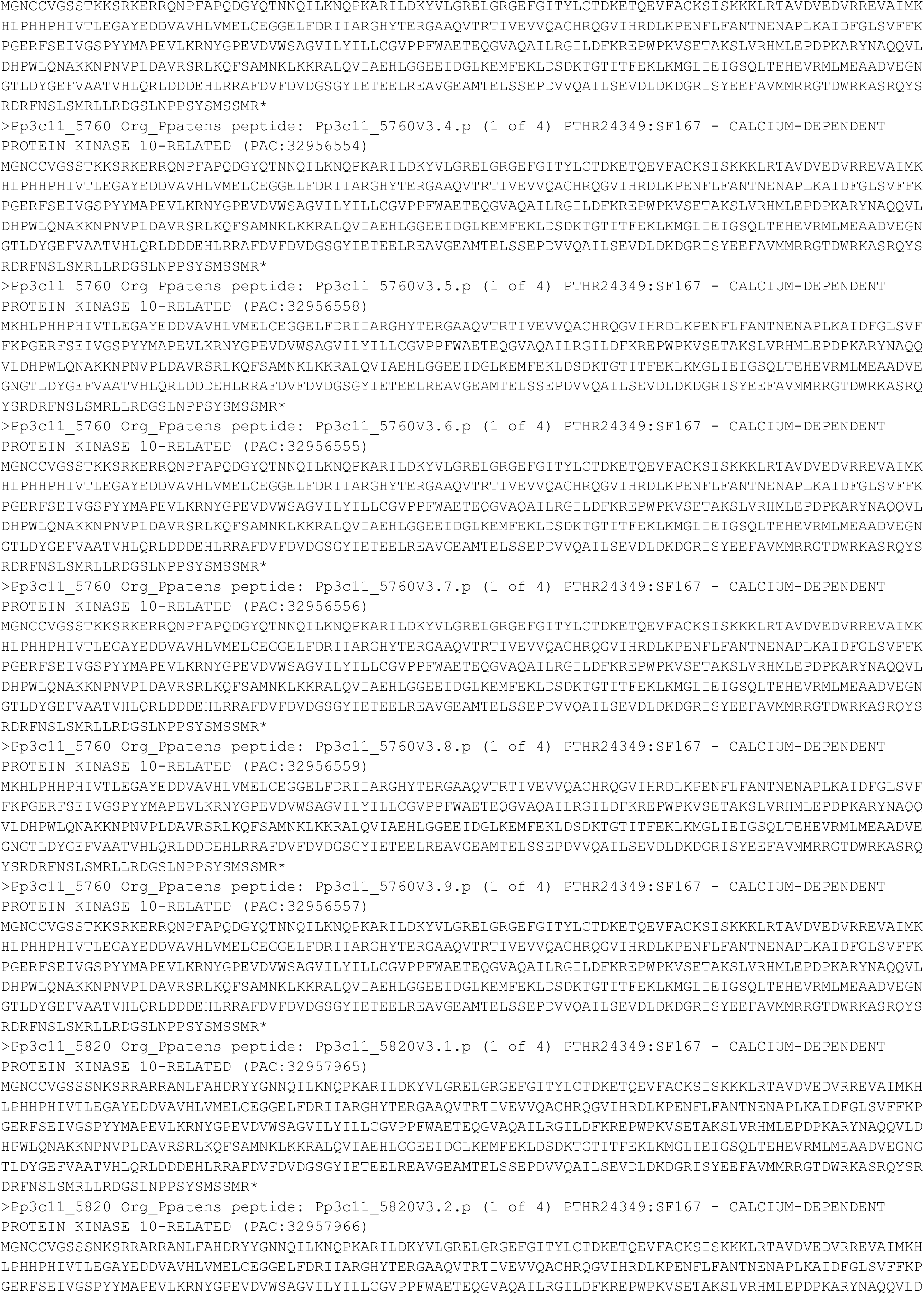
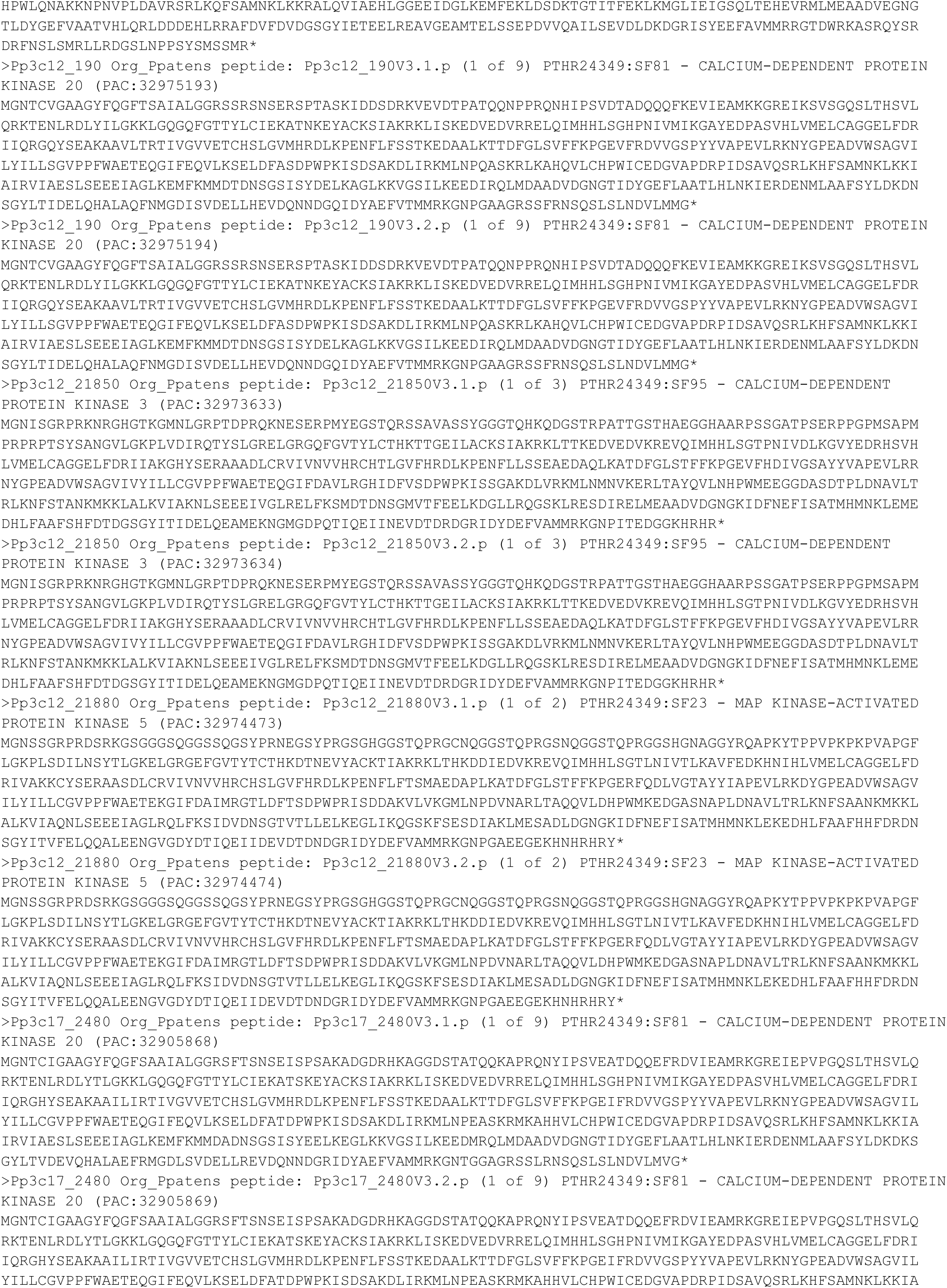
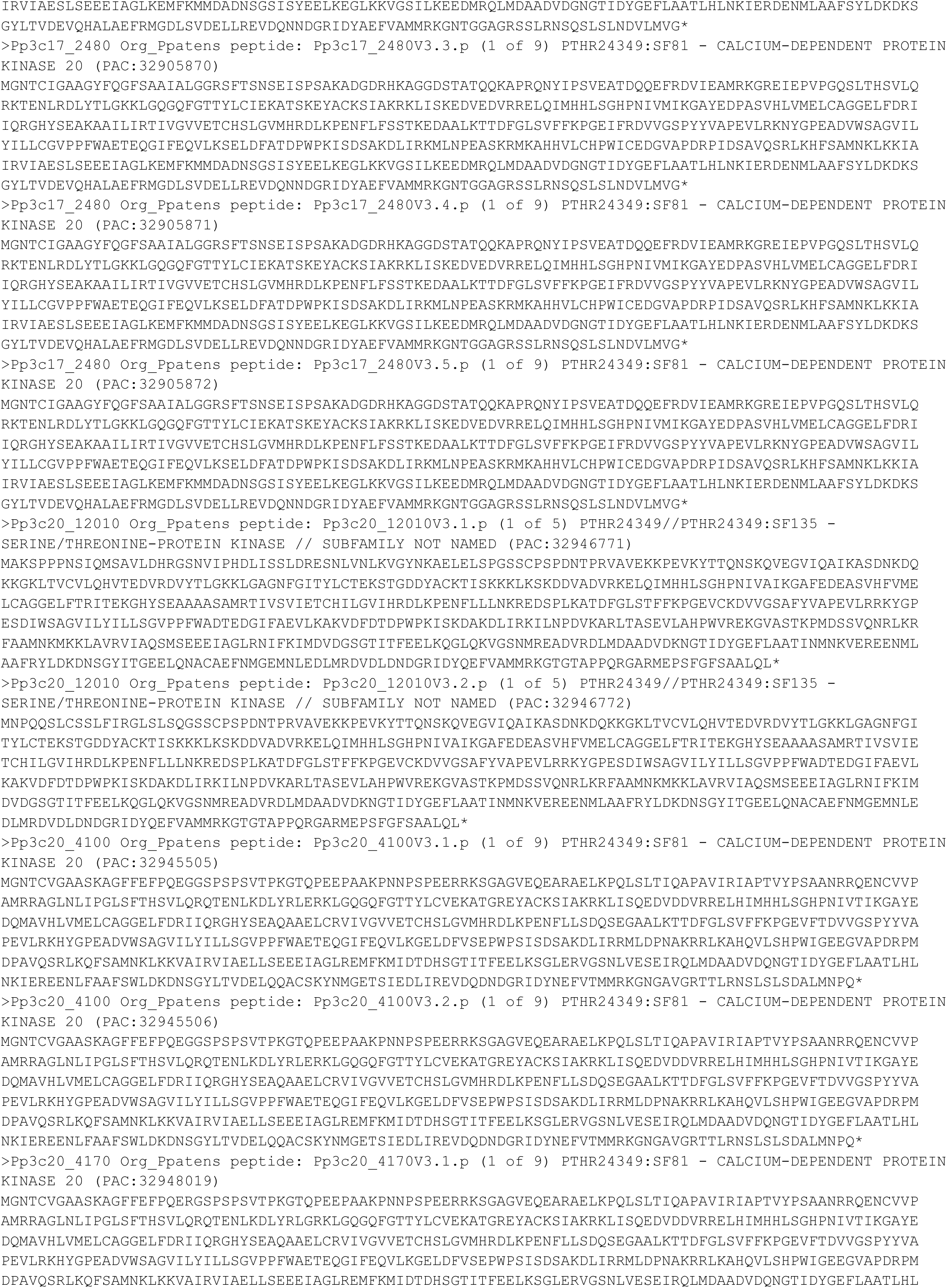
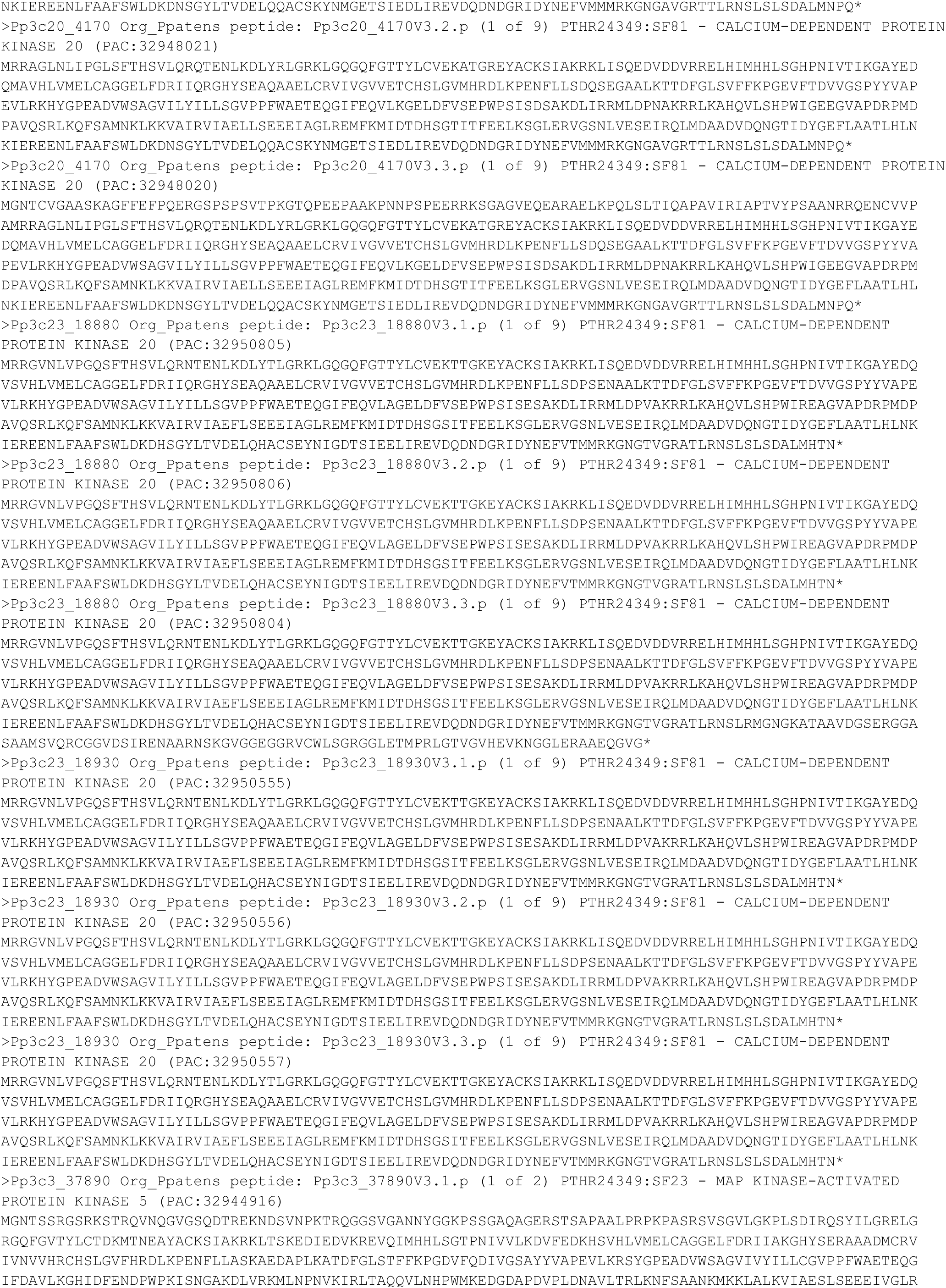
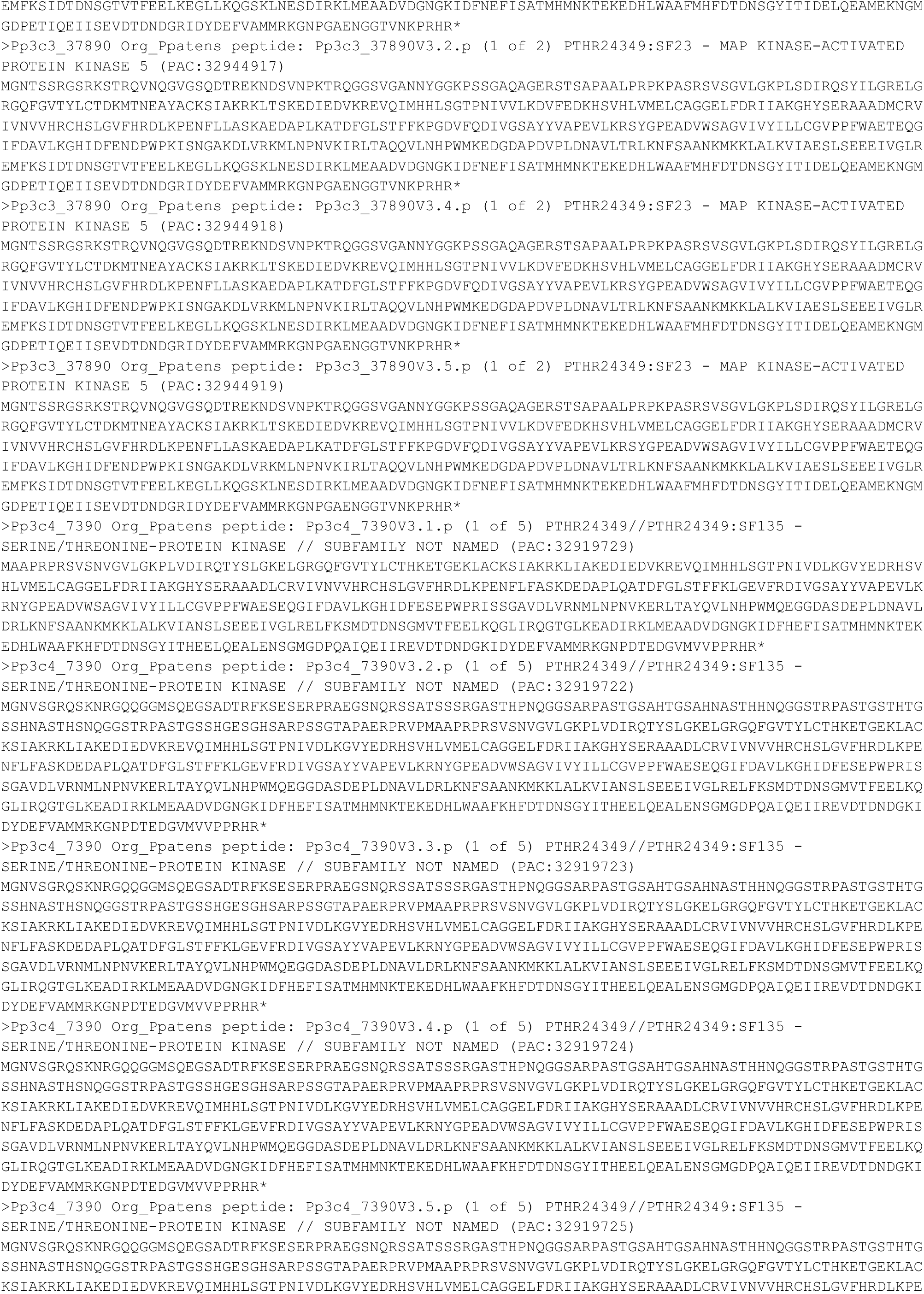
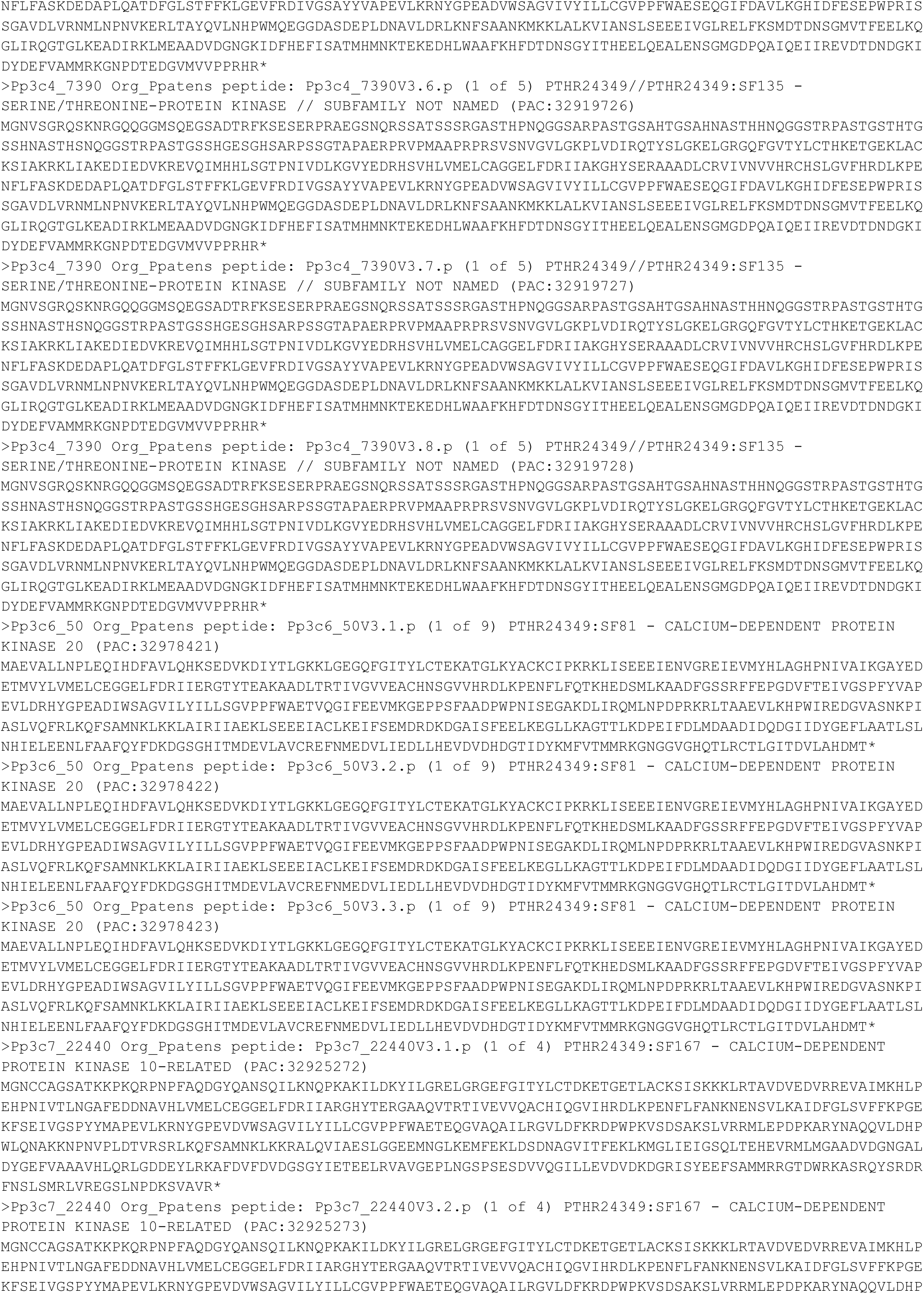
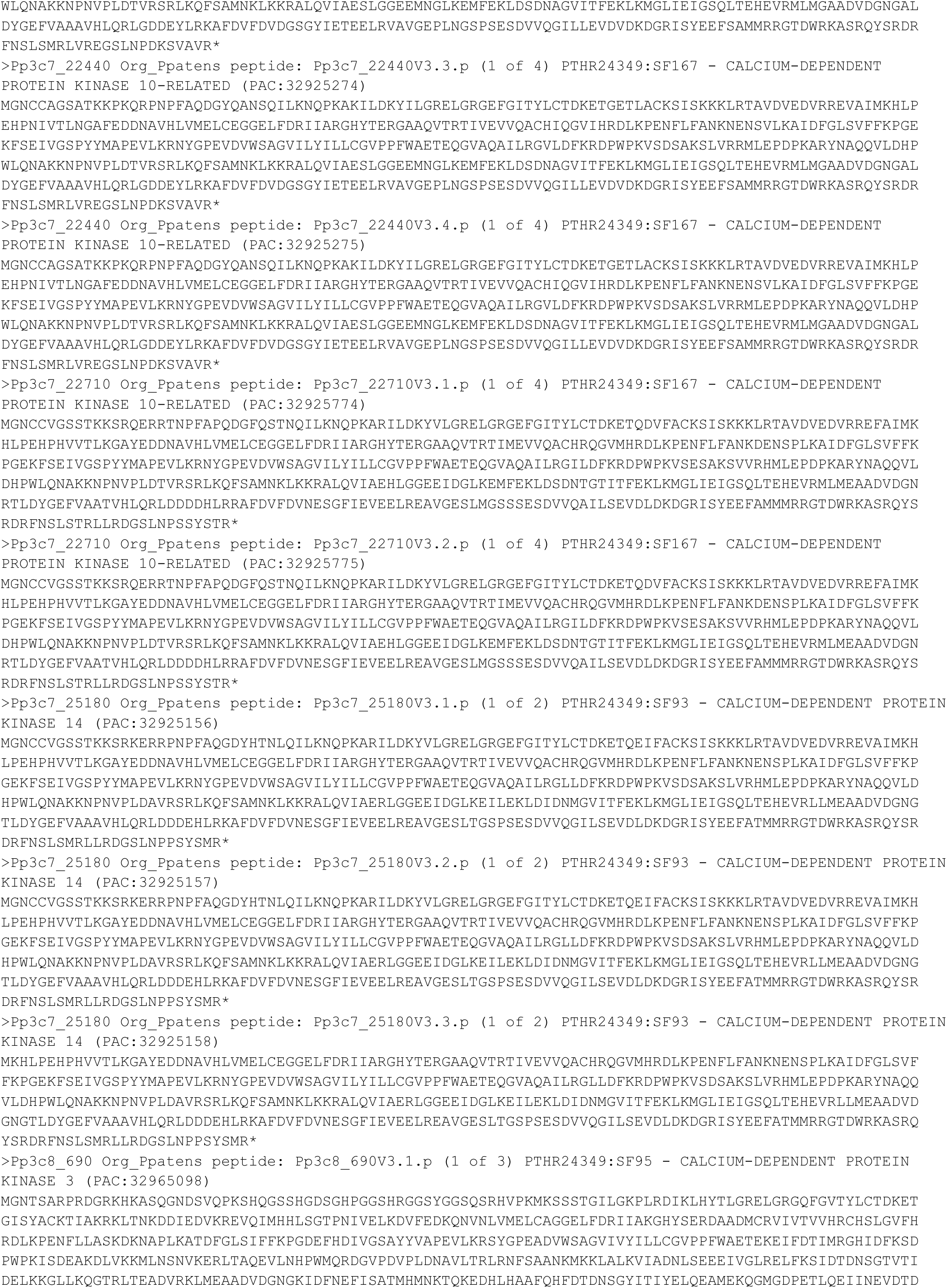
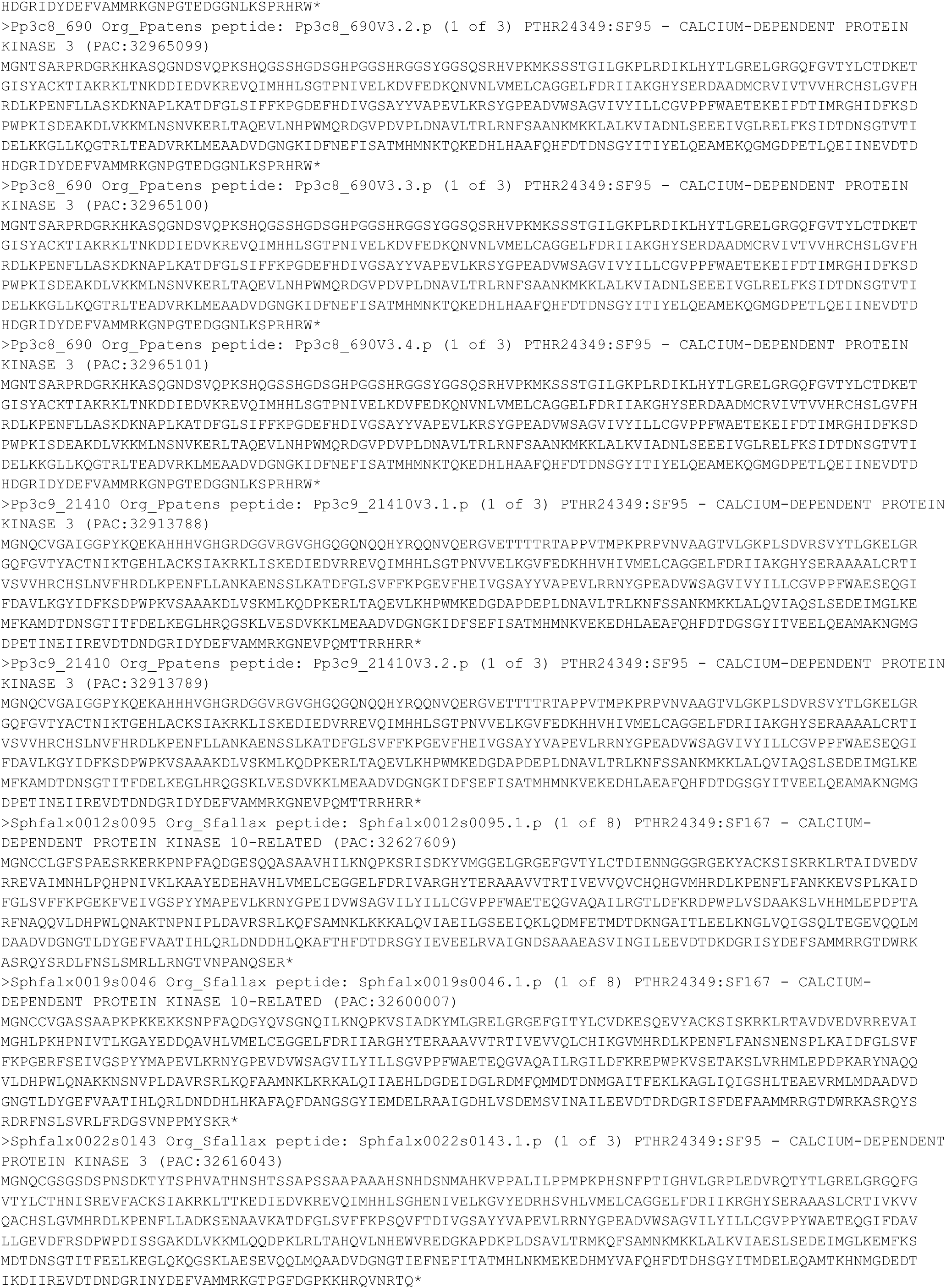
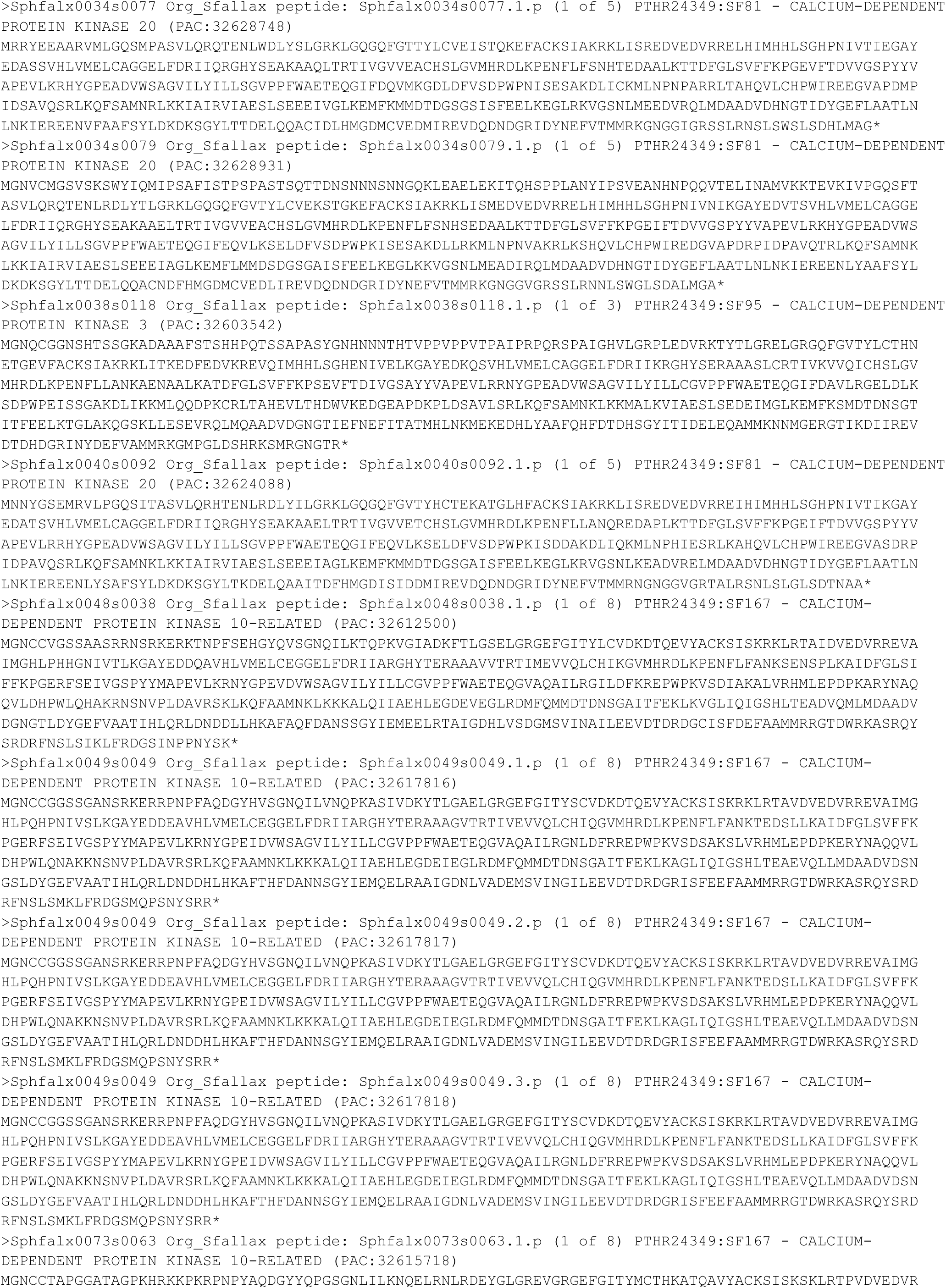
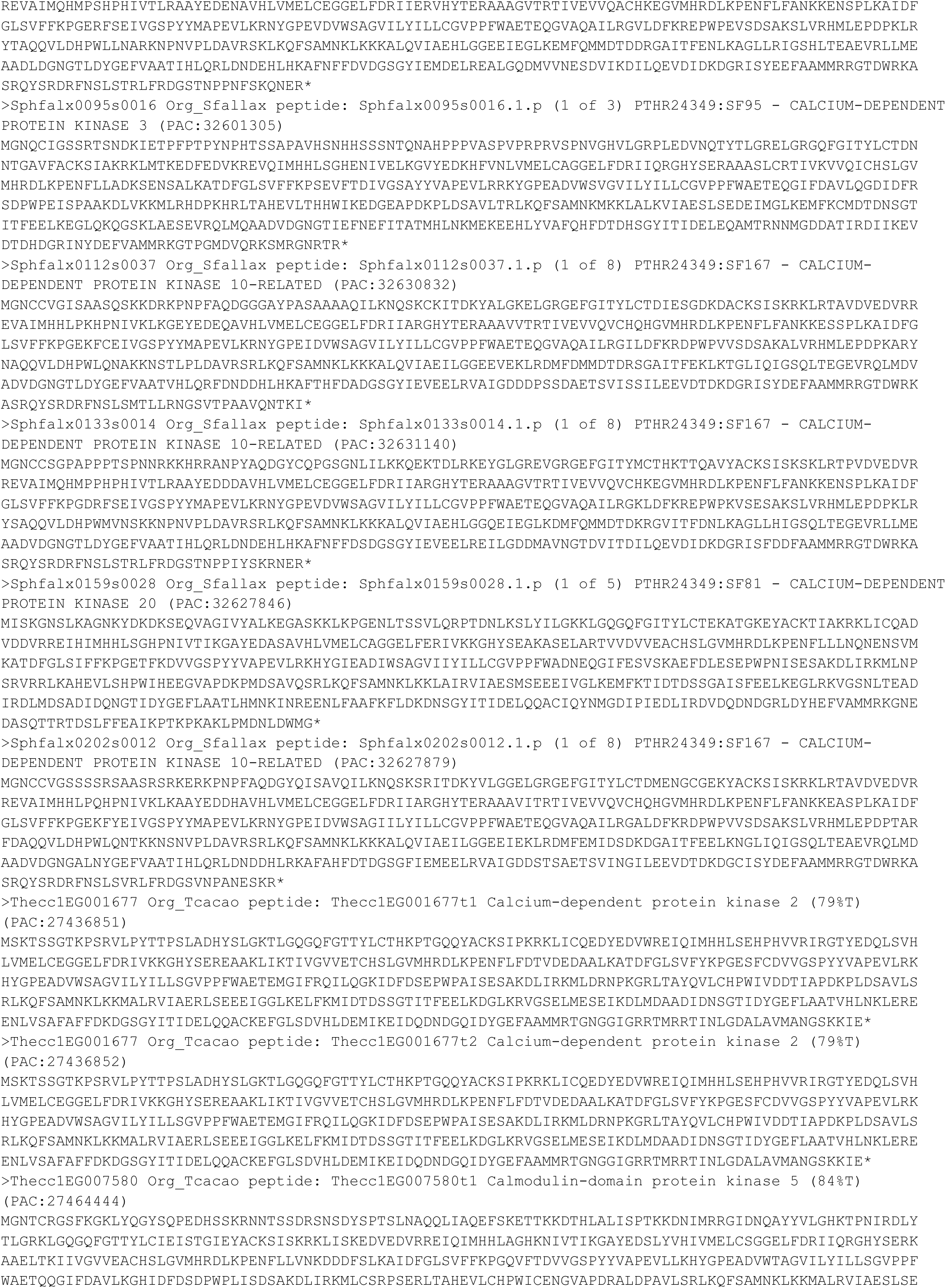
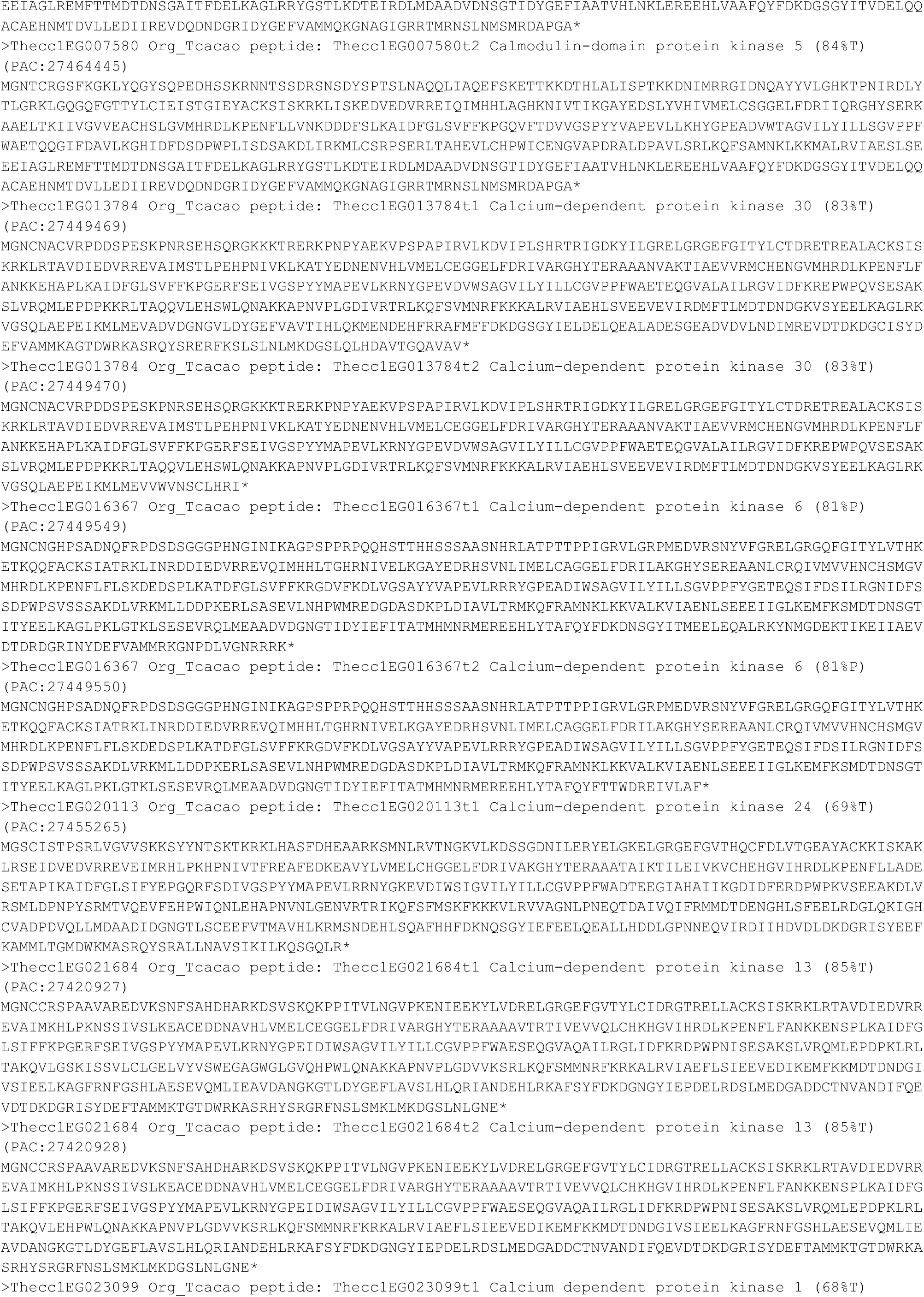
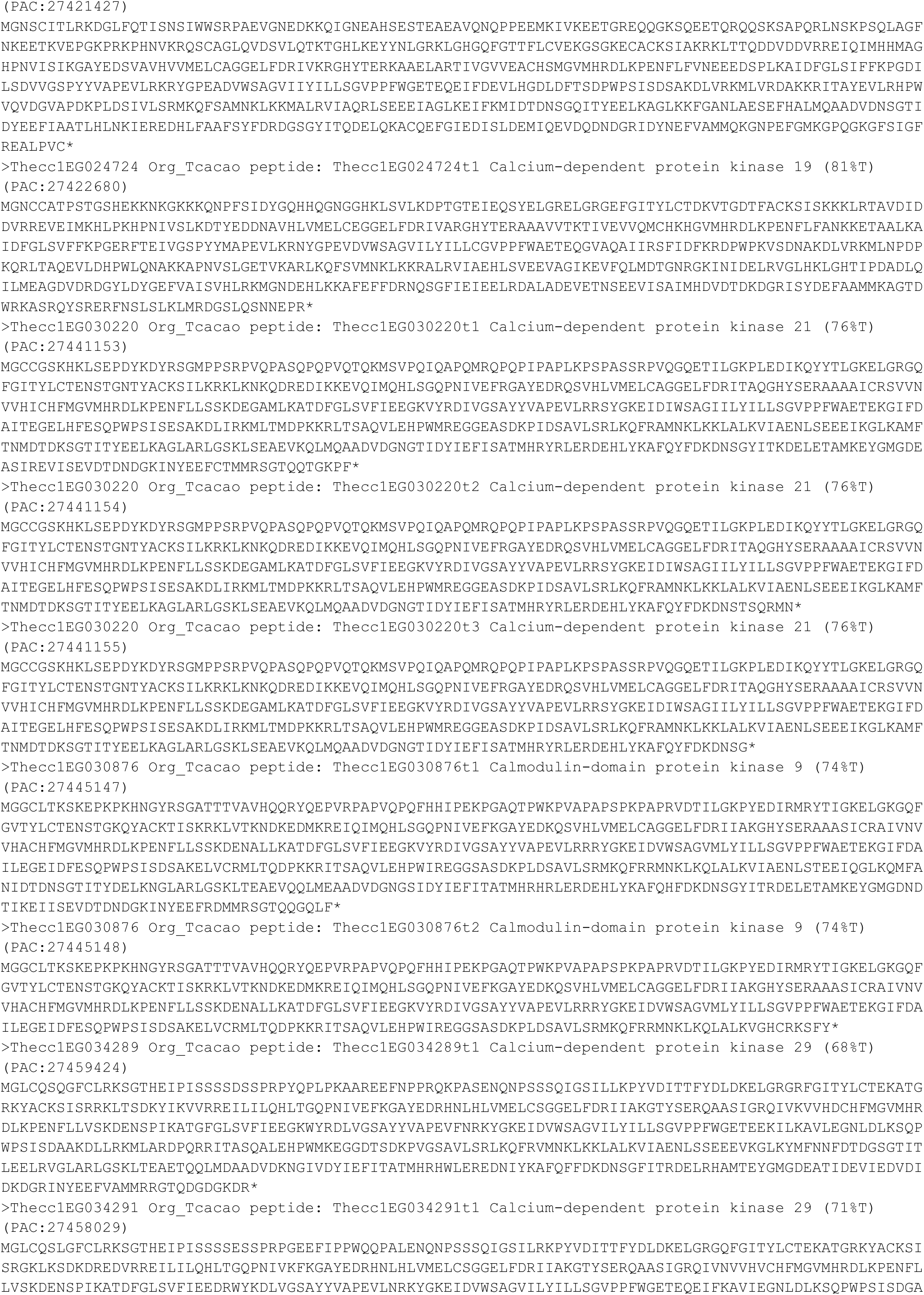
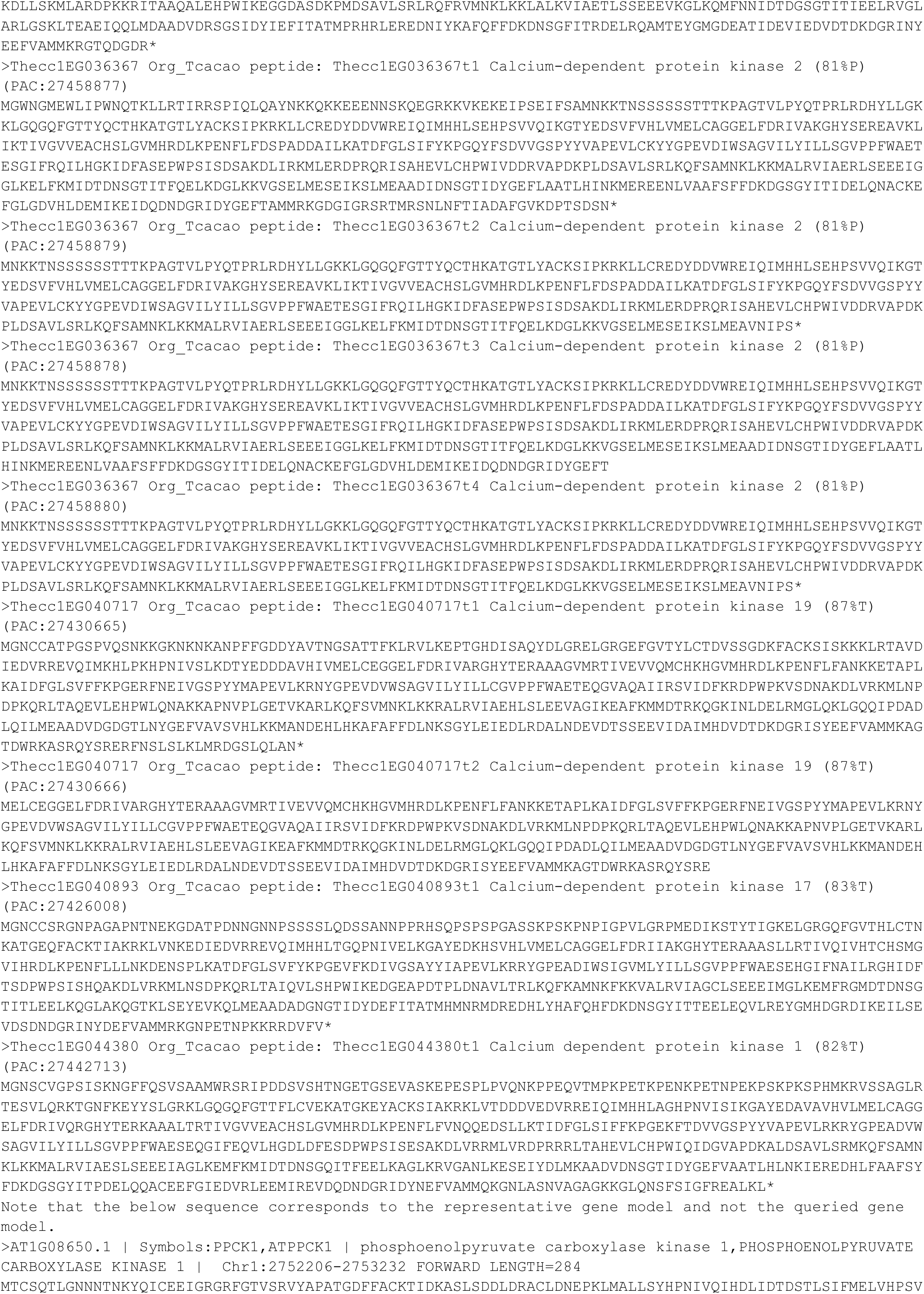
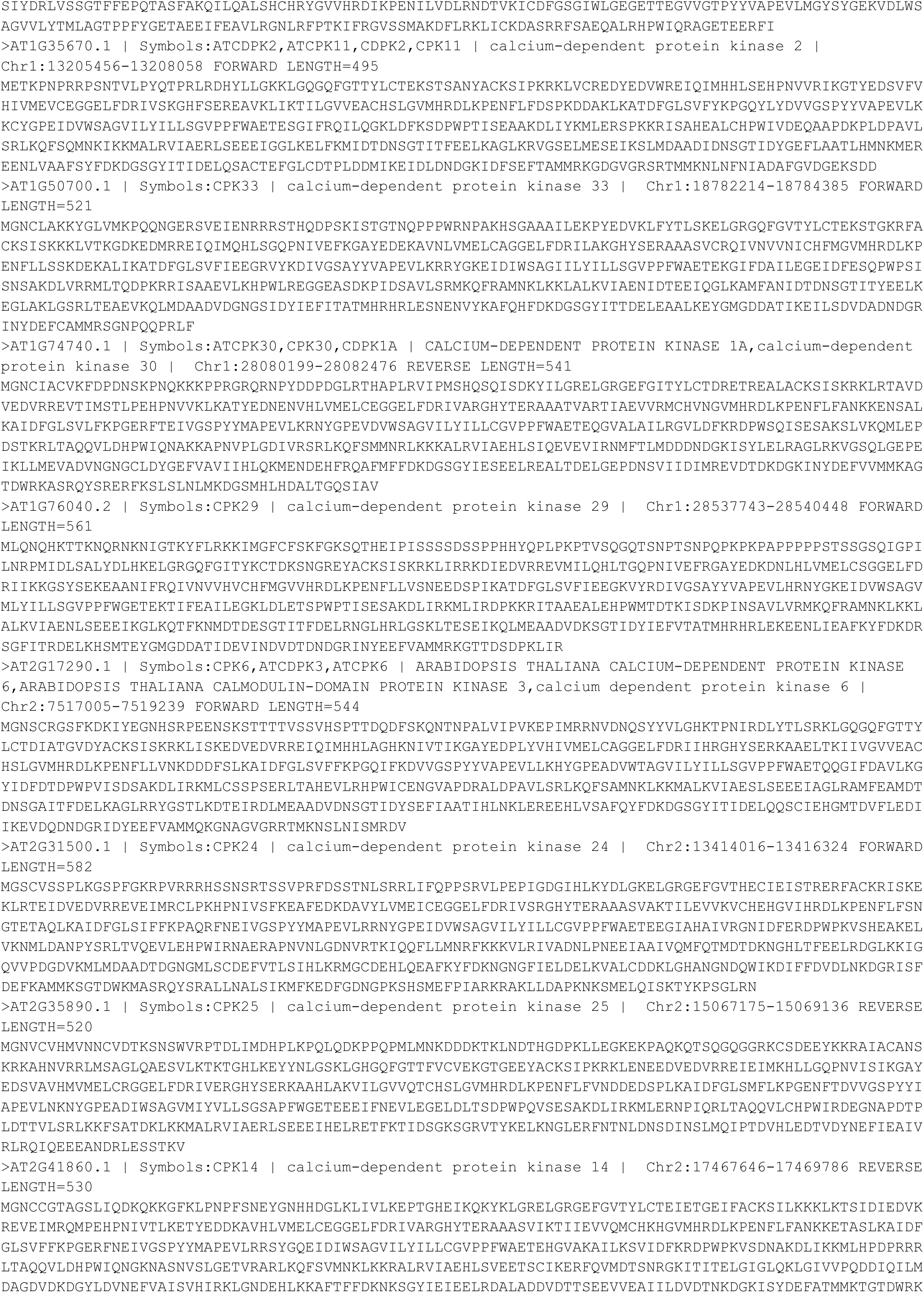
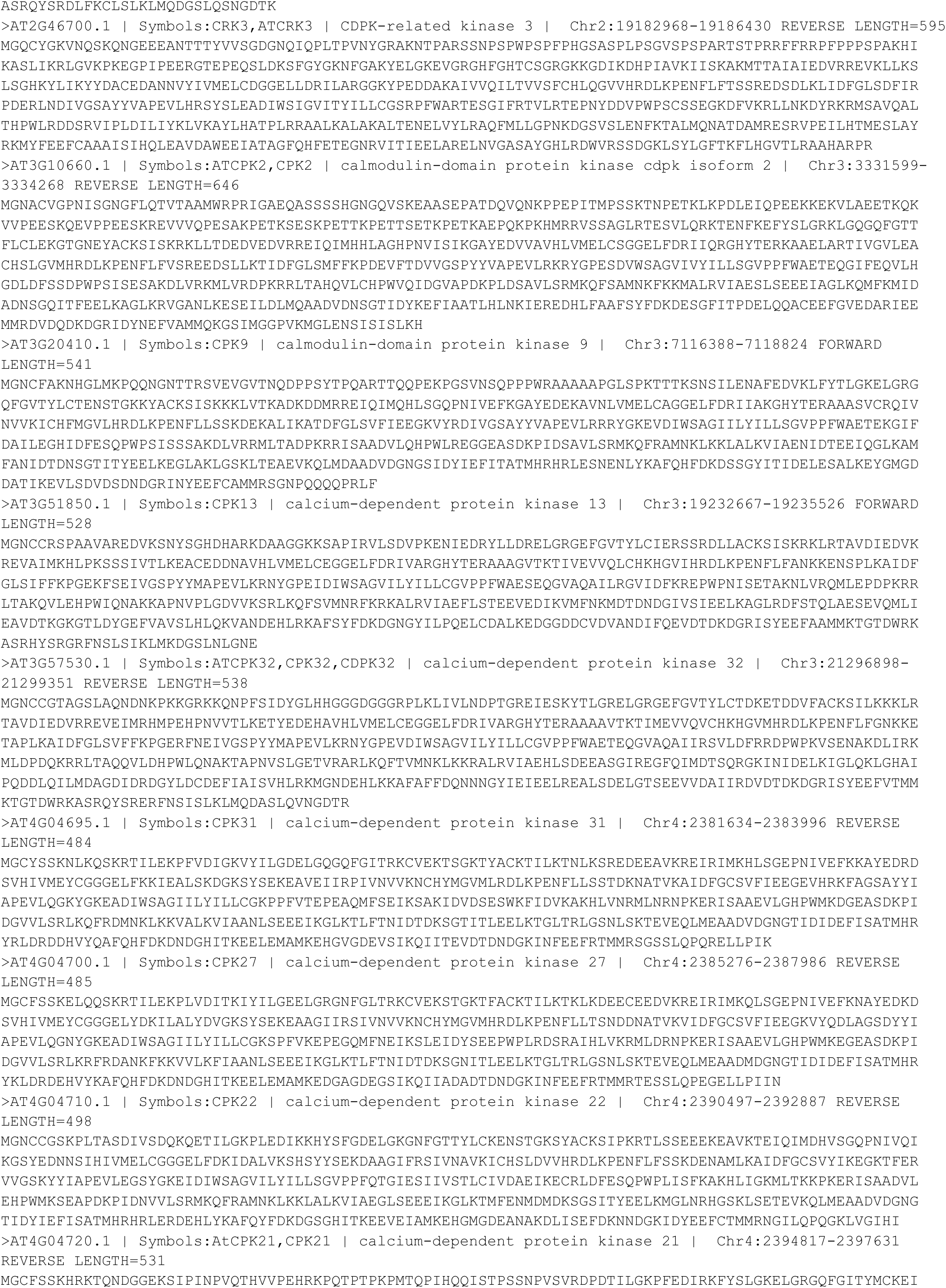
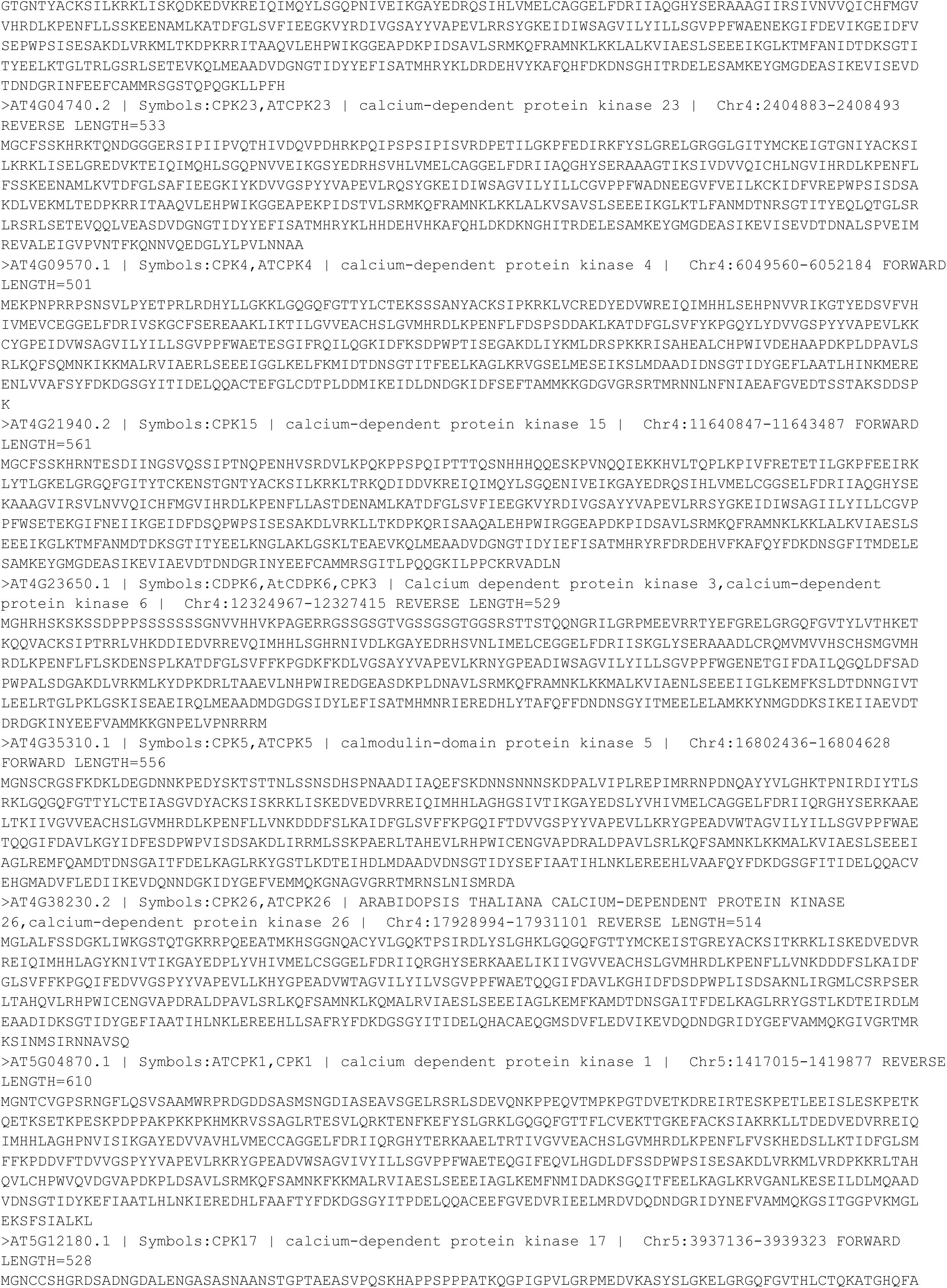
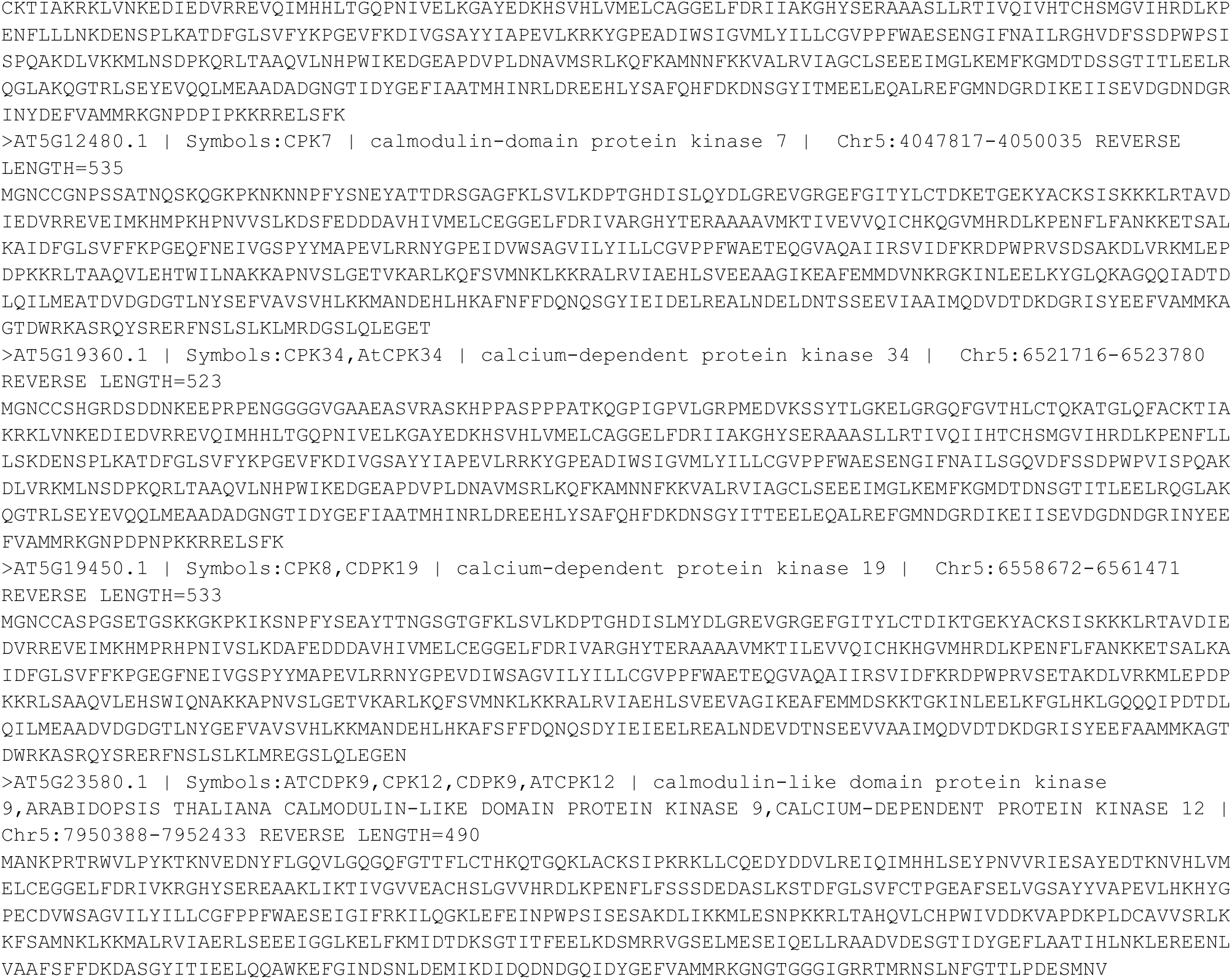

